# The maintenance of polygenic sex determination depends on the dominance of fitness effects which are predictive of the role of sexual antagonism

**DOI:** 10.1101/2020.07.08.193516

**Authors:** Richard P. Meisel

## Abstract

In species with polygenic sex determination, multiple male- and female-determining loci on different proto-sex chromosomes segregate as polymorphisms within populations. The extent to which these polymorphisms are at stable equilibria is not yet resolved. Previous work demonstrated that polygenic sex determination is most likely to be maintained as a stable polymorphism when the proto-sex chromosomes have opposite (sexually antagonistic) fitness effects in males and females. However, these models usually consider polygenic sex determination systems with only two proto-sex chromosomes, or they do not broadly consider the dominance of the alleles under selection. To address these shortcomings, I used forward population genetic simulations to identify selection pressures that can maintain polygenic sex determination under different dominance scenarios in a system with more than two proto-sex chromosomes (modeled after the house fly). I found that overdominant fitness effects of male-determining proto-Y chromosomes are more likely to maintain polygenic sex determination than dominant, recessive, or additive fitness effects. The overdominant fitness effects that maintain polygenic sex determination tend to have proto-Y chromosomes with sexually antagonstic effects (male-beneficial and female-detrimental). In contrast, dominant fitness effects that maintain polygenic sex determination tend to have sexually antagonistic multi-chromosomal genotypes, but the individual proto-sex chromosomes do not have sexually antagonstic effects. These results demonstrate that sexual antagonism can be an emergent property of the multi-chromosome genotype without individual sexually antagonistic chromosomes. My results further illustrate how the dominance of fitness effects has consequences for both the likelihood that polygenic sex determination will be maintained as well as the role sexually antagonistic selection is expected to play in maintaining the polymorphism.

## Introduction

Sex determination is the developmental process by which a genetic signal or environmental cue initiates sexually dimorphic gene regulatory pathways to produce phenotypically different males and females (Bachtrog *et al*. 2014). The master regulators of sex determination evolve fast, often differing between closely related species (Bull 1983; Wilkins 1995; Beukeboom and Perrin 2014). Master regulators can be found on sex chromosomes, and the evolutionary transitions of sex determiners can drive turnover of the sex chromosomes (Abbott *et al*. 2017). These evolutionary transitions include a period of polygenic sex determination (PSD), during which multiple master sex determining loci on different chromosomes segregate as polymorphisms within species (Moore and Roberts 2013). Understanding the population dynamics of PSD is informative of the factors responsible for the evolutionary divergence of sex determination pathways and sex chromosomes.

PSD has been observed in multiple animal and plant species (Moore and Roberts 2013; Beukeboom and Perrin 2014; Bachtrog *et al*. 2014). Most population genetic models predict that PSD will be an unstable intermediate between monogenic equilibria (Rice 1986; van Doorn and Kirkpatrick 2007). Models that do allow for stable PSD often predict that opposing selection pressures in males and females (i.e., sexually antagonistic selection) may be important for maintaining the polymorphism (Bull 1983; Rice 1986; van Doorn and Kirkpatrick 2007; Meisel *et al*. 2016). It is also possible that variable selection pressures across heterogeneous environments could maintain PSD (Bateman and Anholt 2017).

Most previous theoretical treatments of the selection pressures that maintain PSD considered specific cases with a small number of sex chromosomes segregating within a population. For example, in the platyfish, *Xiphophorus maculatus*, a single chromosome can be an X, Y, or Z (Kallman 1973). Orzack et al. (1980) showed that this polymorphism is maintained because heterogametic females (WY or WX) and males (XY) have higher fitness than their homogametic counterparts (XX females or YY males). In addition, van Doorn and Kirkpatrick (2007, 2010) found that sexually antagonistic selection could maintain a polymorphism in which one chromosome segregates as an XY pair and another chromosome is either an XY or ZW pair. However, these models do not capture the full diversity of PSD, which can include more than two XY or ZW pairs segregating within a single species (Moore and Roberts 2013).

Previous models also did not broadly consider how the maintenance of PSD depends on the dominance of fitness effects. The dominance of sexually antagonistic alleles affects their ability to segregate as stable polymorphisms (Kidwell *et al*. 1977; Connallon and Chenoweth 2019), suggesting that dominance may be important for the maintenance of PSD under sexually antagonistic selection. Dominance also differentially affects the evolutionary fate of X-linked and autosomal alleles that experience sex-specific selection pressures (Rice 1984; Charlesworth *et al*. 1987; Fry 2010; Connallon *et al*. 2012). In a species with PSD, there are proto-sex chromosomes that behave neither like autosomes nor like conventional heteromorphic sex chromosomes; proto-sex chromosomes are diploid in both sexes (like autosomes) but have sex-biased modes of inheritance (like conventional sex chromosomes). This is superficially similar to freely recombining pseudoautosomal regions on sex chromosomes (Otto *et al*. 2011), except that in PSD a proto-Y (or proto-W) chromosome can be directly carried by females (males) without X-Y (or Z-W) recombination. Therefore, the expected effects of dominance and sexual antagonism on the evolution of PSD cannot be determined from existing models of autosomal or sex-linked alleles.

The house fly, *Musca domestica*, is a well-suited organism for investigating the maintenance of complex PSD because it has a highly polymorphic sex determination system (Hamm *et al*. 2015). The *M. domestica male determiner* (*Mdmd*) can be found on at least four of the six house fly chromosomes in natural populations (Sharma *et al*. 2017). *Mdmd* causes the house fly ortholog of *transformer* (*Md-tra*) to be spliced into a non-functional isoform, which leads to male-specific splicing and expression of downstream genes in the sex determination pathway, thereby producing fertile males (Hediger *et al*. 2010). In the absence of *Mdmd*, *Md-tra* is spliced into a female-determining transcript that regulates the splicing of downstream targets, which promote the development of female morphological and behavioral traits (Hediger *et al*. 2004; Meier *et al*. 2013).

There are two common male-determining proto-Y chromosomes and one female-determining proto-W chromosome in house fly populations. *Mdmd* is most commonly found on the Y (Y^M^) and third (III^M^) chromosomes in natural populations (Hamm *et al*. 2015). Both Y^M^ and III^M^ are young proto-Y chromosomes that arose recently and are nearly identical in gene content to their homologous proto-X chromosomes, known as X and III (Meisel *et al*. 2017; Son and Meisel 2021). In addition, there is a dominant allele of *Md-tra* that is found in many house fly populations (*Md-tra^D^*), which is resistant to *Mdmd* and causes female development regardless of whether there are copies of *Mdmd* in the genotype (McDonald *et al*. 1978; Hediger *et al*. 2010). *Md-tra* is found on the fourth chromosome, and thus fourth chromosomes carrying *Md-tra^D^* (which I will refer to as IV^F^) are W chromosomes. All sequences of *Md-tra^D^* from around the world are identical (Scott *et al*. 2014), suggesting that *Md-tra^D^* arose recently and IV^F^ is a young proto-W chromosome. In populations where Y^M^, III^M^, and *Md-tra^D^* all segregate, there are 18 possible sex chromosome genotypes (Table 1).

**Table 1.** House fly sex chromosome genotypes.

| genotype |
| --- |

|  | X or Y<sup>M</sup> | III or III<sup>M</sup> | IV or IV<sup>F</sup> | sex |
| --- | --- | --- | --- | --- |
| $f_1$ | X/X | III/III | IV/IV | female |
| $f_2$ | X/X | III/III | IV/IV<sup>F</sup> | female |
| $f_3$ | X/X | III/III<sup>M</sup> | IV/IV<sup>F</sup> | female |
| $f_4$ | X/X | III<sup>M</sup>/III<sup>M</sup> | IV/IV<sup>F</sup> | female |
| $f_5$ | X/Y<sup>M</sup> | III/III | IV/IV<sup>F</sup> | female |
| $f_6$ | X/Y<sup>M</sup> | III/III<sup>M</sup> | IV/IV<sup>F</sup> | female |
| $f_7$ | X/Y<sup>M</sup> | III<sup>M</sup>/III<sup>M</sup> | IV/IV<sup>F</sup> | female |
| $f_8$ | Y<sup>M</sup>/Y<sup>M</sup> | III/III | IV/IV<sup>F</sup> | female |
| $f_9$ | Y<sup>M</sup>/Y<sup>M</sup> | III/III<sup>M</sup> | IV/IV<sup>F</sup> | female |
| $f_{10}$ | Y<sup>M</sup>/Y<sup>M</sup> | III<sup>M</sup>/III<sup>M</sup> | IV/IV<sup>F</sup> | female |
| $m_1$ | X/X | III/III<sup>M</sup> | IV/IV | male |
| $m_2$ | X/X | III<sup>M</sup>/III<sup>M</sup> | IV/IV | male |
| $m_3$ | X/Y<sup>M</sup> | III/III | IV/IV | male |
| $m_4$ | X/Y<sup>M</sup> | III/III<sup>M</sup> | IV/IV | male |
| $m_5$ | X/Y<sup>M</sup> | III<sup>M</sup>/III<sup>M</sup> | IV/IV | male |
| $m_6$ | Y<sup>M</sup>/Y<sup>M</sup> | III/III | IV/IV | male |
| $m_7$ | Y<sup>M</sup>/Y<sup>M</sup> | III/III<sup>M</sup> | IV/IV | male |
| $m_8$ | Y<sup>M</sup>/Y<sup>M</sup> | III<sup>M</sup>/III<sup>M</sup> | IV/IV | male |

There is evidence that natural selection maintains PSD in house fly populations. First, the proto-Y chromosomes have remained at stable frequencies within house fly populations over decades (Kozielska *et al*. 2008; Meisel *et al*. 2016). Second, Y^M^ and III^M^ are distributed along north-south clines on multiple continents (Denholm *et al*. 1986; Tomita and Wada 1989; Hamm *et al*. 2005; Kozielska *et al*. 2008), and temperature is the best predictor of their frequencies (Feldmeyer *et al*. 2008). This suggests that heterogenous (temperature-dependent) selection pressures maintain the Y^M^ and III^M^ chromosomes across populations. The stable frequencies within populations, yet divergent frequencies across populations, suggest that there are specific fitness effects associated with the house fly proto-Y chromosomes within each population. Third, males carrying multiple proto-Y chromosomes (e.g., both Y^M^ and III^M^, or two copies of III^M^) can be found in some populations (e.g., Hamm and Scott 2009). The frequency of *Md-tra^D^* is positively correlated with the frequency of these multi-Y males across populations, suggesting that selection for balanced sex ratios may further maintain PSD (Meisel *et al*. 2016).

The rich body of observational data in house fly makes it a well-suited system around which to develop population genetic models to determine how PSD can be maintained in natural populations. For example, Kozielska et al. (2006) showed that, even in the complex house fly PSD system, sex ratios are not expected to deviate substantially from 1:1 male:female. In addition, despite no experimental evidence for sexually antagonistic effects of any house fly proto-sex chromosomes, another population genetic model demonstrated that the frequencies of the proto-sex chromosomes in natural populations are consistent with sexually antagonistic selection maintaining the polymorphism (Meisel *et al*. 2016). However, that model assumed that the proto-Y chromosomes (Y^M^ and III^M^) have additive fitness effects, which may not be true if, for example, beneficial mutations are recessive or dominant (Orr 2010). In addition, Y chromosomes are expected to carry recessive deleterious alleles (Charlesworth and Charlesworth 2000), which can further violate the assumption of additive fitness effects. To address the shortcomings of previous models, I used population genetic simulations to investigate whether sex chromosomes with non-additive fitness effects can maintain the complex house fly PSD system. I specifically tested if these selection pressures can produce proto-sex chromosome frequencies similar to those observed in natural populations of house fly, and I determined the conditions under which sexually antagonistic effects of those proto-sex chromosomes are expected.

## Materials and Methods

I used a simulation approach (Figure 1A) to identify fitness effects of two proto-Y chromosomes (Y^M^ and III^M^) and one proto-W chromosome (IV^F^) that maintain PSD under four different dominance scenarios. I used simulations to test if fitness effects maintain PSD because the model parameters were too complex to analytically solve for equilibrium conditions. To do so, I first assigned sex-specific fitness effects, *s*_*ij*_, for each *j* proto-Y or proto-W chromosome in each sex *i* by drawing from a random uniform distribution between -1 and 1 (negative values indicate beneficial effects, and positive indicate deleterious). I then used those fitness effects to calculate single chromosome genotype fitness values assuming either additive, dominant, recessive, or overdominant (i.e., heterozygote advantage) fitness effects of the proto-Y chromosomes (Table 2). I calculated the fitness of each of the 18 multi-chromosome genotypes (Table 1) by multiplying all single chromosome fitness values.

**Figure 1.**
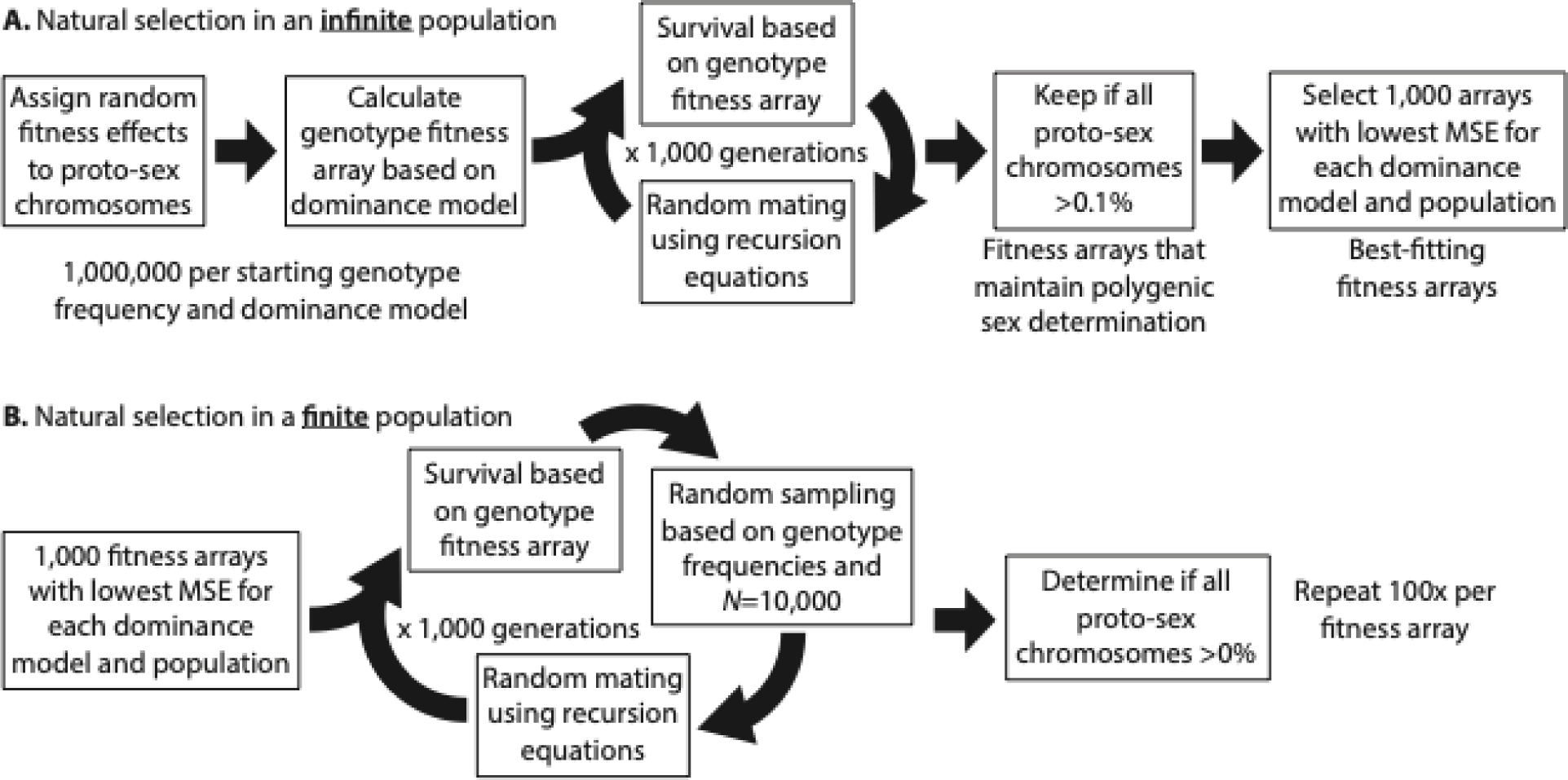
Steps in forward population genetic simulations with **(A)** infinite or **(B)** finite populations. The best-fitting arrays are those with the lowest mean squared error (MSE) comparing simulated proto-sex chromosome frequencies to those observed in each natural population.

**Table 2.** Single proto-Y/proto-X chromosome genotype fitness values with different dominance scenarios.

|  | XX | XY | YY |
| --- | --- | --- | --- |
| *additive* |  |  |  |
| $s_{ij} > 0$ | $1 - s_{ij}$ | $1 - 0.5s_{ij}$ | 1 |
| $s_{ij} < 0$ | 1 | $1 + 0.5s_{ij}$ | $1 + s_{ij}$ |
| *dominant* |  |  |  |
| $s_{ij} > 0$ | $1 - s_{ij}$ | 1 | 1 |
| $s_{ij} < 0$ | 1 | $1 + s_{ij}$ | $1 + s_{ij}$ |
| *recessive* |  |  |  |
| $s_{ij} > 0$ | $1 - s_{ij}$ | $1 - s_{ij}$ | 1 |
| $s_{ij} < 0$ | 1 | 1 | $1 + s_{ij}$ |
| *overdominant (male only)* |  |  |  |
| $s_{ij} > 0$ | $1 - s_{ij}$ | 1 | $1 - s_{ij}$ |
| $s_{ij} < 0$ | $1 + s_{ij}$ | 1 | $1 + s_{ij}$ |

Calculations are shown for determining single chromosome genotype fitness values from the selective effect $s_{ij}$ of proto-Y chromosome $j$ in sex $i$ . Three genotypes are shown (XX, XY, and YY). The Y chromosome could either be $Y^M$ or $III^M$ , and the X chromosome could either be X or III.

To test if the fitness effects result in stable PSD, I determined the frequency of each genotype and proto-sex chromosome after 1,000 generations of random mating with the assigned fitness values (Figure 1A). Simulations were started with either equal frequencies of all 18 genotypes or the genotype frequencies observed in one of three natural populations (CA, NC, or NY) sampled from North America (Hamm and Scott 2008; Scott *et al*. 2013; Meisel *et al*. 2016). I performed simulations for one million different fitness values for each set of starting genotype frequencies and each dominance model using previously developed recursion equations (Hamm 2008; Meisel *et al*. 2016). I retained fitness effects in which all three proto-sex chromosomes remained polymorphic at a frequency >0.1% after 1,000 generations. From those, I selected the 1,000 fitness effects with the smallest mean squared error (MSE) between simulated proto-sex frequencies and those observed in each natural population (CA, NC, or NY) for each dominance model, population, and starting genotype frequency (Figure 1A). In doing so, I am essentially applying an approximate Bayesian computation approach to fit selection pressures to natural populations (Beaumont *et al*. 2002). I observed the same general trends regardless of the starting genotype frequencies, and I only present the results of simulations starting with equal frequencies in the main text (other results are presented in the Supplemental Material). I additionally evaluated how well those fitness effects maintain PSD in a finite population of 10,000 individuals (Figure 1B). I also selected 1,000 random fitness effects for each set of starting genotype frequencies and each dominance model to determine null expectations under my model. Additional details are provided in the Supplemental Methods.

## Results

### Dominance of fitness effects and the maintenance of polygenic sex determination

I find that the probability that PSD is maintained (with a frequency of each proto-sex chromosome >0.1% after 1,000 generations) is affected by the dominance of the fitness effects of the Y^M^ and III^M^ proto-Y chromosomes (Supplemental Figures S1A, S2A). When fitness effects of proto-Y chromosomes are additive, <5% of fitness values maintain PSD. In contrast, when fitness effects of the proto-Y chromosomes are recessive, ∼20% of fitness values maintain PSD. Overdominant fitness effects are the most likely to maintain PSD, with >40% of fitness values maintaining proto-sex chromosomes at a frequency >0.1% after 1,000 generations. When additive, recessive, or overdominant (but not dominant) fitness effects maintain PSD, there is a broad range of frequencies at which the proto-Y chromosomes (Y^M^ and III^M^) can segregate as polymorphisms (Figure 2; Supplemental Figure S3). Randomly chosen fitness effects, in comparison, tend to produce one of three proto-Y chromosome frequency classes: 0 (loss of a proto-Y, because a different XY system has taken over), 1 (fixation of the proto-Y chromosomes, corresponding to a monogenic ZW system), and 0.25 (the expectation for a monogenic XY system).

**Figure 2.**
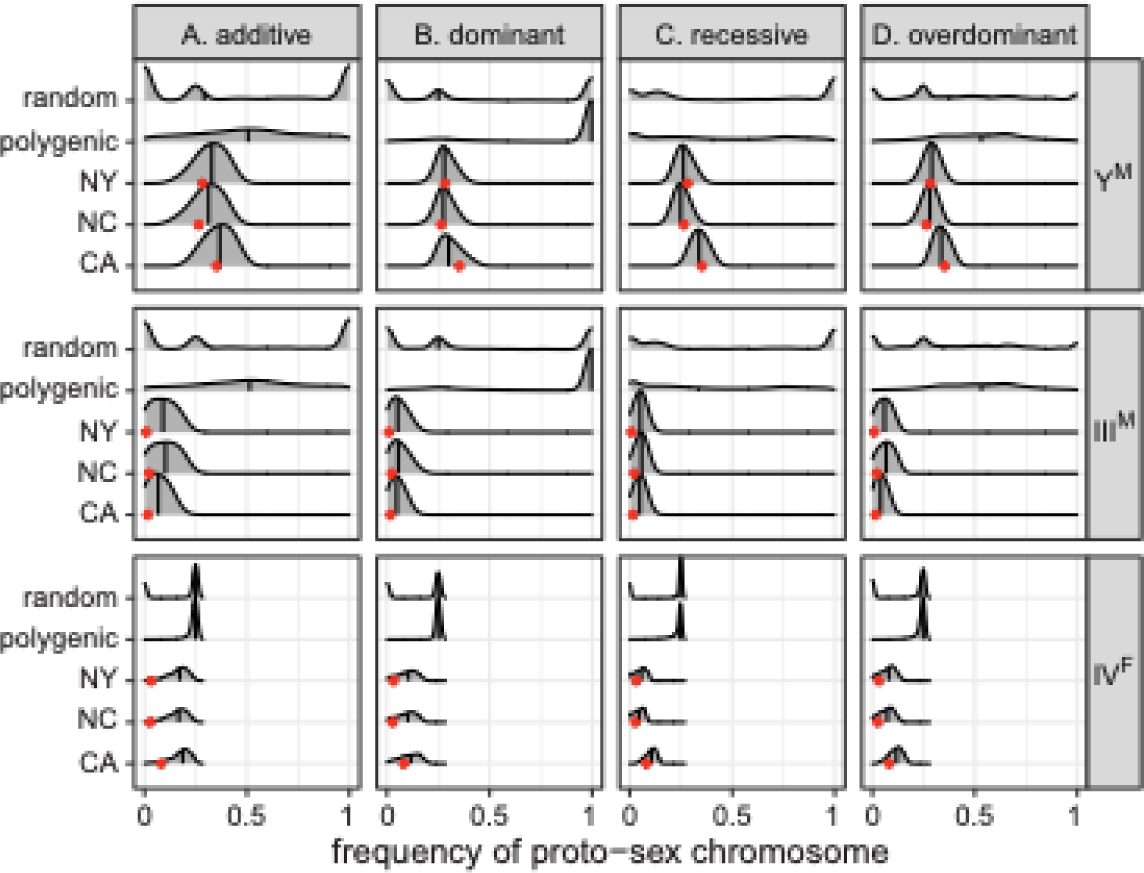
Smoothed histograms show the frequency of each proto-sex chromosome (Y^M^, III^M^, and IV^F^) after 1,000 generations in simulations using either 1,000 random fitness arrays (random), fitness arrays that maintain PSD (polygenic), or the 1,000 best-fitting fitness arrays for each population (CA, NC, or NY). Simulations were started with equal frequencies of all genotypes. The vertical line within each histogram shows the median. Red dots show the observed proto-sex chromosome frequencies in each natural population. Fitness arrays were calculated assuming either **(A)** additive, **(B)** dominant, **(C)** recessive, or **(D)** overdominant fitness effects of the proto-Y chromosomes.

Most dominant fitness effects that appear to maintain PSD are paths to the eventual fixation of the proto-Y chromosomes. More than 25% of dominant fitness maintain PSD for 1,000 generations (Supplemental Figures S1A, S2A), but most of those produce very high frequencies of the Y^M^ and III^M^ chromosomes (Figure 2B; Supplemental Figure S3). A proto-Y chromosome with a dominant fitness effect segregating at very high frequency is consistent with a beneficial effect that masks the recessive variant (in this case the homologous proto-X), allowing the proto-X chromosome to persist at a low frequency for a long time (Hartl and Clark 2007). To test this prediction, I selected fitness effects that maintain PSD for 1,000 generations, and I ran simulations with those fitness effects for 1,000,000 generations. I considered a proto-Y chromosome to reach “fixation” after 1,000,000 generations if it achieves a frequency >99.9% (i.e., the same threshold for maintaining PSD after 1,000 generations). Consistent with the prediction, >50% of dominant fitness effects that maintain PSD for 1,000 generations eventually cause the Y^M^ or III^M^ proto-Y chromosomes to reach fixation within 1,000,000 generations, regardless of the starting genotype frequencies (Supplemental Figures S4–S7). In contrast, ≤1% of additive, recessive, or overdominant fitness effects of proto-Y chromosomes that maintain PSD for 1,000 generations eventually result in fixation of a proto-Y chromosome after 1,000,000 generations. Despite the eventual fixation of proto-Y chromosomes with many dominant fitness effects, there are a subset of dominant fitness effects that maintain Y^M^ and III^M^ at lower frequencies, resembling those observed in natural populations (Figure 2B). For these dominant fitness effects that produce proto-sex chromosome frequencies similar natural populations, fixation of proto-Y chromosomes never occurs after 1,000,000 generations (Supplemental Figures S4–S7). Therefore, dominant fitness effects can maintain PSD at equilibrium, although most dominant fitness effects result in fixation or loss of the proto-Y chromosomes. Below, I describe how dominant fitness effects that can or cannot maintain PSD differ in their signatures of sexual antagonism.

### Evaluating if selection pressures can produce proto-sex chromosome frequencies observed in natural populations

Selecting only the 1,000 fitness effects that produce proto-sex chromosome frequencies most similar to natural populations further affects the relationships between dominance and proto-sex chromosome frequencies. For example, dominant, recessive, and overdominant fitness effects can produce proto-sex chromosome frequencies that are more similar to those in natural populations when compared to additive fitness effects (Figure 2; Supplemental Figures S1, S2, S3). The poorer fit of additive models arises because they struggle to produce frequencies of III^M^ and IV^F^ as low as those observed in the CA, NC, and NY populations (Figure 2; Supplemental Figure S3).

The low frequency of the III^M^ chromosome in the CA, NC, and NY populations has consequences for the ability of natural selection to maintain the polymorphism. As above, I define loss of the III^M^ chromosome (or fixation of the III chromosome) as a frequency below 0.1% (or above 99.9% for III). Additive fitness effects that maintain the III^M^ chromosome at frequencies similar to those in CA, NC, or NY for 1,000 generations often (9–17% of the time) result in the loss of III^M^ within 1,000,000 generations (Supplemental Figures S8, S9). In contrast, <7% of dominant, recessive, or overdominant fitness effects that maintain III^M^ at frequencies observed in natural populations for 1,000 generations lead to the eventual loss of III^M^ within 1,000,000 generations. III^M^ is especially likely to be lost within 1,000,000 generations when it is segregating at a lower frequency after 1,000 generations, regardless of the dominance of fitness effects (Supplemental Figures S8, S9). The Y^M^ chromosome, in comparison, segregates at a higher frequency that III^M^ all three populations (Figure 2B). Y^M^ is lost in ≤2% of simulations after 1,000,000 generations when I apply fitness effects that maintain proto-sex chromosomes at frequencies similar to those observed in natural populations, regardless of the dominance of fitness effects (Supplemental Figure S10). I therefore conclude that low frequency proto-Y chromosomes, such as III^M^, are difficult to maintain as polymorphisms under the model I have applied here. Moderate frequency proto-Y chromosomes, such as Y^M^, on the other hand, can be maintained under a variety of dominance scenarios.

The majority of fitness effects that maintain PSD cause IV^F^ to segregate at a frequency of 25%, regardless of the dominance of the proto-Y chromosomes (Figure 2; Supplemental Figure S3). A frequency of 25% is the expectation for a W chromosome in a randomly mating population with monogenic ZW sex determination. Therefore, in a system with 2 proto-Y chromosomes and one proto-W, PSD is expected to be maintained by the (near) fixation of the ZW female genotype. In contrast, IV^F^ makes up 2–9% of all fourth chromosomes in the CA, NC, and NY populations (Hamm and Scott 2008; Scott *et al*. 2013; Meisel *et al*. 2016). A subset of fitness arrays in my simulations maintain IV^F^ at <25% for all dominance models, including those that produce proto-sex chromosome frequencies most similar to those observed in natural populations (Figure 2; Supplemental Figure S3). It is thus possible for PSD to be maintained with a single proto-W chromosome at a frequency <25%. However, the frequency of IV^F^ is still greater in simulated populations than in natural populations (Figure 2).

### Selection pressures can maintain polygenic sex determination in finite populations

I next used simulations to examine how well natural selection maintains PSD when population sizes are finite (*N*=10^4^ individuals). I calculated the proportion of simulations (out of 100) in which a given proto-sex chromosome reached fixation in a finite population for each fitness array to estimate of the probability of fixation (*P_fix_*). The *P_fix_* value for a given proto-Y or proto-W chromosome (Y^M^, III^M^, or IV^F^) is equal to the probability of loss (*P_loss_*) of the homologous proto-X or proto-Z (X, III, or IV), and *vice versa*. I also simulated finite populations of the same size but without selection (i.e., genetic drift only). I used Fisher’s exact test to determine if the *P_fix_* or *P_loss_* of a proto-sex chromosome is significantly different between populations with and without selection.

When a proto-sex chromosome is at low frequency, selection pressures that maintain PSD decrease *P_fix_* and *P_loss_* in finite populations, relative to populations without selection.

For example, III^M^ is rare in all three populations, and it is the most likely proto-sex chromosome to be lost when population size is finite (Supplemental Figures S11, S12). In turn, selection pressures that maintain PSD, decrease *P_fix_* of chromosome III relative to drift alone (Figure 3; Supplemental Figure S13), which is equivalent to decreasing *P_loss_*of III^M^. The IV^F^ proto-W chromosome is also at a low frequency in all three populations, and the maintenance of the IV^F^ polymorphism (i.e., the standard IV chromosome does not fix) is substantially greater when there is selection (Figure 3; Supplemental Figure S13). Therefore, when loss via drift is likely because a proto-sex chromosome is at low frequency (such as III^M^ and IV^F^), selection pressures that maintain PSD decrease the probability of loss.

**Figure 3.**
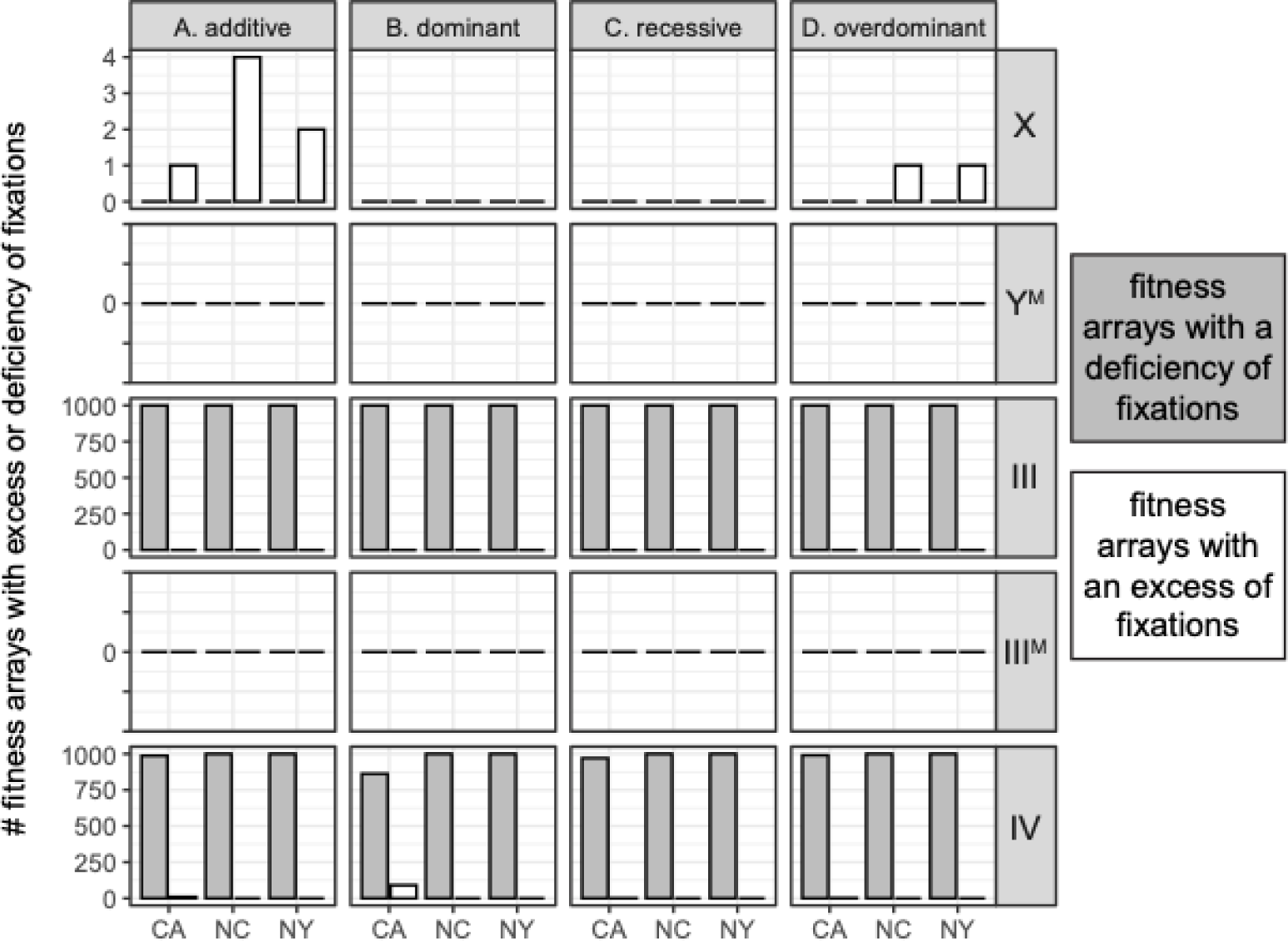
Bar plots show the number of fitness arrays that result in a deficiency (gray) or excess (white) of fixations of each proto-sex chromosome (X, Y^M^, III, III^M^, or IV) in a finite population, relative to simulations in which there are no fitness differences across proto-sex chromosomes (i.e., drift only). The fitness arrays are the 1,000 best-fitting fitness values for each population (CA, NC, or NY) when simulations were started with equal frequencies of all genotypes. Fitness effects of the proto-Y chromosomes are either **(A)** additive, **(B)** dominant, **(C)** recessive, or **(D)** overdominant.

A different effect is observed for sex chromosomes segregating at higher frequencies. The X and Y^M^ chromosomes rarely fix in finite populations with selection (Supplemental Figures S11, S12). Y^M^ is at a higher frequency than III^M^ and IV^F^ across all three populations (Meisel *et al*. ^2^016^)^, and the high frequency of Y^M^ is likely responsible for its low *P_loss_* within 1,000 generations. However, for a small fraction of fitness effects that maintain proto-sex chromosomes at frequencies similar to those in natural populations (<10%), *P_fix_* for both the X and Y^M^ chromosomes is greater with selection and drift than under drift alone (Figure 3). The slightly elevated *P_fix_* of the X and Y^M^ chromosomes with selection is observed for all fourdominance models (Supplemental Figure S13). Therefore, when fixation or loss is unlikely via drift (because the proto-Y chromosome is at high frequency), selection pressures that maintain PSD do not decrease *P_fix_* or *P_loss_* of a proto-sex chromosome in a finite population.

### Sexual antagonism and the maintenance of polygenic sex determination

Sexually antagonistic selection is predicted to maintain PSD (Bull 1983; Rice 1986; van Doorn and Kirkpatrick 2007; Meisel *et al*. 2016). A negative intersexual fitness correlation is a hallmark of sexually antagonistic genetic variation (Rice and Chippindale 2001; Chippindale *et al*. 2001; Bonduriansky and Chenoweth 2009). I therefore inspected if there is a negative correlation between male and female fitness when PSD is maintained. Each of the eight male house fly proto-sex chromosome genotypes has a corresponding female genotype that differs only because females carry a copy of the IV^F^ proto-W chromosome and males do not (Table 1). There are two additional female genotypes (X/X; III/III; IV/IV and X/X; III/III; IV^F^/IV) that do not have a corresponding male genotype because they do not have a proto-Y chromosome. To test for sexually antagonistic multi-chromosome genotypes, I calculated Spearman’s rank order correlation of fitness between males and females (*ρMF*) for the eight pairs of male and female genotypes in each fitness array.

Additive fitness effects that maintain PSD (i.e., all proto-sex chromosomes at frequency>0.1%) tend to have *ρMF<0* (Figure 4A; Supplemental Figure S14). However, when proto-sex chromosome frequencies resemble those in natural populations and fitness effects are additive, *ρMF* can be either positive or negative. There is also no consistent signal of *ρMF<0* for random fitness effects, regardless of dominance (Figure 4; Supplemental Figure S14), demonstrating that *ρMF<0* is not an intrinsic property of the model. In comparison,*ρMF<0* when I previously applied a model with additive fitness effects both within and across chromosomes (Meisel *et al*. 2016). In the model presented here, multi-chromosome genotype fitness is calculated as the product of single chromosome genotype fitness values, rather than by summing across chromosomes. Therefore, there is some evidence that additive fitness effects that maintain PSD result in *ρMF<0*, but it depends on specifics of how the model is parameterized. I address the causes and effects of *ρMF<0* in more detail in the following section.

**Figure 4.**
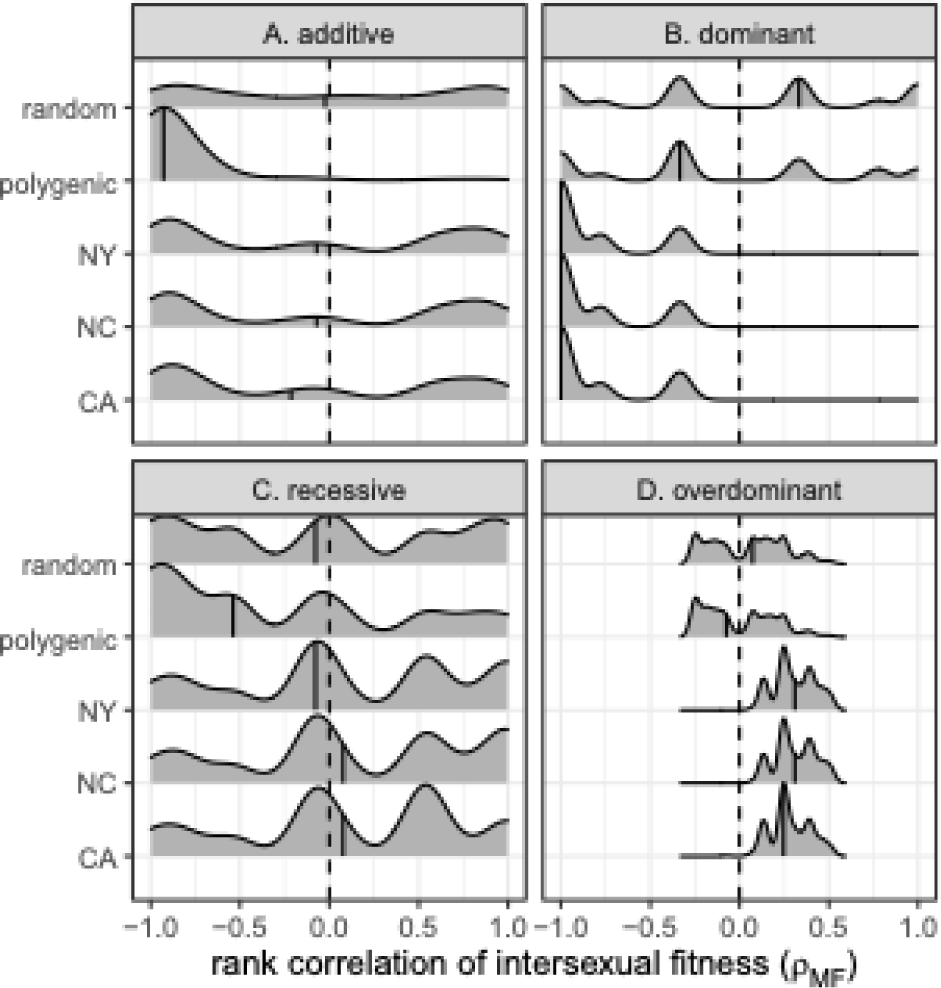
Smoothed histograms show distributions of intersexual fitness correlations (*ρMF*) for genotype fitness arrays used in simulations started with equal frequencies of all genotypes. Correlations are shown for the 1,000 best-fitting genotypic fitness arrays for the CA, NC, and NY populations. Correlations are also shown for all fitness arrays that maintain PSD (polygenic), as well as 1,000 random fitness arrays. Dashed vertical lines show a correlation of 0, and solid vertical lines within histograms show the median.

Intersexual fitness correlations tend to be negative (*ρMF<0*) when fitness effects are dominant and proto-sex chromosome frequencies resemble those in natural populations (Figure 4B; Supplemental Figure S14B). In contrast, *ρMF* can be positive or negative for dominant fitness effects that maintain PSD regardless of the proto-sex chromosome frequency. However, as noted above, many of the cases where dominant fitness effects appear to maintain PSD are instead on a path to fixation that has not yet been reached at 1,000 generations. When I only sample the dominant fitness effects that maintain PSD for 1,000,000 generations, I find that *ρMF<0* (Supplemental Figure S15). Therefore, assuming 1,000,000 generations adequately approximates the equilibrium state, *ρMF<0* when dominant fitness effects maintain PSD.

When fitness effects are overdominant or recessive, *ρMF* does not provide evidence for sexually antagonistic selection. First, there is no clear pattern of *ρMF<0* when fitness effects are recessive (Figure 4C; Supplemental Figure S14C). Second, overdominant fitness effects that produce the best-fitting genotype fitness arrays nearly all have *ρMF>0* (Figure 4D; Supplemental Figure S14D), but this result is difficult to interpret because overdominant effects produce a concave relationship between the number of copies of a proto-Y chromosome and male fitness (i.e., heterozygous males are defined as the most fit). Therefore, a simple rank order correlation (i.e., *ρMF*) does not adequately capture the relationship between male and female fitness when the proto-Y chromosomes have overdominant effects in males.

### Negative intersexual fitness correlations are not caused by sexually antagonistic proto-Y chromosomes

The negative *ρMF* that I observe for additive and dominant fitness effects that maintain PSD (Figure 4; Supplemental Figure S14) are suggestive of sexually antagonistic selection. A potential cause of *ρMF<0* is sexually antagonistic effects of the proto-Y chromosomes if, for example, Y^M^ and III^M^ carry male-beneficial and female-detrimental alleles (Fisher 1931; Rice 1987). In contrast to this hypothesis, I do not observe a consistent signal of male-beneficial and female-detrimental proto-Y chromosomes when fitness effects are additive or dominant (Figure 5; Supplemental Figure S16).

**Figure 5.**
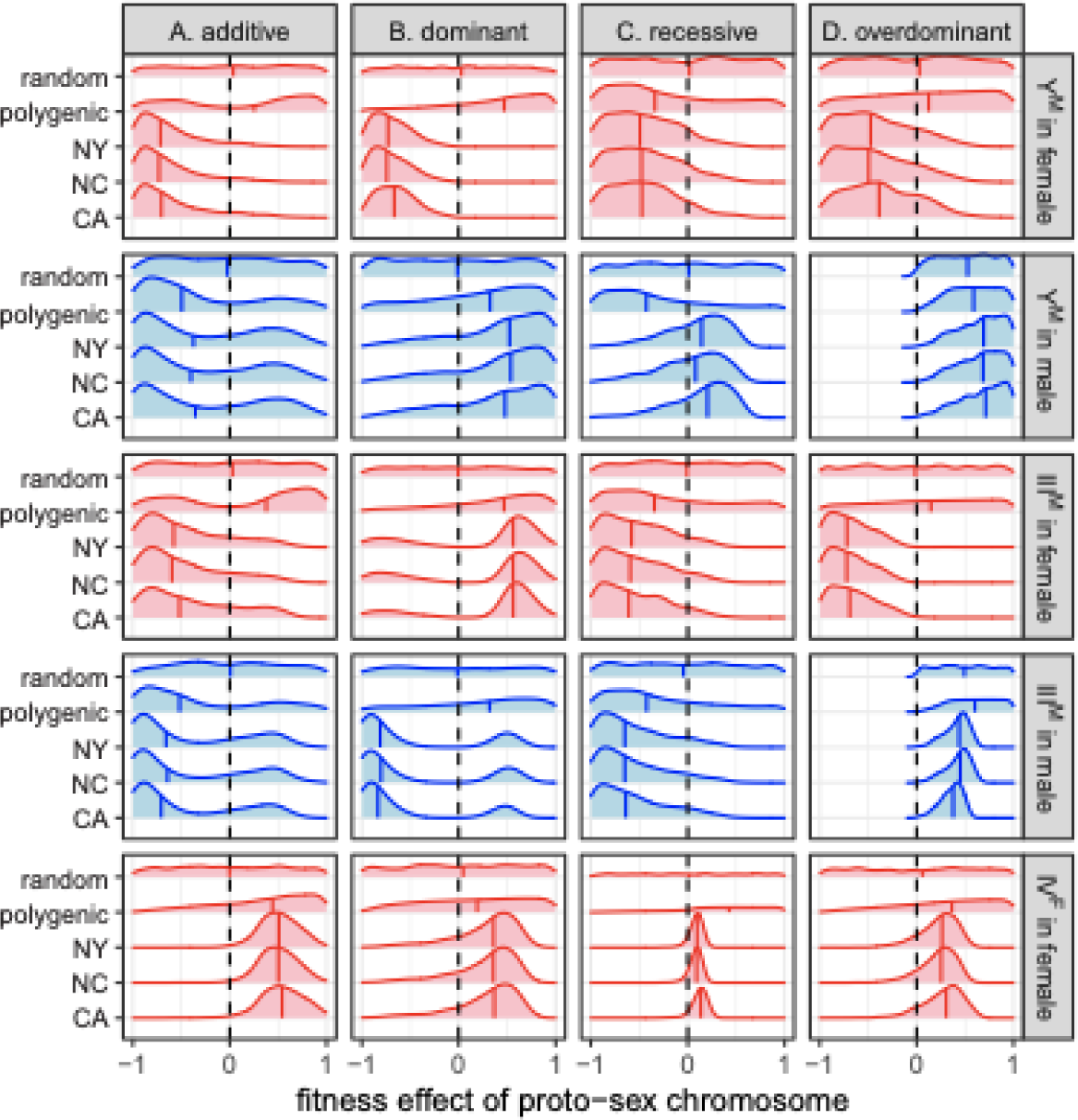
Smoothed histograms show the distributions of fitness effects of each proto-sex chromosome in each sex for 1,000 random fitness arrays (random), fitness arrays that maintain PSD (polygenic), or the 1,000 best-fitting fitness arrays for each population (CA, NC, or NY). Simulations were started with equal frequencies of all genotypes. The vertical line within each histogram shows the median, and dashed lines in each panel are at fitness value of 0. Fitness arrays were calculated assuming either (**A)** additive, **(B)** dominant, **(C)** recessive, or **(D)** overdominant fitness effects of the proto-Y chromosomes.

There is some evidence that Y^M^ is male-beneficial and female-detrimental when dominant fitness effects maintain PSD (Figure 5B; Supplemental Figure S16B). Additionally, both Y^M^ and III^M^ often confer a greater fitness benefit to males than females when dominant fitness effects maintain PSD (Supplemental Figures S17, S18, S19). However, III^M^ does not tend to have male-beneficial and female-detrimental effects in the models with dominant fitness effects (Figure 5B; Supplemental Figure S16B). Therefore, when dominant fitness effects maintain PSD, the multi-chromosome genotypes tend to have sexually antagonistic fitness effects (Figure 4; Supplemental Figures S14, S15), but the individual chromosomes are not necessarily sexually antagonistic themselves.

In contrast to most expectations, additive fitness effects that maintain PSD often have female-beneficial and male-detrimental effects of the proto-Y chromosomes (Figure 5A; Supplemental Figure S16A). In addition, the proto-Y chromosomes do not consistently confer a greater fitness benefit to males than females when PSD is maintained by additive fitness effects (Supplemental Figures S17, S18). Female-beneficial and male-detrimental Y chromosomes are opposite of the expected effects under most models of sex chromosome evolution (Rice 1987, 1996). However, some previous population genetic modeling has suggested the possibility of “feminization” of the Y chromosome (Cavoto *et al*. 2018).

Overdominant fitness effects that maintain proto-sex chromosomes at frequencies similar to those observed in natural populations have the strongest evidence of male-beneficial and female-detrimental proto-Y chromosomes (Figure 5D; Supplemental Figures S16D, S17D, S18D). The overdominant model is constrained to require male-beneficial effects of the proto-Y chromosomes in heterozygotes, but female fitness effects can be beneficial or deleterious (see Supplemental Methods). Despite the full range of possible female fitness effects in the overdominant model, when proto-sex chromosomes are at frequencies similar to those in the CA, NC, or NY populations, the proto-Y chromosomes are almost always female-deterious (Figure 5D; Supplemental Figures S16D). These sexually antagonistic (female-deleterious and male-beneficial) effects of the Y^M^ and III^M^ proto-Y chromosomes in the overdominant model does not cause negative intersexual correlations of multi-chromosomal genotype fitness values (Figure 4; Supplemental Figure S14) because heterozygous males are defined as the most fit.

### Female-beneficial effects of the proto-W chromosome

The IV^F^ proto-W chromosome is nearly always beneficial to females when PSD is maintained. This is the case regardless of the dominance of fitness effects of the proto-Y chromosomes, whether or not proto-sex chromosomes frequencies are similar to those in natural populations, and independently of the genotype frequencies used to start the simulations (Figure 5; Supplemental Figure S16). There also tends to be a positive correlation between the fitness effect and frequency of IV^F^ when PSD is maintained (Supplemental Figures S20, S21). A positive correlation indicates that the more female-beneficial the IV^F^ chromosome, the higher its frequency. The only exception to this rule is when dominant fitness effects maintain PSD for 1,000 generations; however, as described above, these cases are on a path to fixation and therefore not at equilibrium

## Discussion

### Dominance of fitness effects, sexually antagonistic sex chromosomes, and sexually antagonistic genotypes

I used multiple approaches to test if there is evidence of sexually antagonistic selection when PSD is maintained in my simulations. Sexual antagonism has previously been identified as a key component of the maintenance of PSD within populations (Bull 1983; Rice 1986; van Doorn and Kirkpatrick 2007; Meisel *et al*. 2016). A negative inter-sexual correlation of fitness across genotypes is a hallmark of sexually antagonistic selection (Rice and Chippindale 2001; Chippindale *et al*. 2001; Bonduriansky and Chenoweth 2009). I find strong evidence for*ρMF<0* when dominant fitness effects maintain PSD (Figure 4B; Supplemental Figure S15).

There is also some evidence that additive fitness effects that maintain PSD tend to have negative *ρMF*, although it is dependent on the frequency at which the proto-sex chromosomes are maintained (Figure 4A; Supplemental Figure S14A). However, neither dominant nor additive fitness effects that maintain PSD have male-beneficial and female-detrimental effects of all individual proto-Y chromosomes (Figure 5; Supplemental Figure S16). Therefore, sexually antagonistic multi-chromosomal genotypes that maintain PSD can be emergent properties of cumulative fitness effects across chromosomes, without each chromosome having the same sexually antagonistic effects. This is evidence for a decoupling between single chromosome sexually antagonistic effects and sexual antagonism across multi-chromosome genotypes. It also suggests that testing for sexually antagonistic effects of individual proto-sex chromosomes may not be a sufficient assay of the role sexual antagonism plays in the maintenance of complex, multi-chromosomal PSD.

It has generally been observed that the dominance of fitness effects is an important parameter affecting the ability sexually antagonistic selection to maintain genetic variation (Connallon and Chenoweth 2019). For example, a sexually antagonistic allele is more likely to be maintained as a polymorphism if the sex-specific deleterious effects are recessive (Kidwell *et al*. 1977). In contrast, when fitness effects are dominant, stronger selection pressures are required to maintain the polymorphism (Kidwell *et al*. 1977). My work presented here contributes to the special case where the alleles under selection are on proto-sex chromosomes (as opposed to autosomes or heteromorphic sex chromosomes). Consistent with Kidwell *et al*. (1977), I find that dominant fitness effects that maintain PSD do indeed have strong selection in favor of or against proto-Y chromosomes in one sex (Figure 5; Supplemental Figure S16).

In most of the models I consider here, the dominance of fitness effects of the proto-Y chromosomes is the same in both sexes. However, the dominance of fitness effects may be sex-specific (Barson *et al*. 2015; Grieshop and Arnqvist 2018). Such dominance reversals could be important for the maintenance of sexually antagonistic alleles (Spencer and Priest 2016; Connallon and Chenoweth 2019), including at sex-linked loci (Fry 2010). Therefore, it is possible that sex-specific dominance could create a broader range of fitness values than I observe to maintain PSD via sexually antagonistic selection. Consistent with this prediction, I observed PSD maintained most frequently when fitness effects of proto-Y chromosomes are overdominant in males and additive in females (see below for more discussion of overdominant fitness effects). Future work should further examine how sex-specific dominance affects the maintenance of PSD.

Sexually antagonistic proto-Y chromosomes require there to be segregating sexually antagonistic genetic variation. Male-beneficial and female-deleterious alleles are likely to accumulate on Y chromosomes because of their male-limited inheritance (Rice 1984). It is not clear, however, if that prediction applies when the Y chromosome can be transmitted through females in a complex PSD system. A more appropriate point of comparison for PSD systems may be autosomal and X-linked sexually antagonistic genetic variation, of which there is evidence for a substantial amount (Rice 1992; Chippindale *et al*. 2001; Calsbeek and Sinervo 2004; Fedorka and Mousseau 2004; Foerster *et al*. 2007; Brommer *et al*. 2007; Innocenti and Morrow 2010). The fact that such variation exists in a broad range of taxa suggests that sexually antagonistic proto-Y chromosomes may be possible in a PSD system.

My results are consistent with theory and data that suggest sexually antagonistic selection is important for the early evolution of sex chromosomes. For example, van Doorn and Kirkpatrick (2007, 2010) showed that selection on linked sexually antagonistic alleles can favor a new sex determining locus because sex-limited inheritance can resolve the sexual conflict. Empirical studies have also identified sexually antagonistic variants on young Y and W chromosomes (Lindholm and Breden 2002; Roberts *et al*. 2009). In addition, once a new sex chromosome is established, fixation of additional alleles with sex-specific beneficial effects can be favored on the sex-specific Y or W chromosome (Abbott *et al*. 2017). It remains to be determined, however, whether young sex chromosomes are enriched for sexually antagonistic variants, if those variants are maintained by balancing selection, and if the sexually antagonistic alleles contribute to the maintenance of PSD. Investigating these problems could follow the approaches previously developed to test if balancing selection maintains sexually antagonistic alleles throughout the genome (e.g., Dutoit *et al*. 2018; Ruzicka *et al*. 2019).

### Overdominance and the maintenance of polygenic sex determination

Overdominant fitness effects of proto-Y chromosomes in males are more likely to maintain PSD than any other dominance scenario that I tested. Nearly half of the 4,000,000 overdominant fitness effects that I tested maintain all three proto-sex chromosomes at a frequency >0.1% for at least 1,000 generations (Supplemental Figures S1A, S2A). In addition, nearly all of those overdominant fitness effects that maintain PSD for 1,000 generations continue to do so for at least 1,000,000 generations (Supplemental Figures S4–S7), suggesting they are indeed stable polymorphisms at equilibrium. The frequencies with which overdominant fitness effects can maintain multiple polymorphic proto-sex chromosomes covers the full range of possible values, from close to 0 to nearly 1 (Figure 2D; Supplemental Figure S3D).

The extent of overdominant genetic variation—and balancing selection more generally—has been a topic of considerable debate in population genetics (Crow 1987; Delph and Kelly 2014; Fijarczyk and Babik 2015), with a few classic examples of heterozygote advantage (e.g., Dobzhansky 1947; Allison 1956). Contemporary evidence for overdominant fitness effects in natural populations is mixed, including some work that suggests overdominance is rare (Andrés *et al*. 2009; Sellis *et al*. 2011, 2016; Hedrick 2012; Bitarello *et al*. 2018). Notably, in platyfish, which has a single chromosome PSD system, heterozygous males (XY) and females (WY or WX) have higher fitness than homozygotes (Orzack *et al*. 1980), providing evidence for overdominant fitness effects of sex chromosomes. Future work should evaluate the prevalence of proto-sex chromosomes with overdominant fitness effects in order to determine if the appropriate genetic variation exists for overdominance to maintain PSD more generally.

Regardless of the prevalence of overdominance in natural populations, it is one of the most straightforward mechanisms by which any polymorphism can be maintained (Hartl and Clark 2007). In my overdominant model, I assume that the proto-Y chromosomes carry male-beneficial additive or dominant alleles (Rice 1984, 1992), along with recessive deleterious alleles that have opposing effects equal in magnitude to the beneficial alleles (Charlesworth and Charlesworth 2000). Testing if these assumptions are biologically realistic would require measuring the magnitude and dominance of beneficial and deleterious alleles on proto-Y chromosomes. I also assume that the two homozygous genotypes (i.e., XX and YY) have equal fitness (Table 2), which may not be biologically realistic. Future work should consider how different fitness values of the two homozygotes affects the ability of overdominance to maintain polygenic sex determination.

When overdominant fitness effects in males maintain proto-sex chromosomes at frequencies most similar to those observed in natural populations, I find that they do so with female-deterious (additive) effects of the proto-Y chromosomes (Figure 5D; Supplemental Figure S16D). This is the strongest evidence I observe for sexually antagonistic fitness effects of individual proto-Y chromosomes (as opposed to multi-chromosomal genotypes) in any dominance scenario. It is also consistent with previous work demonstrating that overdominance in one sex can maintain genetic variants at individual loci even if there is directional selection against one allele in the other sex (Kidwell *et al*. 1977). In contrast to the additive and dominant fitness effects, there is no evidence for *ρMF<0* when overdominant fitness effects maintain PSD (Figure 4; Supplemental Figure S14). This provides additional evidence for a decoupling between single chromosome sexually antagonistic effects and sexual antagonism across multi-chromosome genotypes.

### Recombination on the proto-sex chromosomes and Y-linked alleles

My model makes assumptions about recombination on the sex chromosomes that may affect my conclusions about the maintenance of PSD. Specifically, I assume that there is no recombination between the male- or female-determining locus on each proto-Y or proto-W chromosome and the allele(s) under selection. Suppressed recombination is predicted to evolve in order to ensure that a sex-determining locus and sexually antagonistic alleles are inherited together on a young Y or W chromosome (Rice 1987) Suppressed recombination can also facilitate the invasion of a new sex determiner by increasing the effects of indirect selection on linked sexually antagonistic alleles, and it can further stabilize an ancestral sex chromosome through similar indirect effects (van Doorn and Kirkpatrick 2007, 2010). X-Y and Z-W recombination may be suppressed by chromosomal inversions that create tight genetic linkage between the sex-determining locus and sexually antagonistic alleles (Bergero and Charlesworth 2009; Wright *et al*. 2016).

While there is no direct evidence for inversions or other recombination suppressors on the house fly sex chromosomes, male meiosis in flies is thought to occur without crossing over (Gethmann 1988). A lack of crossing over in males would prevent X-Y recombination in males without the need for sex-chromosome-specific modifiers. However, there is evidence for male recombination in house fly (Feldmeyer *et al*. 2010), which opens up the possibility for X-Y recombination. Furthermore, in the absence of Y-linked suppressors, X-Y recombination could also occur in the multiple female genotypes that carry a proto-Y chromosome (Table 1).

There are at least three consequences of X-Y or Z-W recombination that could affect how sex-specific selection maintains PSD in my model. First, my model assumes that all copies of each proto-Y and proto-W chromosome have identical fitness effects. This assumption is consistent with population genetics theory that predicts Y-linked variants cannot segregate as protected polymorphisms when there is no X-Y recombination (Clark 1987). In contrast to this prediction, however, there are numerous examples of Y-linked polymorphisms with phenotypic effects in *Drosophila*, where the X and Y do not recombine (Clark 1990; Zhang *et al*. 2000; Lemos *et al*. 2008, 2010; Griffin *et al*. 2015; Brown *et al*. 2020). Relevant to the house fly, these examples include Y-linked variation with temperature-dependent effects that are distributed across the species’ geographic range (Rohmer *et al*. 2004). Future work could incorporate context-dependent variation in fitness effects of each proto-Y and proto-W chromosome into my model to assess how segregating polymorphisms on the proto-sex chromosomes affect the maintenance of PSD.

A second consequence of X-Y recombination could affect how overdominant fitness effects maintain PSD in my model. I assume that overdominance arises from proto-Y chromosomes that carry recessive deleterious alleles. This assumption is based on the prediction that suppressed X-Y recombination leads to the “degeneration” of the Y chromosome via accumulation of (recessive) deleterious alleles because of Muller’s ratchet and hitchhiking (Charlesworth and Charlesworth 2000; Bachtrog 2013). In contrast, X-Y recombination in males and/or females could help purge recessive deleterious mutations from the Y^M^ and III^M^ chromosomes (Muller 1964; Felsenstein 1974). Moreover, there are multiple house fly genotypes that are homozygous for a proto-Y chromosome (Table 1), which would expose recessive deleterious mutations to selection. Selection in these proto-Y chromosome homozygotes would further remove recessive deleterious alleles from populations (Charlesworth *et al*. 1990; Barrett and Charlesworth 1991). Therefore, while overdominant fitness effects are likely to maintain PSD, proto-Y chromosomes may not possess the necessary alleles for such selection pressures to act upon. However, overdominance can also occur by other mechanisms besides degeneration of the Y chromosome and independent of accumulation of recessive deleterious Y-linked alleles (Sellis *et al*. 2011; Connallon and Clark 2014). These other causes of overdominance may allow for heterozygote fitness advantages to maintain PSD without degeneration of the proto-Y chromosomes.

Third, X-Y recombination would introduce X-linked alleles onto the proto-Y chromosomes. Male-limited inheritance of a Y chromosome is expected to lead to the accumulation of male-beneficial alleles in Y-linked genes, even if they have female-deleterious effects (Rice 1987, 1996). Conversely, female-biased inheritance of the X chromosome is expected to result in stronger selection on X-linked alleles in females if those alleles are partially dominant or segregating at moderate frequencies (Rice 1984; Charlesworth *et al*. 1987; Orr and Betancourt 2001), potentially feminizing the X chromosome. If there is X-Y recombination, those feminized X-linked alleles would be continuously introduced onto the Y chromosome, possibly feminizing the Y as well (Cavoto *et al*. 2018). This could provide a biological mechanism that allows for female-beneficial and male-detrimental effects of the proto-Y chromosomes that I predict when additive fitness effects maintain PSD (Figure 5A; Supplemental Figure S16A).

### Low frequency proto-sex chromosomes are unlikely to be maintained by selection

My simulations suggest that it is difficult for selection to maintain proto-sex chromosomes at low frequencies within populations. For example, the III^M^ proto-Y represents <3% of all third chromosomes in each of the three natural populations I considered (Hamm *et al*. 2005; Meisel *et al*. 2016). In my simulations, many of the selection pressures that maintain III^M^ at a frequency similar to those observed in each of the populations for 1,000 generations eventually allow for the loss of III^M^ within 1,000,000 generations (Supplemental Figures S8, S9). Additive fitness effects are especially likely to allow for the loss of III^M^. In addition, when PSD is maintained with proto-sex chromosomes at frequencies similar to natural populations, III^M^ tends to be at a higher frequency in my simulations than in any of the actual populations (Figure 2; Supplemental Figure S3). III^M^ is also frequently lost when population size is finite, even when selection pressures exist that should maintain the polymorphism (Supplemental Figures S11, S12). IV^F^ is also found at a low frequency (2–9% of all fourth chromosomes in the natural populations); it is similarly found at a higher frequency in the simulations than in the actual populations (Figure 2; Supplemental Figure S3), and it is often lost in finite populations with selection (Supplemental Figures S11, S12).

The behavior of III^M^ and IV^F^ in my simulations suggests that natural selection within populations may not be sufficient to maintain low frequency proto-sex chromosomes, and other factors may be required. Variation in proto-sex chromosome frequencies across house fly populations are suggestive of what those factors may be. In particular, spatially variable, temporally fluctuating, and other heterogeneous selection pressures can contribute to the maintenance of genetic variation across populations (Levene 1953; Hedrick 2006), including maintaining PSD (Bateman and Anholt 2017). The house fly Y^M^ and III^M^ proto-Y chromosomes are distributed along north-south clines on multiple continents (Denholm *et al*. 1986; Tomita and Wada 1989; Hamm *et al*. 2005; Kozielska *et al*. 2008), suggesting spatially heterogeneous selection pressures may be important for maintaining PSD across house fly populations. Migration between these populations could continuously introduce rare proto-Y and proto-W chromosomes, potentially overwhelming selection pressures that would otherwise allow for the loss of those proto-sex chromosomes within populations (Lenormand 2002; Tigano and Friesen 2016). Future work should directly model the effect of migration across populations with different selection pressures on the maintenance of PSD within populations.

### Proto-sex chromosomes are reminiscent of pseudoautosomal regions, with some notable differences

Proto-sex chromosomes have some superficial similarities to pseudoautosomal regions (PARs) of heteromorphic sex chromosomes, within which there is X-Y (or Z-W) recombinatioin (Otto *et al*. 2011). Recombination in the PAR moves Y-linked alleles onto the X where they can be exposed to selection in females, reminiscent of how proto-Y chromosomes can be carried by females in a PSD system. Notably, PARs are capable of maintaining genetic variation, including sexually antagonistic alleles, under broader conditions than autosomes or even non-recombining regions of sex chromosomes (Jordan and Charlesworth 2012). These unique evolutionary dynamics of PARs are limited to loci close to the boundary with the non-recombining region of the sex chromosomes (Charlesworth *et al*. 2014). There is evidence for sexually antagonistic alleles in PARs from plants (Delph *et al*. 2010; Spigler *et al*. 2011) and fish (Tripathi *et al*. 2009; Kitano *et al*. 2009), but it is not yet clear whether PARs are enriched for sexually antagonistic alleles. Nonetheless, the possibility that PARs and proto-Y chromosomes are both hotspots of sexually antagonistic variation suggests they may experience similar selection pressures.

Despite the superficial similarities, there are important differences that make PARs imperfect analogs to proto-sex chromosomes. For example, when there is an epistatic dominant female determining proto-W chromosome (as in house fly), the proto-Y chromosome itself can be transmitted through females. In contrast, the Y PAR cannot ever be carried by females. Instead, Y-linked PAR alleles must first recombine onto the X PAR in order to be carried by females. Moreover, males can be homozygous for the proto-Y chromosome, which is superficially similar to a male carrying an X PAR that has an ancestrally Y-linked allele.

However, as above, this ancestral Y allele on the X PAR must recombine onto the X PAR, whereas, when there is PSD, homozygosity for the proto-Y occurs without X-Y recombination. Additional theoretical analyses are required to directly compare the invasion dynamics, fixation probabilities, and regions of stable polymorphisms across PARs and proto-sex chromosomes.

### Generalizing to even more complex polygenic sex determination systems

I anticipate that many of the general patterns I observe here would be similar in other complex PSD systems with at least three different proto-sex chromosomes. For example, in a system with more than two proto-Y chromosomes, I expect to observe similar effects of dominance on the maintenance of PSD and sexual antagonism as in the system I modeled with two proto-Y chromosomes. In addition, if there are multiple proto-W chromosomes and a single (epistatic) proto-Y (i.e., the opposite of the house fly system), I would expect similar patterns as I observe in the house fly system, but with the sexes reversed. These expectations could be tested in future work. It is not clear, however, what would be expected if there were both multiple proto-W and multiple proto-Y chromosomes in a population. It is worth noting that polygenic systems with many sex-determining loci are expected to be evolutionarily unstable (Rice 1986). Zebrafish may represent a promising model organism for combining empirical and theoretical approaches to make additional progress in understanding the maintenance of many sex determining loci (Liew *et al*. 2012; Wilson *et al*. 2014).

### Conclusions

My results demonstrate that the maintenance of PSD depends on the dominance of fitness effects, which are further predictive of the sexually antagonistic effects of proto-sex chromosomes and genotypes. Notably, overdominant fitness effects are more likely to maintain PSD than fitness effects with other types of dominance. However, that conclusion requires at least one of two important assumptions: genetic variation with overdominant fitness effects must exist in natural populations, or proto-Y chromosomes must carry recessive deleterious alleles (in addition to additive or dominant beneficial alleles). My results also suggest that sexually antagonistic multi-chromosomal genotypes should be most prominent when dominant fitness effects (and to a lesser extent additive fitness effects) maintain PSD. This sexual antagonism is an emergent property of multi-chromosomal genotypes, and not intrinsic to the fitness effects of all individual proto-Y chromosomes. In contrast, when overdominant fitness effects in males maintain PSD, the proto-Y chromosomes tend to be male-beneficial and female-detrimental, but the multi-chromosomal genotypes do not have opposing fitness effects in males and females. Lastly, future work is needed to evaluate how X-Y (and Z-W) recombination, heterogeneous selection pressures, and migration across demes affect the ability of sex-specific selection pressures to maintain complex PSD, including in systems with other combinations of proto-sex chromosomes.

## Acknowledgments

This work was funded by the National Science Foundation (DEB-1845686), and it was completed in part with the use of the Maxwell Cluster in the Research Computing Data Core at the University of Houston. This manuscript was improved by comments from editors, multiple anonymous reviewers, and members of the Meisel lab.

## Supplemental Figures

**Supplemental Figure S1.**
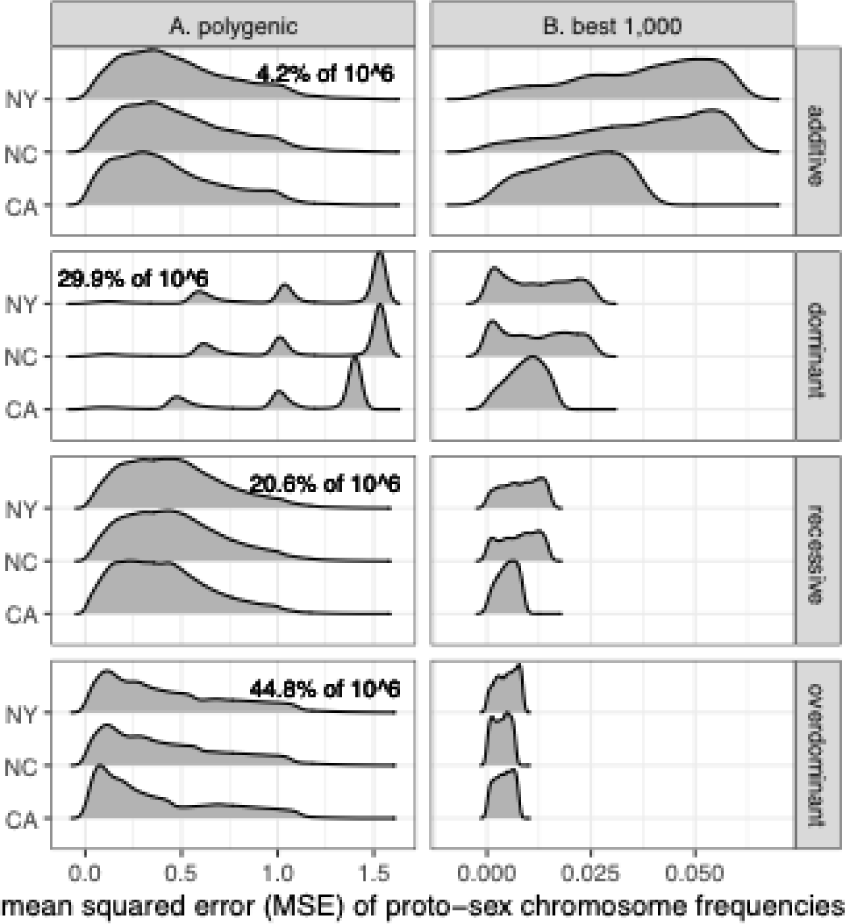
Smoothed histograms show the distributions of mean squared error (MSE) values for fitness arrays that maintain PSD (i.e., all proto-sex chromosomes at a frequency >0.1%). MSE is calculated based on the frequencies of Y^M^, III^M^, and IV^F^ found in either CA, NC, or NY. Simulations were started with equal frequencies of all genotypes. Fitness arrays are based on additive, dominant, recessive, or overdominant fitness effects of Y^M^ and III^M^. **(A)** Distributions of MSE are shown for all fitness arrays that maintain PSD, with the percent of those arrays that maintain PSD shown in each panel. **(B)** Distributions of MSE are shown for the 1,000 best-fitting arrays with the lowest MSE.

**Supplemental Figure S2.**
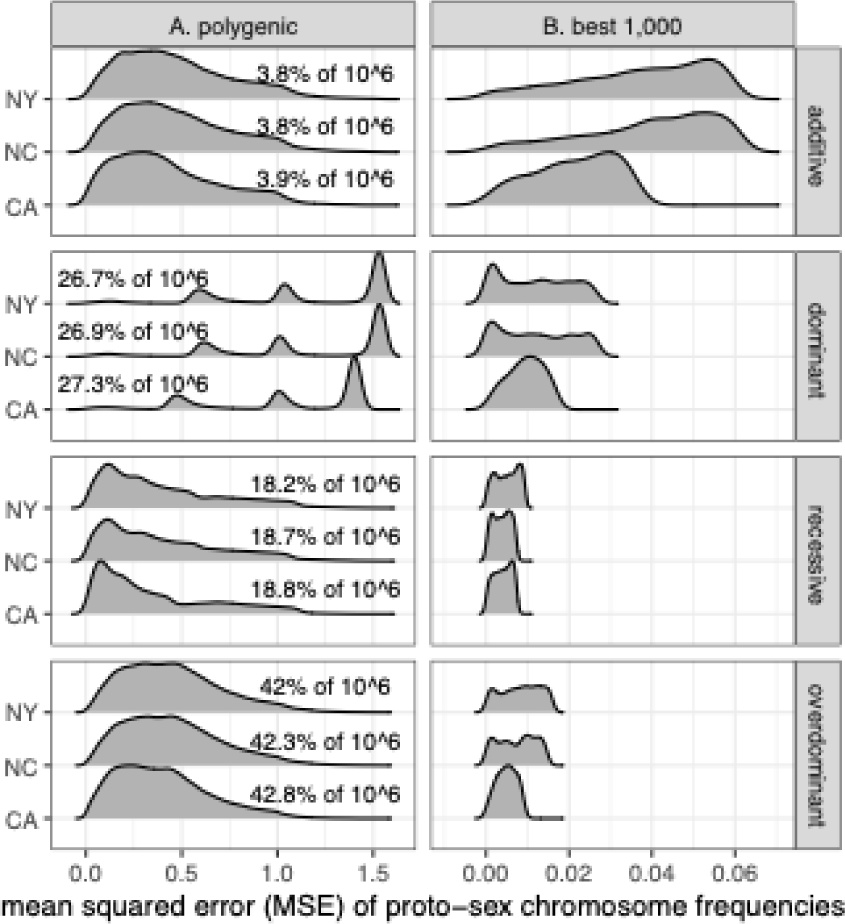
Smoothed histograms show the distributions of mean squared error (MSE) values for fitness arrays that maintain PSD (i.e., all proto-sex chromosomes at a frequency >0.1%). MSE is calculated based on the frequencies of Y^M^, III^M^, and IV^F^ found in either CA, NC, or NY. Simulations were started with genotype frequencies observed in natural populations. Fitness arrays are based on additive, dominant, recessive, or overdominant fitness effects of Y^M^ and III^M^. **(A)** Distributions of MSE are shown for all fitness arrays that maintain PSD, with the percent of those arrays that maintain PSD shown in each panel. **(B)** Distributions of MSE are shown for the 1,000 best-fitting arrays with the lowest MSE.

**Supplemental Figure S3.**
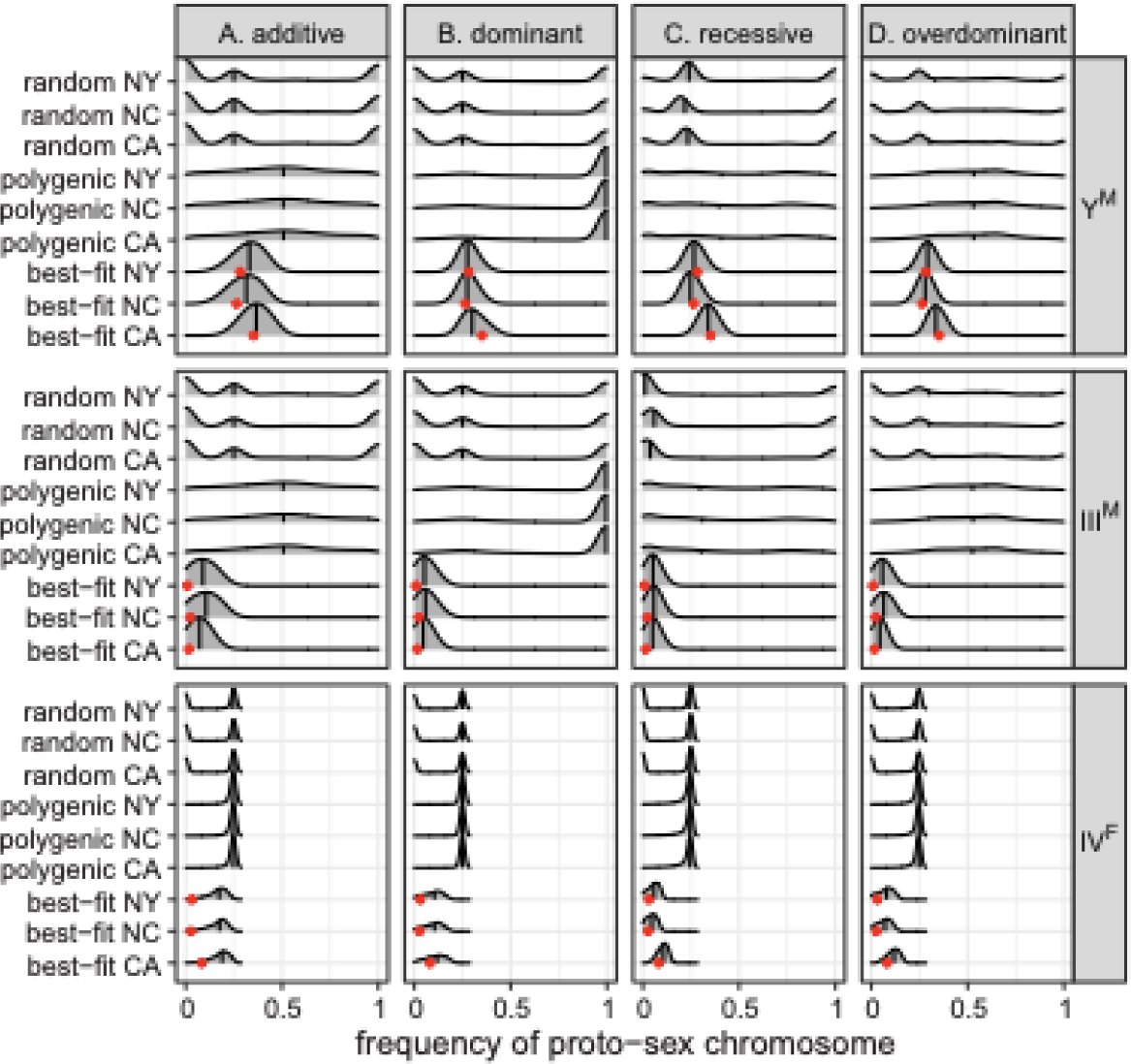
Smoothed histograms show the frequency of each proto-sex chromosome (Y^M^, III^M^, and IV^F^) after 1,000 generations in simulations using either 1,000 random fitness arrays (random), fitness arrays that maintain PSD (polygenic), or the 1,000 best-fitting fitness arrays for each population (CA, NC, or NY). Simulations were started with genotype frequencies observed in each population. The vertical line within each histogram shows the median. Red dots show the observed proto-sex chromosome frequencies in each natural population. Fitness arrays were calculated assuming either (**A)** additive, **(B)** dominant, **(C)** recessive, or **(D)** overdominant fitness effects of the proto-Ychromosomes.

**Supplemental Figure S4.**
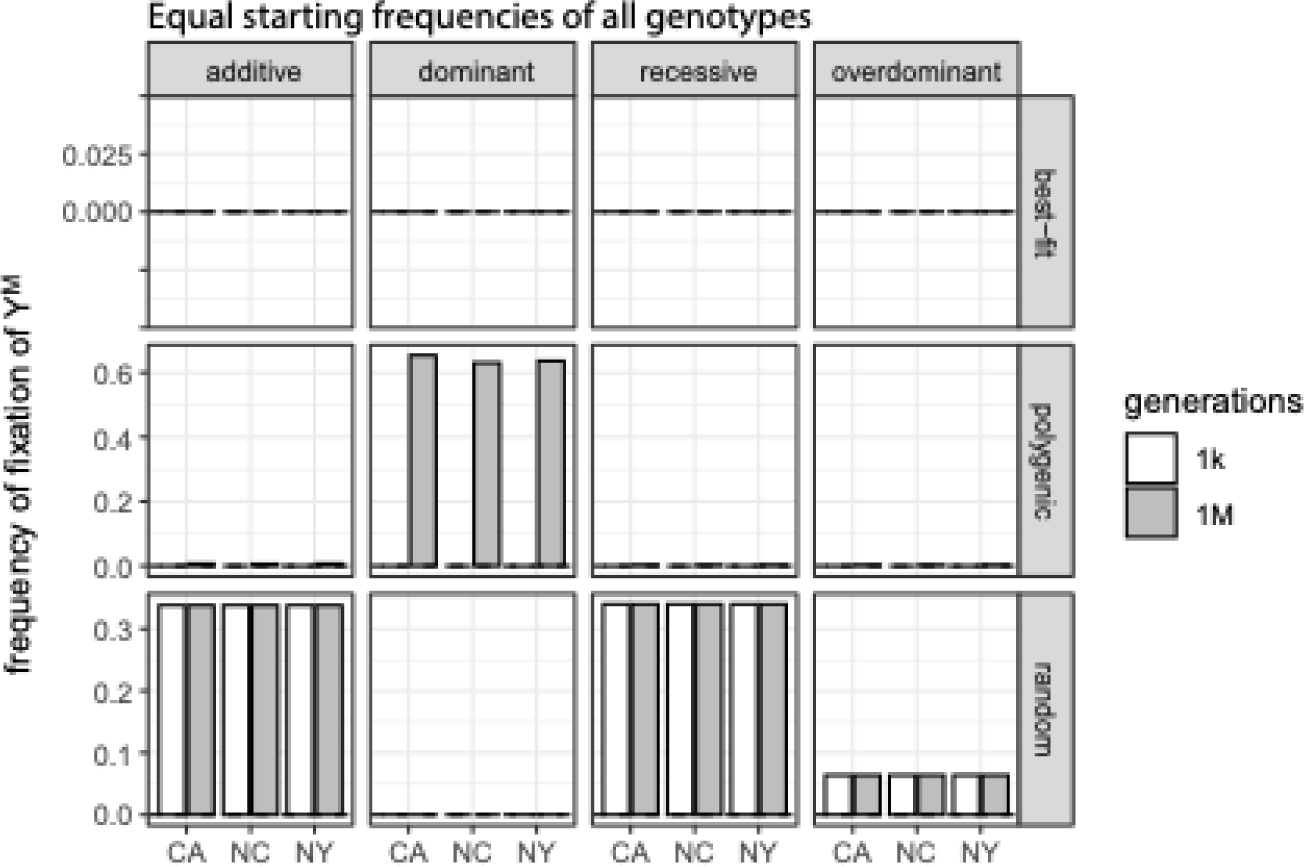
The frequency of fixation of the Y^M^ proto-Y chromosome after 1,000 generations (white) and 1,000,000 generations (gray) is shown for the 1,000 best-fitting genotype fitness arrays (best-fit), 1,000 fitness arrays that maintain PSD (polygenic), and 1,000 randomly chosen fitness arrays (random) in each population (CA, NC, or NY). Best-fitting fitness values and fitness values that maintain PSD were identified using simulation in which all genotypes started at equal frequencies. Fitness effects of each proto-Y chromosome were either additive, dominant, recessive, or overdominant.

**Supplemental Figure S5.**
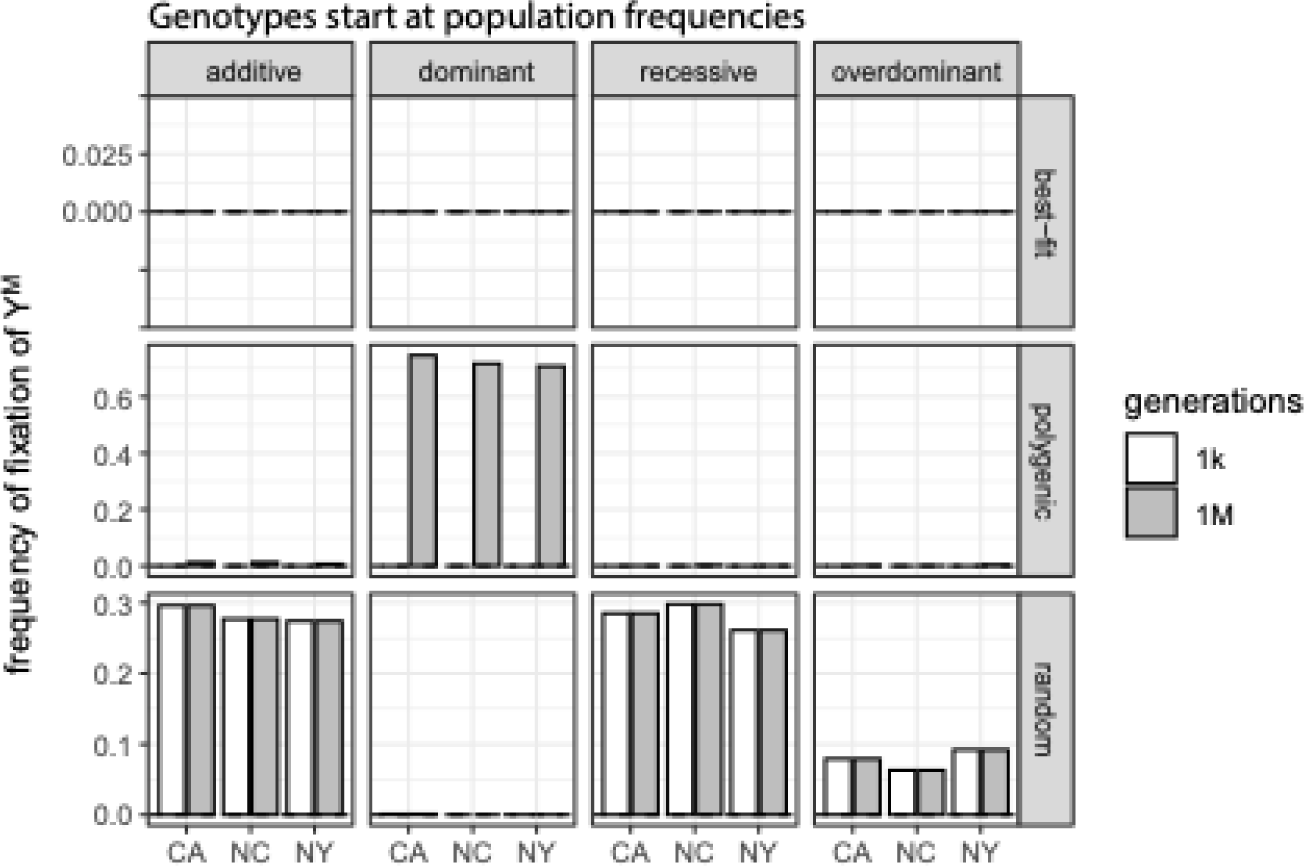
The frequency of fixation of the Y^M^ proto-Y chromosome after 1,000 generations (white) and 1,000,000 generations (gray) is shown for the 1,000 best-fitting genotype fitness arrays (best-fit), 1,000 fitness arrays that maintain PSD (polygenic), and 1,000 randomly chosen fitness arrays (random) in each population (CA, NC, or NY). Best-fitting fitness values and fitness values that maintain PSD were identified using simulation in which all genotypes started at the frequencies observed in each population. Fitness effects of each proto-Y chromosome were either additive, dominant, recessive, or overdominant.

**Supplemental Figure S6.**
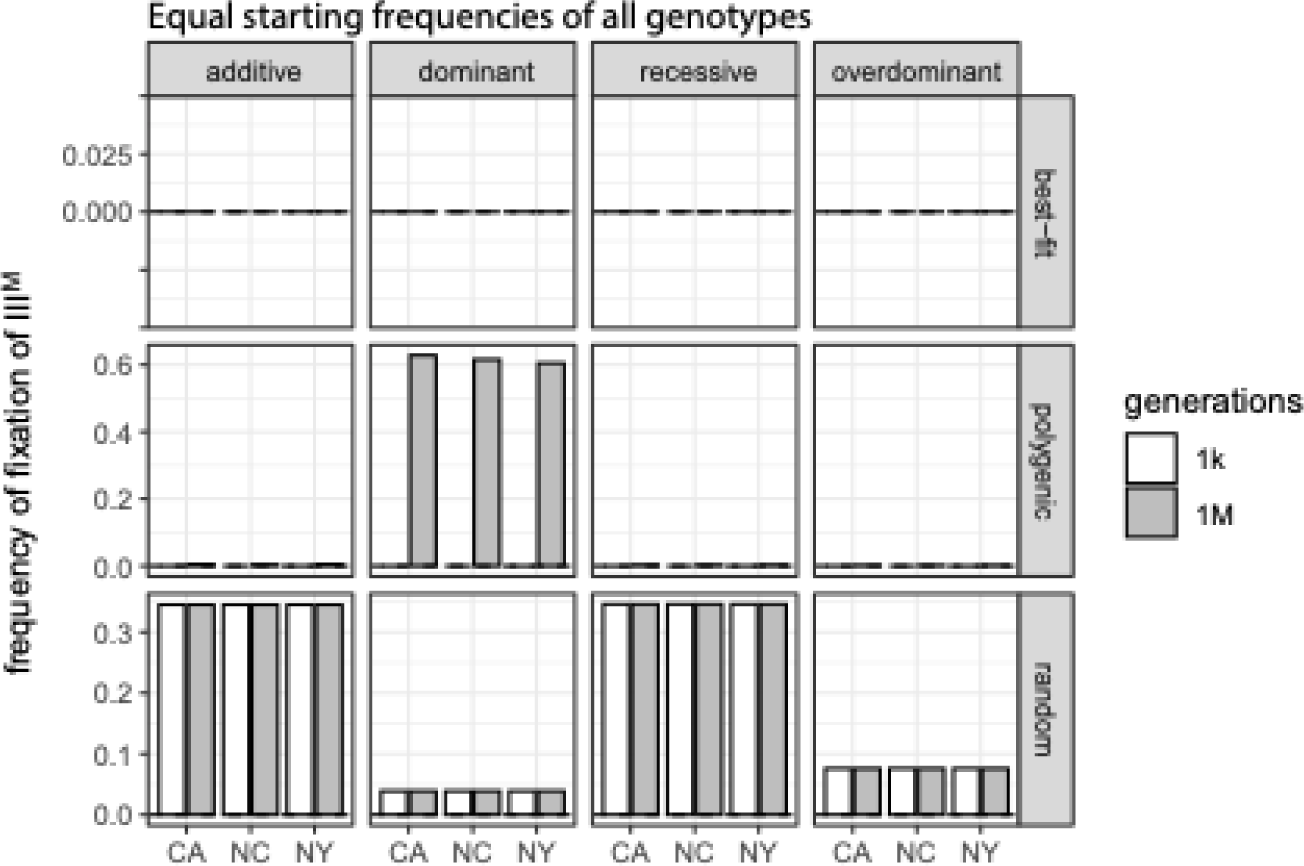
The frequency of fixation of the III^M^ proto-Y chromosome after 1,000 generations (white) and 1,000,000 generations (gray) is shown for the 1,000 best-fitting genotype fitness arrays (best-fit), 1,000 fitness arrays that maintain PSD (polygenic), and 1,000 randomly chosen fitness arrays (random) in each population (CA, NC, or NY). Best-fitting fitness values and fitness values that maintain PSD were identified using simulation in which all genotypes started at equal frequencies. Fitness effects of each proto-Y chromosome were either additive, dominant, recessive, or overdominant.

**Supplemental Figure S7.**
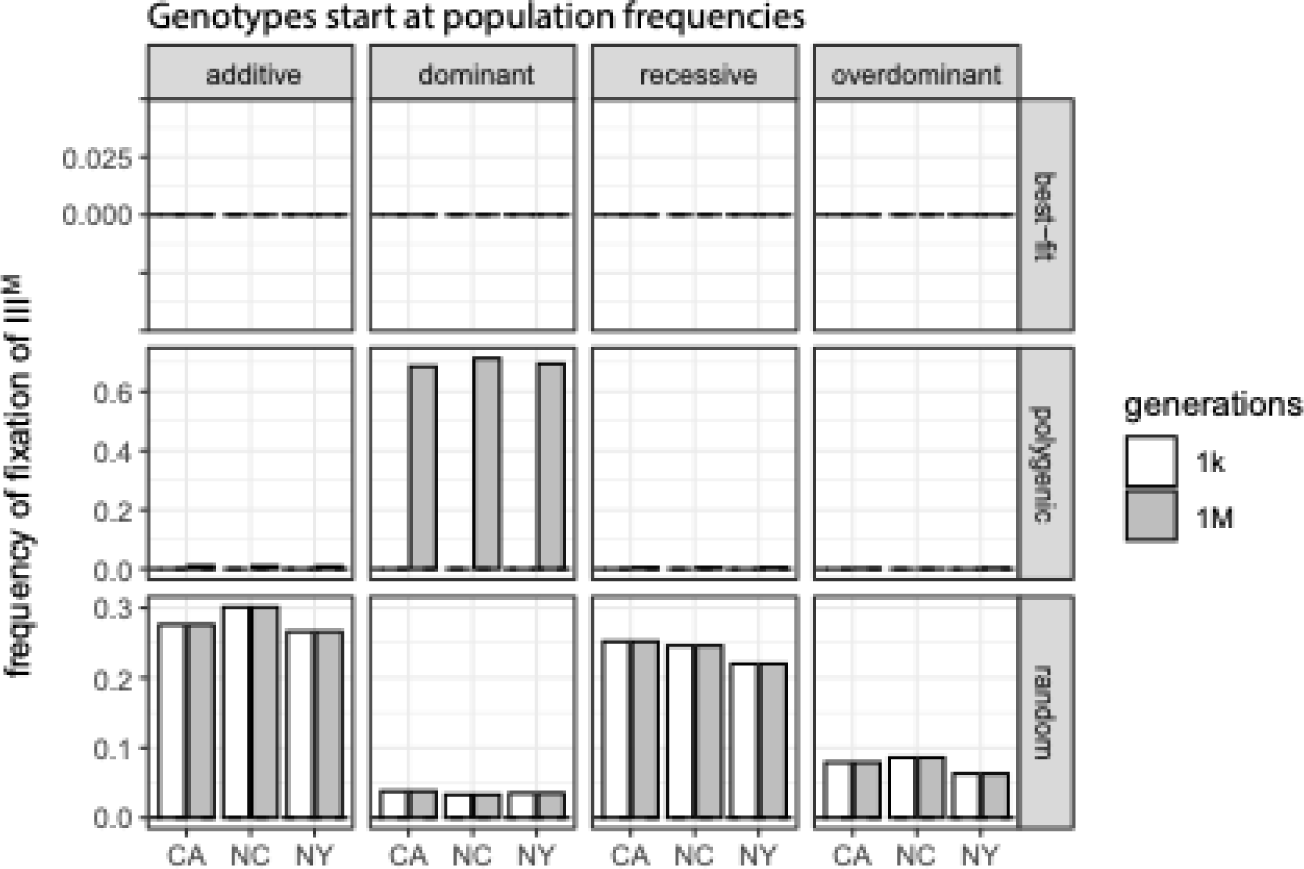
The frequency of fixation of the III^M^ proto-Y chromosome after 1,000 generations (white) and 1,000,000 generations (gray) is shown for the 1,000 best-fitting genotype fitness arrays (best-fit), 1,000 fitness arrays that maintain PSD (polygenic), and 1,000 randomly chosen fitness arrays (random) in each population (CA, NC, or NY). Best-fitting fitness values and fitness values that maintain PSD were identified using simulation in which all genotypes started at the frequencies observed in each population. Fitness effects of each proto-Y chromosome were either additive, dominant, recessive, or overdominant.

**Supplemental Figure S8.**
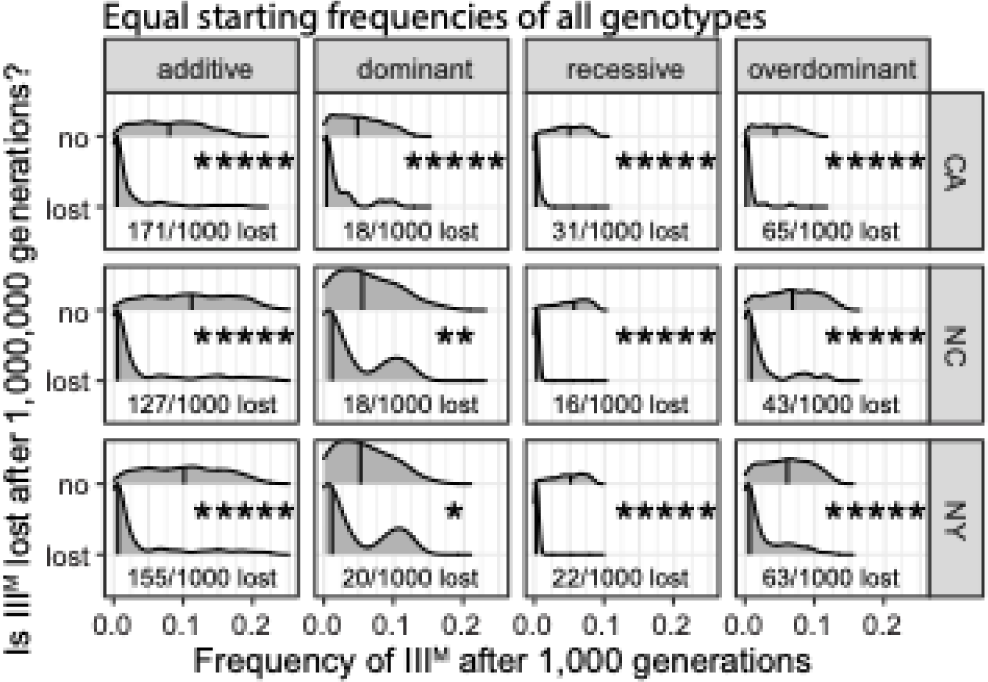
Histograms of the frequency of the III^M^ proto-Y chromosome after 1,000 generations are shown for the 1,000 fitness arrays that produce proto-sex chromosomes most similar to those observed in each natural population (CA, NC, or NY). Fitness arrays are divided into those in which the III^M^ chromosome eventually reaches a frequency <0.1% after 1,000,000 generations (lost) and those in which the III^M^ chromosome is maintained at a frequency >0.1% (no). Fractions in each panel indicate how often the III^M^ chromosome was lost after 1,000,000 generations in the 1,000 best-fitting arrays. Simulations were started with equal frequencies of all genotypes, and fitness effects of proto-Y chromosomes were either additive, dominant, recessive, or overdominant.

**Supplemental Figure S9.**
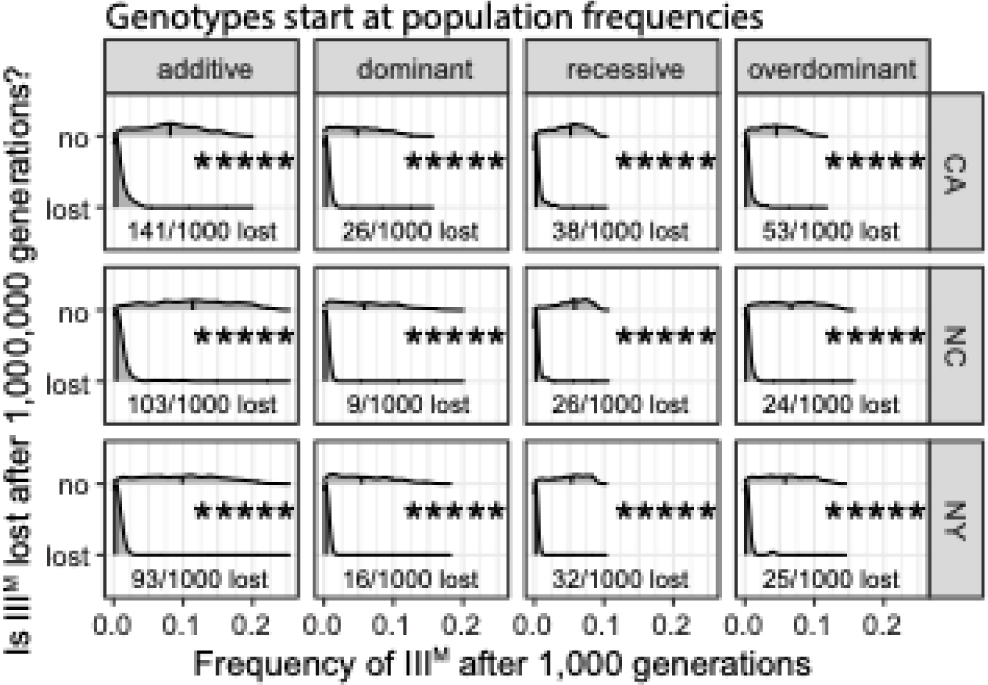
Histograms of the frequency of the III^M^ proto-Y chromosome after 1,000 generations are shown for the 1,000 fitness arrays that produce proto-sex chromosomes most similar to those observed in each natural population (CA, NC, or NY). Fitness arrays are divided into those in which the III^M^ chromosome eventually reaches a frequency <0.1% after 1,000,000 generations (lost) and those in which the III^M^ chromosome is maintained at a frequency >0.1% (no). Fractions in each panel indicate how often the III^M^ chromosome was lost after 1,000,000 generations in the 1,000 best-fitting arrays. Simulations were started with genotypes frequencies observed in each population, and fitness effects of proto-Y chromosomes were either additive, dominant, recessive, or overdominant.

**Supplemental Figure S10.**
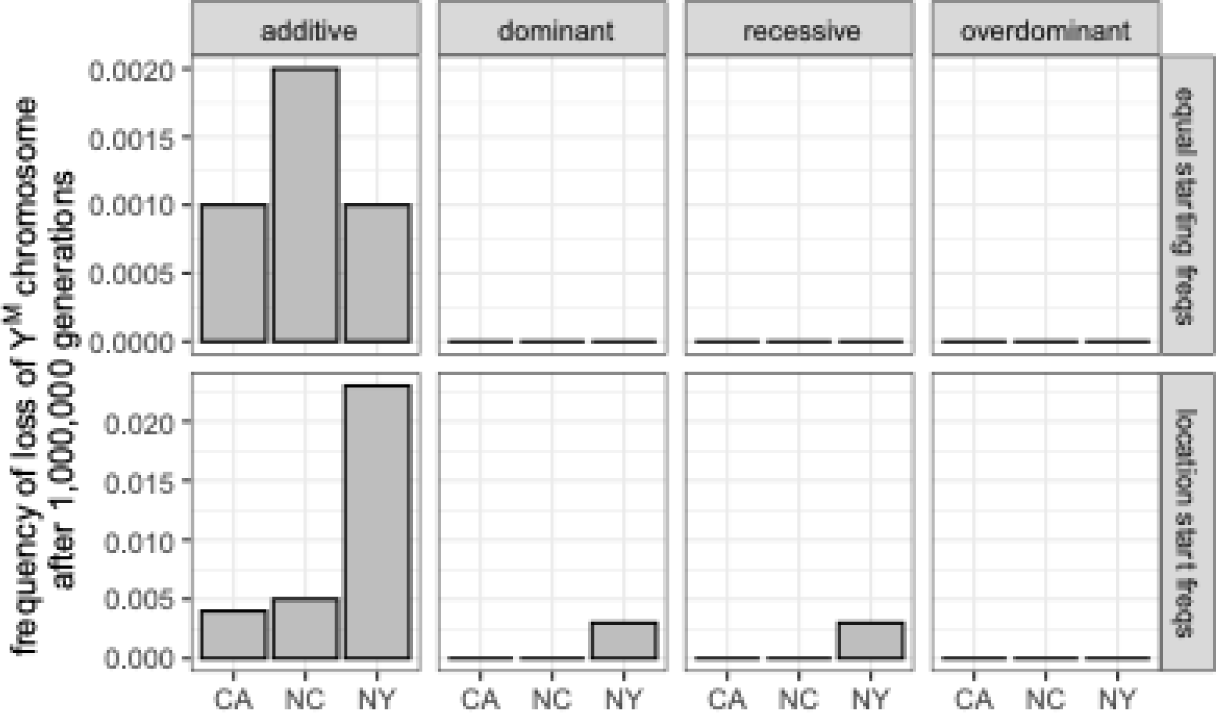
Bar graphs show the frequency with which the Y^M^ chromosome is lost after 1,000,000 generation using fitness effects that maintain proto-sex chromosome frequencies similar to those observed in each natural population (CA, NC, or NY). The best-fitting arrays were identified with simulations starting with equal frequencies of all genotypes (equal starting freqs) or frequencies observed in each population (location start freqs). Genotype fitness values were calculating assuming either additive, dominant, recessive, or overdominant fitness effects.

**Supplemental Figure S11.**
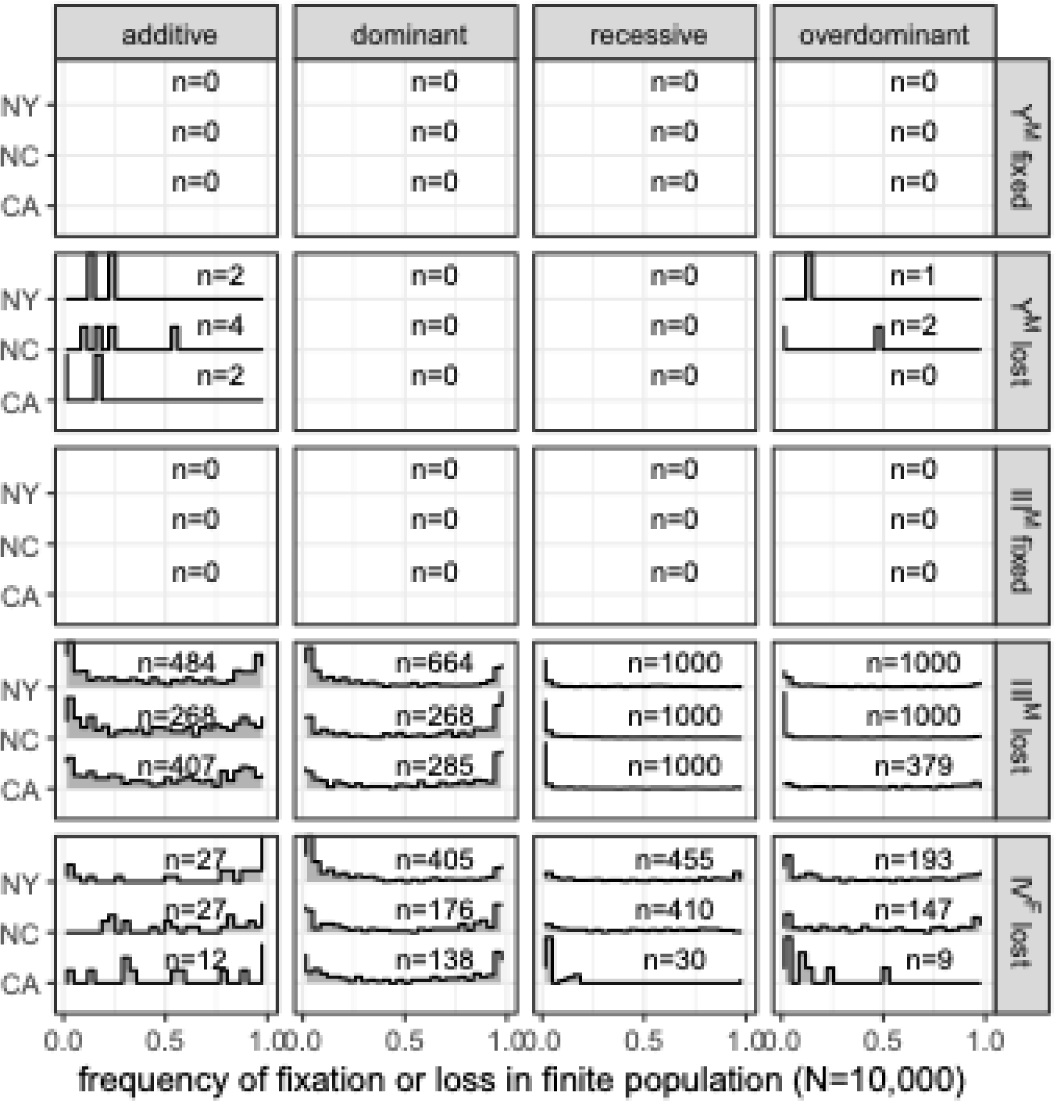
Histograms show the distributions of frequency of fixation or loss each proto-Y or proto-W chromosome in each population when *N*=10^4^. Numbers in each panel (n) indicate the number of fitness arrays with at least one fixation in 100 simulations. The x-axis is the frequency of fixations in 100 simulations for those arrays with at least one fixation. Simulations were performed with the 1,000 best-fitting selection pressures for each population (CA, NC, or NY) that were identified from simulations started with equal frequencies of all genotypes, using either **(A)** additive, **(B)** dominant, **(C)** recessive, or **(D)** overdominant fitness effects of the proto-Y chromosomes.

**Supplemental Figure S12.**
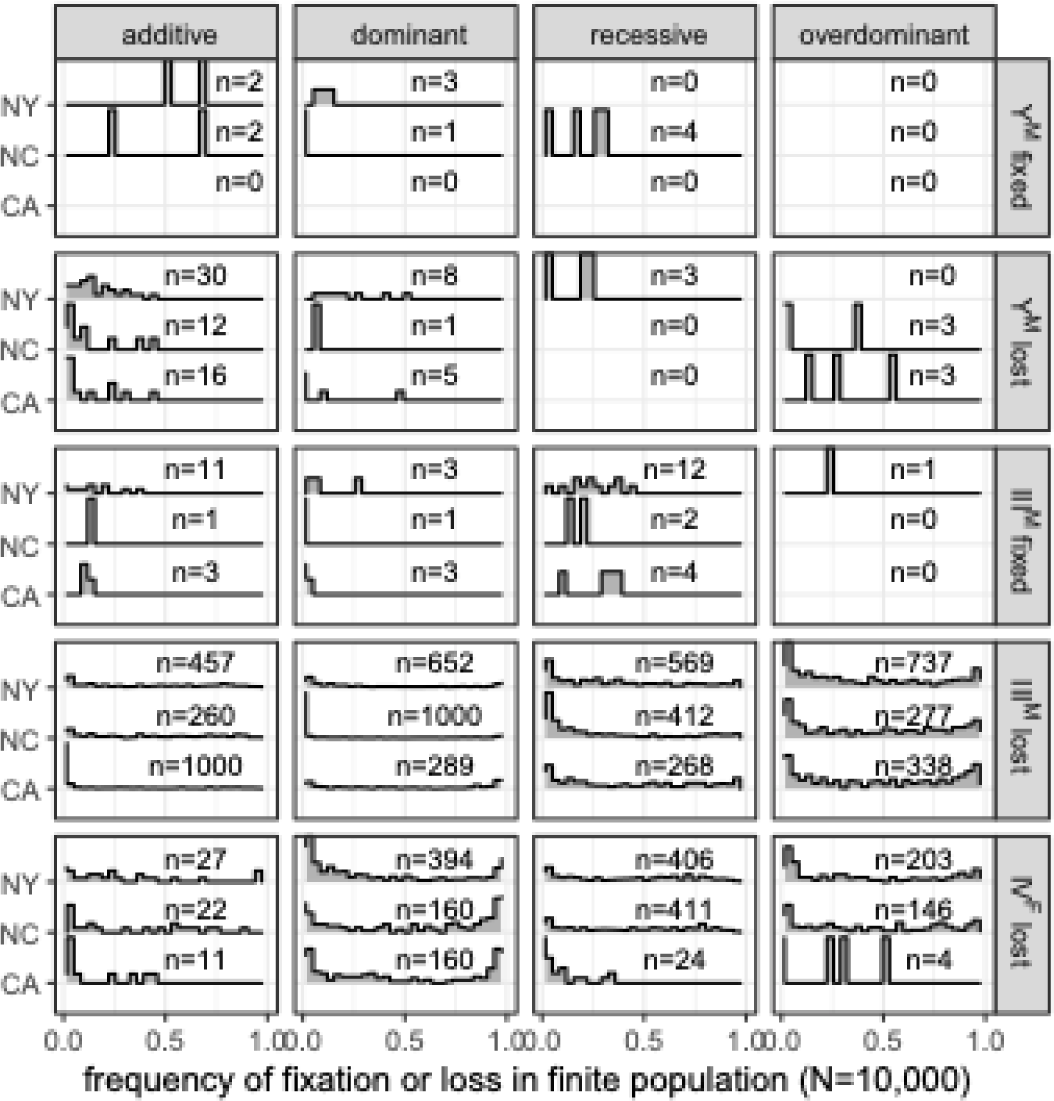
Histograms show the distributions of frequency of fixation or loss each proto-Y or proto-W chromosome in each population when *N*=10^4^. Numbers in each panel (n) indicate the number of fitness arrays with at least one fixation in 100 simulations. The x-axis is the frequency of fixations in 100 simulations for those arrays with at least one fixation. Simulations were performed with the 1,000 best-fitting selection pressures for each population (CA, NC, or NY) that were identified from simulations started with the observed frequencies of each genotype in each population, using either **(A)** additive, **(B)** dominant, **(C)** recessive, or **(D)** overdominant fitness effects of the proto-Y chromosomes.

**Supplemental Figure S13.**
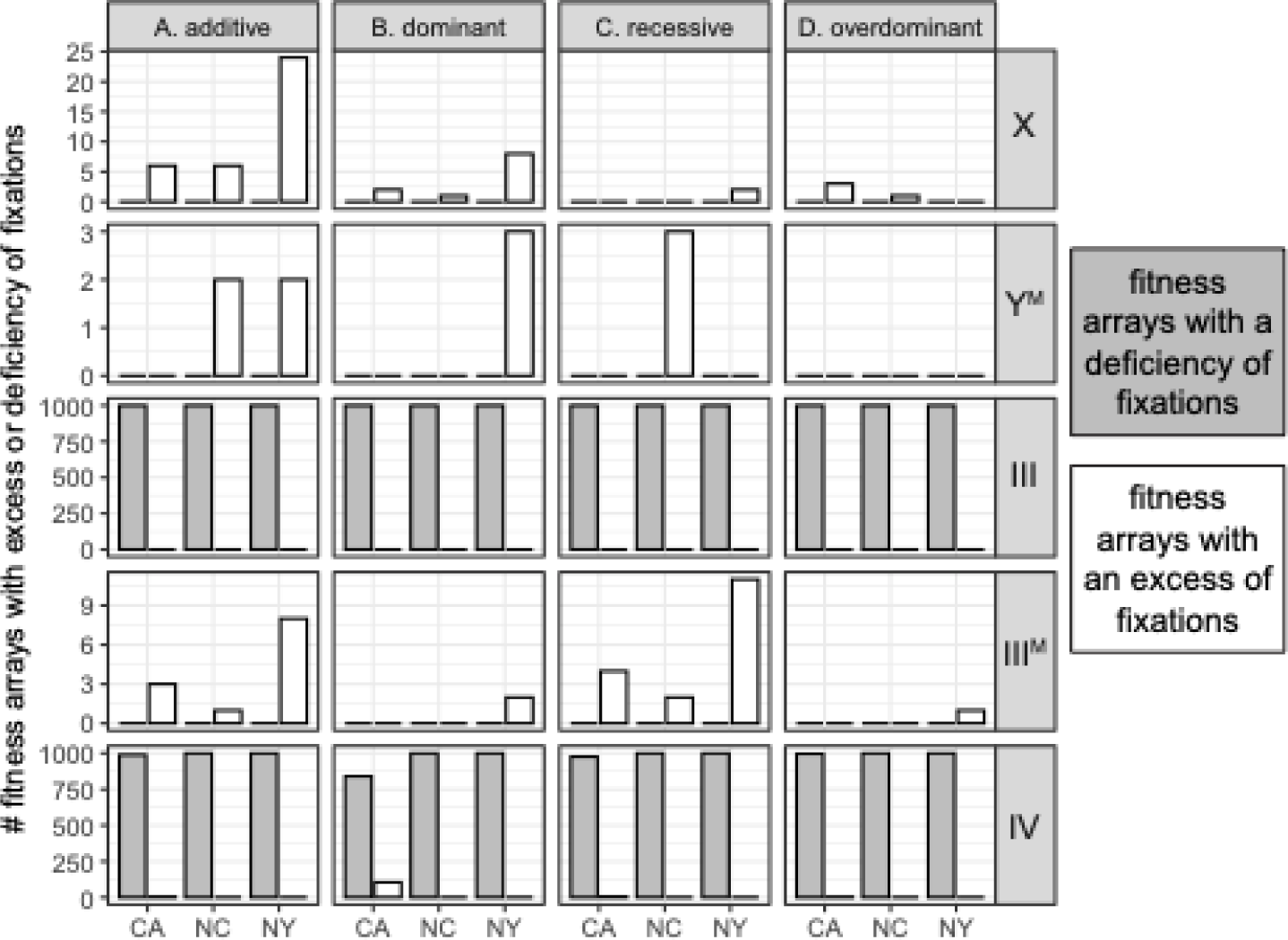
Bar plots show the number of fitness arrays that result in a deficiency (gray) or excess (white) of fixations of each proto-sex chromosome (X, Y^M^, III, III^M^, or IV) relative to simulations in which there are no fitness differences across proto-sex chromosomes (i.e., drift only). The fitness arrays are the 1,000 best-fitting fitness values for each population (CA, NC, or NY) when simulations were started with genotype frequencies observed in natural populations. Fitness effects of the proto-Y chromosomes are either **(A)** additive, **(B)** dominant, **(C)** recessive, or **(D)** overdominant.

**Supplemental Figure S14.**
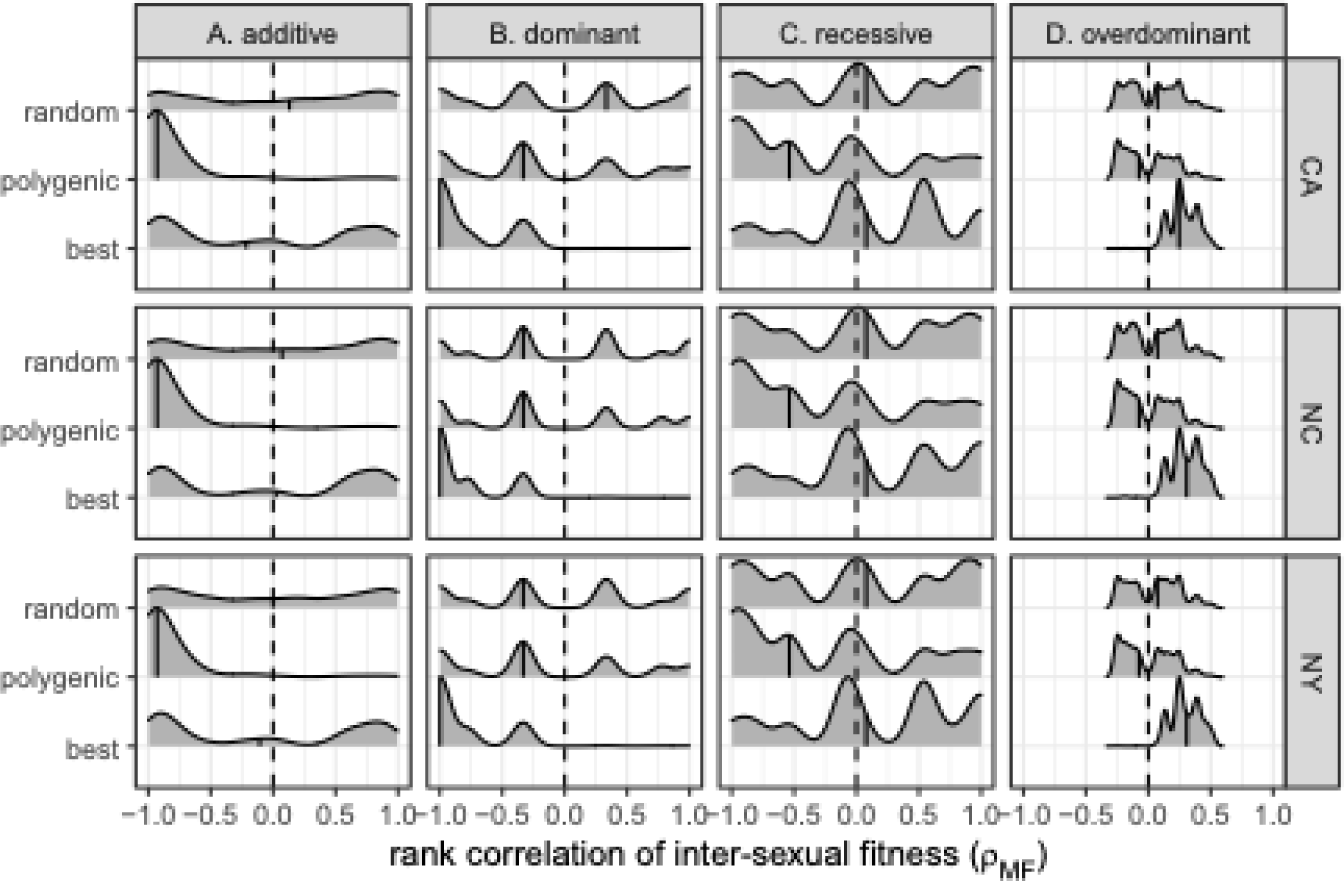
Smoothed histograms show intersexual fitness correlations (*ρMF*) for the 1,000 best-fitting genotypic fitness arrays for the CA, NC, and NY populations (best), all fitness arrays that maintain PSD (polygenic), and 1,000 random fitness arrays. Simulations were started with genotype frequencies observed in natural populations. Dashed vertical lines show *ρMF=0*, and solid vertical lines within histograms show the median. Fitness arrays were calculated assuming either (**A)** additive, **(B)** dominant, **(C)** recessive, or **(D)** overdominant fitness effects of the proto-Y chromosomes.

**Supplemental Figure S15.**
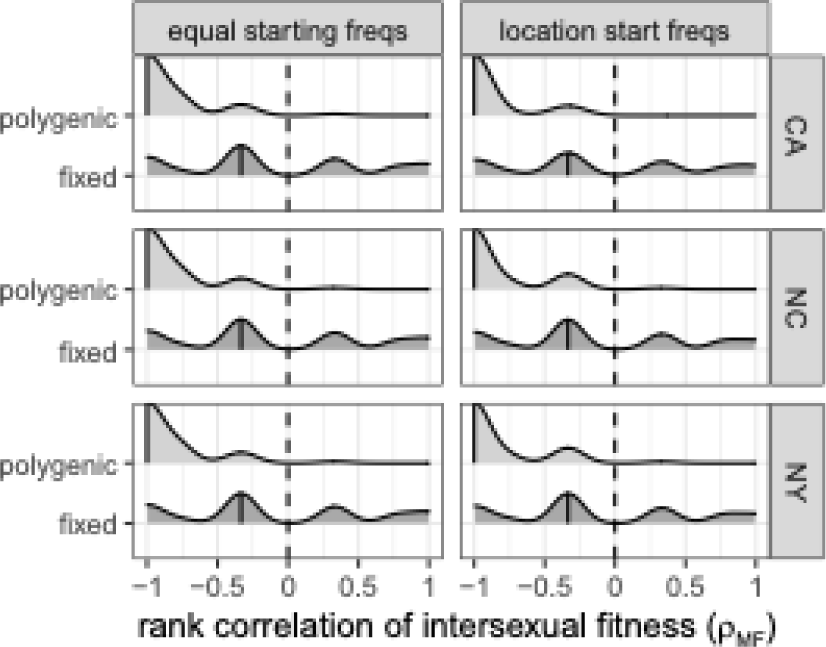
Smoothed histograms show intersexual fitness correlations (*ρMF*) for the 1,000 best-fitting genotypic fitness arrays based on dominant fitness effects of the proto-Y chromosomes for the CA, NC, and NY populations after 1,000,000 generations. Fitness arrays that maintain PSD for 1,000 generations are divided into those the maintain PSD for 1,00,000 generations (polygenic) and those that allow at least one proto-sex chromosome to reach fixation (fixed). Simulations were started with either equal frequencies of all genotypes (left) or genotype frequencies observed in natural populations (right). Dashed vertical lines show *ρMF=0*, and solid vertical lines within histograms show the median.

**Supplemental Figure S16.**
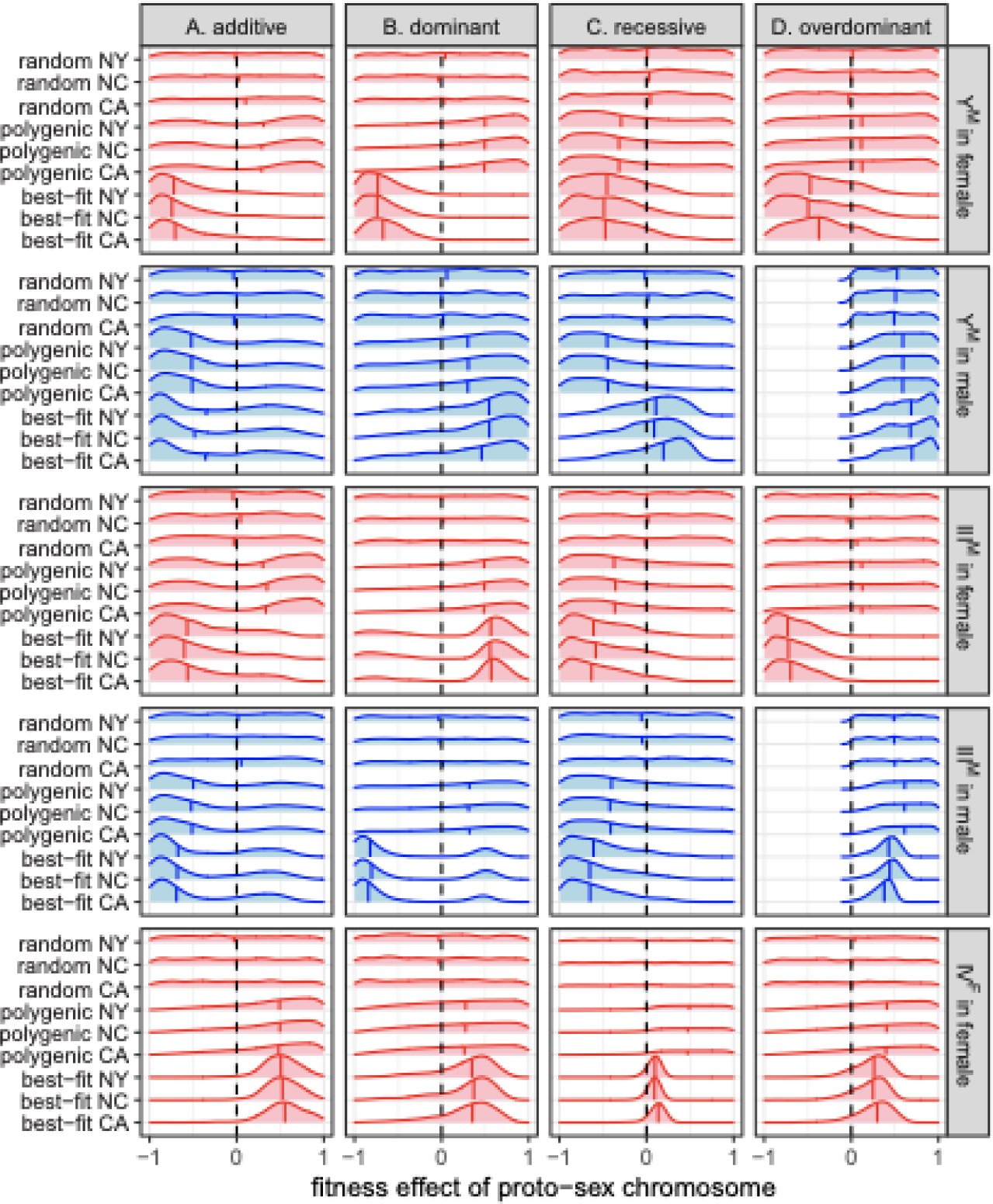
Smoothed histograms show the distributions of fitness effects of each proto-sex chromosome in each sex for 1,000 random fitness arrays (random), fitness arrays that maintain PSD (polygenic), or the 1,000 best-fitting fitness arrays for each population (CA, NC, or NY). Simulations were started with genotype frequencies observed in natural populations. The vertical line within each histogram shows the median, and dashed lines in each panel are at fitness value of 0. Fitness arrays were calculated assuming either (**A)** additive, **(B)** dominant, **(C)** recessive, or **(D)** overdominant fitness effects of the proto-Y chromosomes.

**Supplemental Figure S17.**
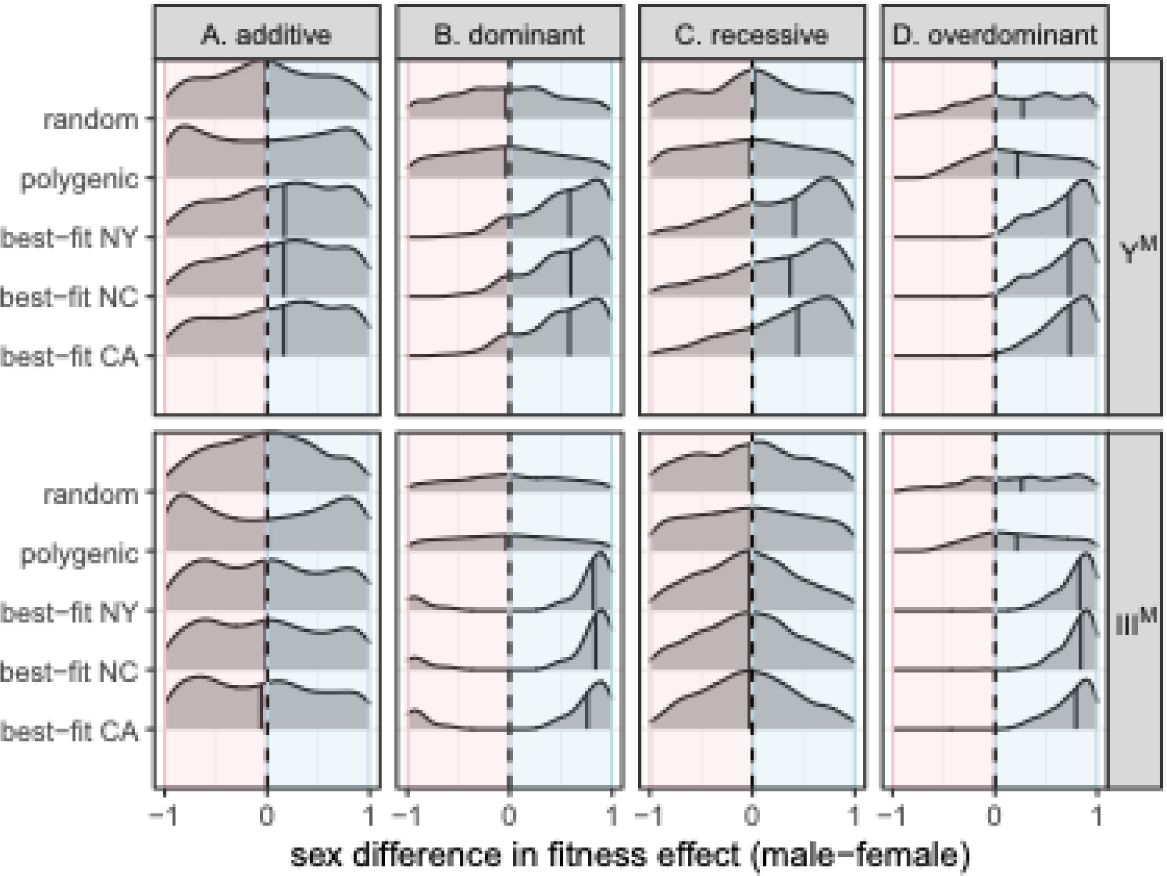
Smoothed histograms show the distributions of the difference in the fitness effect of each proto-Y chromosome (Y^M^ or III^M^) between males and females within individual parameterizations of the model. If the fitness effect is greater in males, the values are positive. If the fitness effect is greater in females, the values are negative. Fitness effects of each proto-Y chromosome come from the 1,000 random fitness arrays (random), fitness arrays that maintain PSD (polygenic), or the 1,000 best-fitting fitness arrays for each population (CA, NC, or NY). Simulations were started with equal frequencies of all genotypes. The vertical line within each histogram shows the median, and dashed lines in each panel are a difference of 0 (i.e., equal fitness effects in males and females). Genotype fitness arrays were calculated assuming either (**A)** additive, **(B)** dominant, **(C)** recessive, or **(D)** overdominant fitness effects of the proto-Y chromosomes.

**Supplemental Figure S18.**
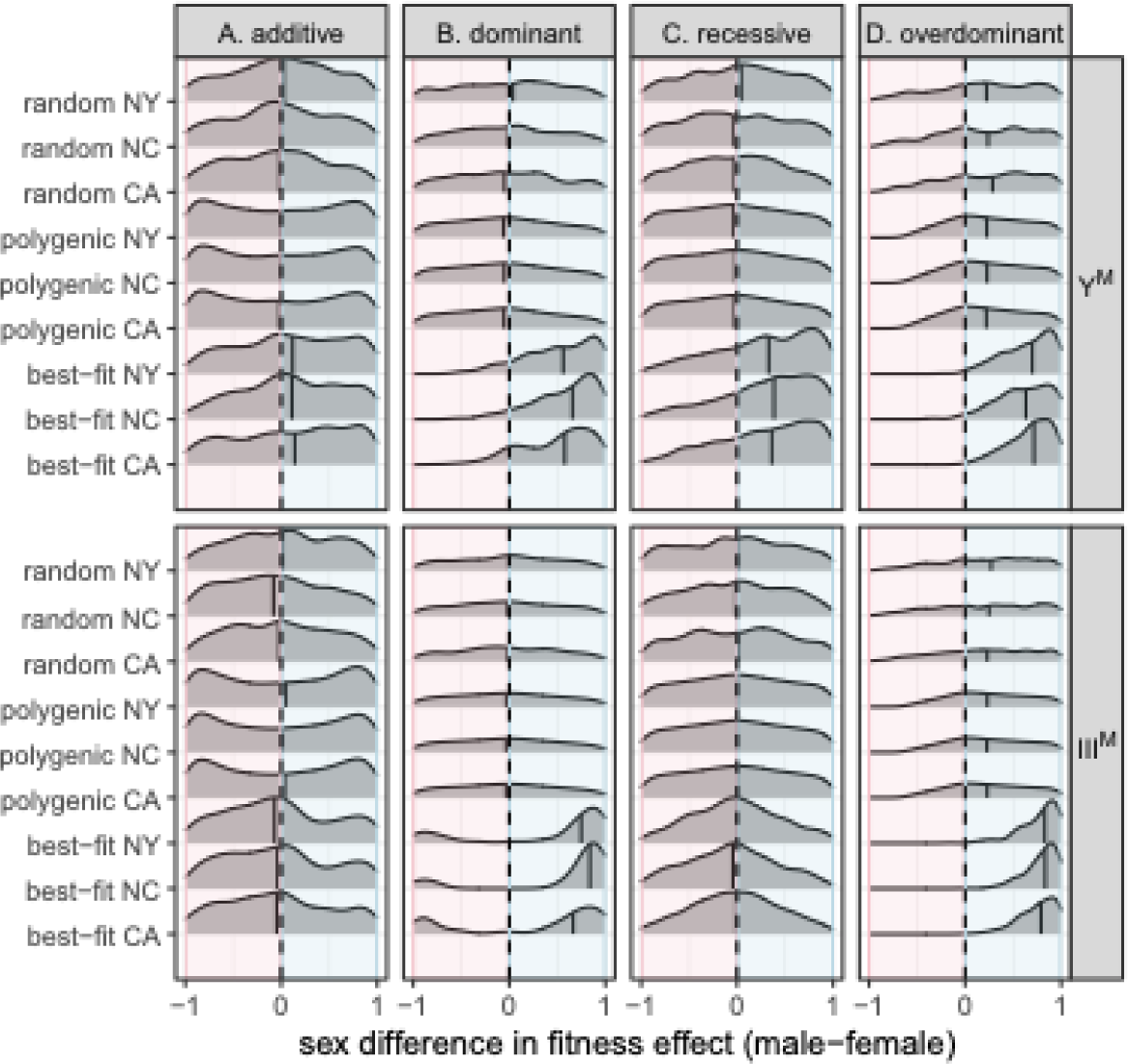
Smoothed histograms show the distributions of the difference in the fitness effect of each proto-Y chromosome (Y^M^ or III^M^) between males and females within individual parameterizations of the model. If the fitness effect is greater in males, the values are positive. If the fitness effect is greater in females, the values are negative. Fitness effects of each proto-Y chromosome come from the 1,000 random fitness arrays (random), fitness arrays that maintain PSD (polygenic), or the 1,000 best-fitting fitness arrays for each population (CA, NC, or NY). Simulations were started with genotype frequencies observed in natural populations. The vertical line within each histogram shows the median, and dashed lines in each panel are a difference of 0 (i.e., equal fitness effects in males and females). Genotype fitness arrays were calculated assuming either (**A)** additive, **(B)** dominant, **(C)** recessive, or **(D)** overdominant fitness effects of the proto-Y chromosomes.

**Supplemental Figure S19.**
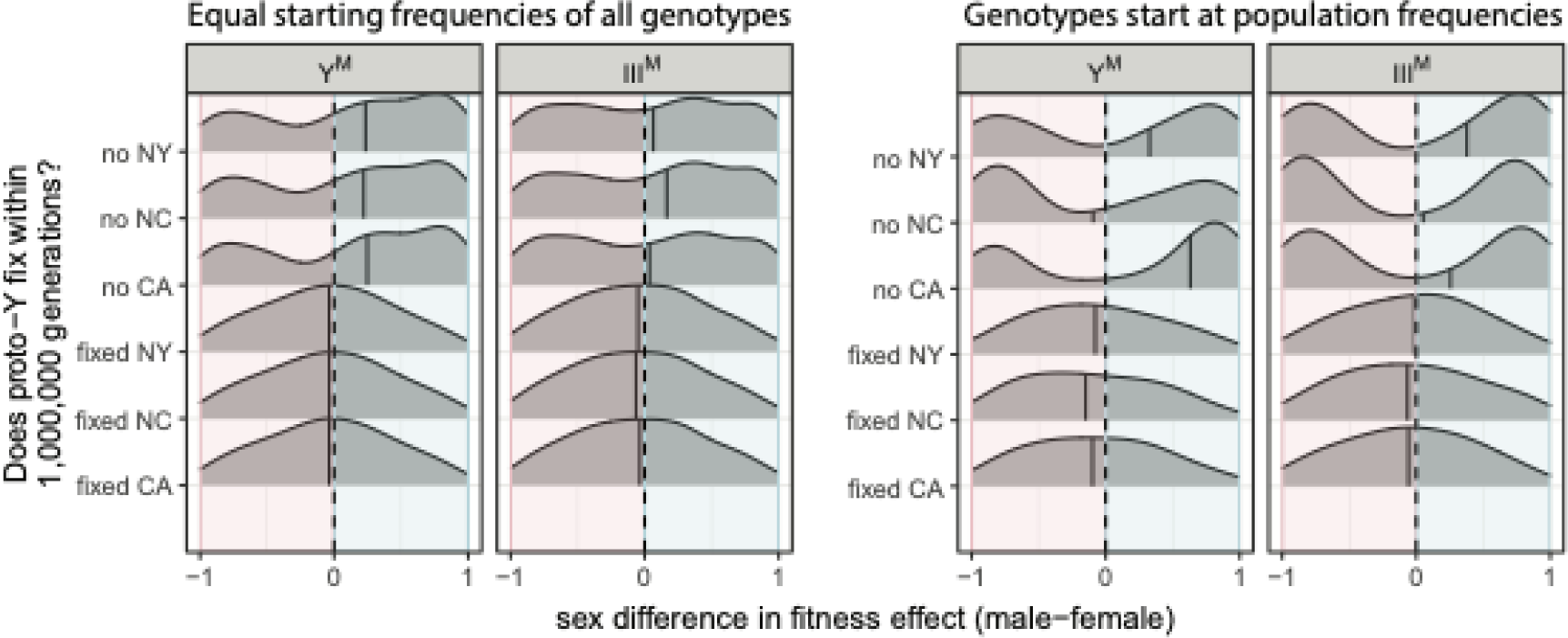
Smoothed histograms show the distributions of the difference in the fitness effect of each proto-Y chromosome (Y^M^ or III^M^) between males and females within individual parameterizations of the model. If the fitness effect is greater in males, the values are positive. If the fitness effect is greater in females, the values are negative. Simulations were performed using 1,000 fitness arrays based on dominant fitness effects of the proto-Y chromosomes that maintain PSD for 1,000 generations. Fitness arrays that maintain PSD for 1,000 generations are divided into those in which the proto-Y chromosome reaches fixation within 1,00,000 generations (fixed) and those in which it does not reach fixation (no) in each population (CA, NC, or NY). Simulations were started with either equal frequencies of all genotypes (left) or genotype frequencies observed in natural populations (right). The vertical line within each histogram shows the median, and dashed lines in each panel are a difference of 0 (i.e., equal fitness effects in males and females).

**Supplemental Figure S20.**
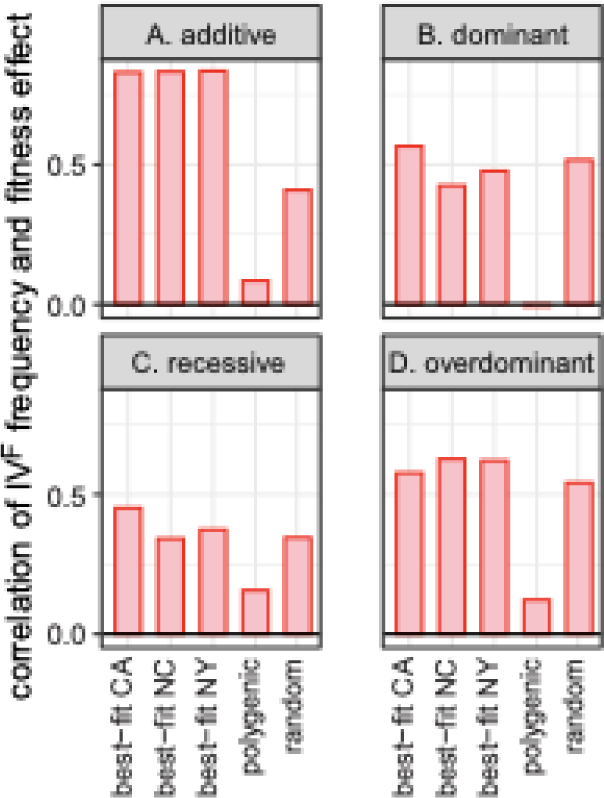
Bar plots show Spearman’s rank order correlation between the frequency the IV^F^ proto-W chromosome across simulated populations and its fitness effect in the populations. Simulated populations include those that best-fit proto-sex chromosome frequencies in natural populations (CA, NC, or NY), those that maintain PSD (polygenic), and 1,000 random populations (random). Simulations were started with equal frequencies of all genotypes. Genotype fitness in populations was calculated assuming **(A)** additive, **(B)** dominant, **(C)** recessive, or **(D)** overdominant effects of the proto-Y chromosomes.

**Supplemental Figure S21.**
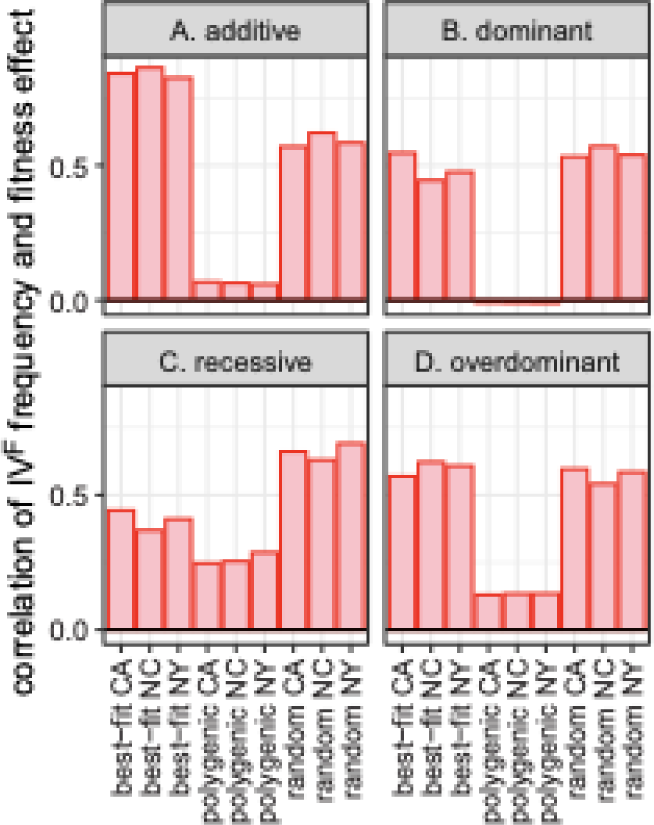
Bar plots show Spearman’s rank order correlation between the frequency the IV^F^ proto-W chromosome across simulated populations and its fitness effect in the populations. Simulated populations include those that best-fit proto-sex chromosome frequencies in natural populations (CA, NC, or NY), those that maintain PSD (polygenic), and 1,000 random populations (random). Simulations were started with the observed frequencies of each genotype in each population. Genotype fitness in populations was calculated assuming **(A)** additive, **(B)** dominant, **(C)** recessive, or **(D)** overdominant effects of the proto-Y chromosomes.

## Supplemental Table

**Supplemental Table S1.** Recursion equations to simulate random mating.

| genotype | frequency | frequency after random mating |
| --- | --- | --- |
| X/X; III/III; IV/IV | $f_1$ | $\begin{aligned} & f_1 m_1/2 + f_1 m_3/2 + f_1 m_4/4 \\ & + f_2 m_1/4 + f_2 m_3/4 + f_2 m_4/8 \\ & + f_3 m_1/4 + f_3 m_3/8 + f_3 m_4/16 \\ & + f_5 m_1/8 + f_5 m_3/8 + f_5 m_4/16 \\ & + f_6 m_1/16 + f_6 m_3/16 + f_6 m_4/32 \end{aligned}$ |
| X/X; III/III; IV/IV<sup>F</sup> | $f_2$ | $\begin{aligned} & f_2 m_1/4 + f_2 m_3/4 + f_2 m_4/8 \\ & + f_3 m_1/8 + f_3 m_3/8 + f_3 m_4/16 \\ & + f_5 m_1/8 + f_5 m_3/8 + f_5 m_4/16 \\ & + f_6 m_1/16 + f_6 m_3/16 + f_6 m_4/32 \end{aligned}$ |
| X/X; III/III<sup>M</sup>; IV/IV<sup>F</sup> | $f_3$ | $\begin{aligned} & f_2 m_1/4 + f_2 m_2/2 + f_2 m_4/8 + f_2 m_5/4 \\ & + f_3 m_1/4 + f_3 m_2/4 + f_3 m_3/8 \\ & + f_3 m_4/8 + f_3 m_5/8 \\ & + f_4 m_1/4 + f_4 m_3/4 + f_4 m_4/8 \\ & + f_5 m_1/8 + f_5 m_2/4 + f_5 m_4/16 + f_5 m_5/8 \\ & + f_6 m_1/8 + f_6 m_2/8 + f_6 m_3/16 \\ & + f_6 m_4/16 + f_6 m_5/16 \\ & + f_7 m_1/8 + f_7 m_3/8 + f_7 m_4/16 \end{aligned}$ |
| X/X; III<sup>M</sup>/III<sup>M</sup>; IV/IV<sup>F</sup> | $f_4$ | $\begin{aligned} & f_3 m_1/8 + f_3 m_2/4 + f_3 m_4/16 + f_3 m_5/8 \\ & + f_4 m_1/4 + f_4 m_2/2 + f_4 m_4/8 + f_4 m_5/4 \\ & + f_6 m_1/16 + f_6 m_2/8 + f_6 m_4/32 + f_6 m_5/16 \\ & + f_7 m_1/8 + f_7 m_2/4 + f_7 m_4/16 + f_7 m_5/8 \end{aligned}$ |
| X/Y<sup>M</sup>; III/III; IV/IV<sup>F</sup> | $f_5$ | $\begin{aligned} & f_2 m_3/4 + f_2 m_4/8 + f_2 m_6/2 + f_2 m_7/4 \\ & + f_3 m_3/8 + f_3 m_4/16 + f_3 m_6/4 + f_3 m_7/8 \\ & + f_5 m_1/8 + f_5 m_3/4 + f_5 m_4/8 \\ & + f_5 m_6/4 + f_5 m_7/8 \\ & + f_6 m_1/16 + f_6 m_3/8 + f_6 m_4/16 \\ & + f_6 m_6/8 + f_6 m_7/16 \\ & + f_8 m_1/4 + f_8 m_3/4 + f_8 m_4/8 \\ & + f_9 m_1/8 + f_9 m_3/8 + f_9 m_4/16 \end{aligned}$ |
| $X/Y^M; III/III^M; IV/IV^F$ | $f_6$ | $\begin{aligned} & f_2m_4/8 + f_2m_5/4 + f_2m_7/4 + f_2m_8/2 \\ & + f_3m_3/8 + f_3m_4/8 + f_3m_5/8 \\ & + f_3m_6/4 + f_3m_7/4 + f_3m_8/4 \\ & + f_4m_3/4 + f_4m_4/8 + f_4m_6/2 + f_4m_7/4 \\ & + f_5m_1/8 + f_5m_2/4 + f_5m_4/8 \\ & + f_5m_5/4 + f_5m_7/8 + f_5m_8/4 \\ & + f_6m_1/8 + f_6m_2/8 + f_6m_3/8 + f_6m_4/8 \\ & + f_6m_5/8 + f_6m_6/8 + f_6m_7/8 + f_6m_8/8 \\ & + f_7m_1/8 + f_7m_3/4 + f_7m_4/8 \\ & + f_7m_6/4 + f_7m_7/8 \\ & + f_8m_1/4 + f_8m_2/2 + f_8m_4/8 + f_8m_5/4 \\ & + f_9m_1/4 + f_9m_2/4 + f_9m_3/8 \\ & + f_9m_4/8 + f_9m_5/8 \\ & + f_{10}m_1/4 + f_{10}m_3/4 + f_{10}m_4/8 \end{aligned}$ |
| $X/Y^M; III^M/III^M; IV/IV^F$ | $f_7$ | $\begin{aligned} & f_3m_4/16 + f_3m_5/8 + f_3m_7/8 + f_3m_8/4 \\ & + f_4m_4/8 + f_4m_5/4 + f_4m_7/4 + f_4m_8/2 \\ & + f_6m_1/16 + f_6m_2/8 + f_6m_4/16 \\ & + f_6m_5/8 + f_6m_7/16 + f_6m_8/8 \\ & + f_7m_1/8 + f_7m_2/4 + f_7m_4/8 \\ & + f_7m_5/4 + f_7m_7/8 + f_7m_8/4 \\ & + f_9m_1/8 + f_9m_2/4 + f_9m_4/16 + f_9m_5/8 \\ & + f_{10}m_1/4 + f_{10}m_2/2 + f_{10}m_4/8 + f_{10}m_5/4 \end{aligned}$ |
| $Y^M/Y^M; III/III; IV/IV^F$ | $f_8$ | $\begin{aligned} & f_5m_3/8 + f_5m_4/16 + f_5m_6/4 + f_5m_7/8 \\ & + f_6m_3/16 + f_6m_4/32 + f_6m_6/8 + f_6m_7/16 \\ & + f_8m_3/4 + f_8m_4/8 + f_8m_6/2 + f_8m_7/4 \\ & + f_9m_3/8 + f_9m_4/16 + f_9m_6/4 + f_9m_7/8 \end{aligned}$ |
| $Y^M/Y^M; III/III^M; IV/IV^F$ | $f_9$ | $\begin{aligned} & f_5m_4/16 + f_5m_5/8 + f_5m_7/8 + f_5m_8/4 \\ & + f_6m_3/16 + f_6m_4/16 + f_6m_5/16 \\ & + f_6m_6/8 + f_6m_7/8 + f_6m_8/8 \\ & + f_7m_3/8 + f_7m_4/16 + f_7m_6/4 + f_7m_7/8 \\ & + f_8m_4/8 + f_8m_5/4 + f_8m_7/4 + f_8m_8/2 \\ & + f_9m_3/8 + f_9m_4/8 + f_9m_5/8 \\ & + f_9m_6/4 + f_9m_7/4 + f_9m_8/4 \\ & + f_{10}m_3/4 + f_{10}m_4/8 + f_{10}m_6/2 + f_{10}m_7/4 \end{aligned}$ |
| $Y^M/Y^M; III^M/III^M;$<br>$IV/IV^F$ | $f_{10}$ | $\begin{aligned} & f_6 m_4/32 + f_6 m_5/16 + f_6 m_7/16 + f_6 m_8/8 \\ & + f_7 m_4/16 + f_7 m_5/8 + f_7 m_7/8 + f_7 m_8/4 \\ & + f_9 m_4/16 + f_9 m_5/8 + f_9 m_7/8 + f_9 m_8/4 \\ & + f_{10} m_4/8 + f_{10} m_5/4 + f_{10} m_7/4 + f_{10} m_8/2 \end{aligned}$ |
| $X/X; III/III^M; IV/IV$ | $m_1$ | $\begin{aligned} & f_1 m_1/2 + f_1 m_2/1 + f_1 m_4/4 + f_1 m_5/2 \\ & + f_2 m_1/4 + f_2 m_2/2 + f_2 m_4/8 + f_2 m_5/4 \\ & + f_3 m_1/8 + f_3 m_2/4 + f_3 m_3/8 \\ & + f_3 m_4/8 + f_3 m_5/8 \\ & + f_4 m_1/4 + f_4 m_3/4 + f_4 m_4/8 \\ & + f_5 m_1/8 + f_5 m_2/4 + f_5 m_4/16 + f_5 m_5/8 \\ & + f_6 m_1/8 + f_6 m_2/8 + f_6 m_3/16 \\ & + f_6 m_4/16 + f_6 m_5/16 \\ & + f_7 m_1/8 + f_7 m_3/8 + f_7 m_4/16 \end{aligned}$ |
| $X/X; III^M/III^M; IV/IV$ | $m_2$ | $\begin{aligned} & f_3 m_1/8 + f_3 m_2/4 + f_3 m_4/16 + f_3 m_5/8 \\ & + f_4 m_1/4 + f_4 m_2/2 + f_4 m_4/8 + f_4 m_5/4 \\ & + f_6 m_1/16 + f_6 m_2/8 + f_6 m_4/32 + f_6 m_5/16 \\ & + f_7 m_1/8 + f_7 m_2/4 + f_7 m_4/16 + f_7 m_5/8 \end{aligned}$ |
| $X/Y^M; III/III; IV/IV$ | $m_3$ | $\begin{aligned} & f_1 m_3/2 + f_1 m_4/4 + f_1 m_6/1 + f_1 m_7/2 \\ & + f_2 m_3/4 + f_2 m_4/8 + f_2 m_6/2 + f_2 m_7/4 \\ & + f_3 m_3/8 + f_3 m_4/16 + f_3 m_6/4 + f_3 m_7/8 \\ & + f_5 m_1/8 + f_5 m_3/4 + f_5 m_4/8 \\ & + f_5 m_6/4 + f_5 m_7/8 \\ & + f_6 m_1/16 + f_6 m_3/8 + f_6 m_4/16 \\ & + f_6 m_6/8 + f_6 m_7/16 \\ & + f_8 m_1/4 + f_8 m_3/4 + f_8 m_4/8 \\ & + f_9 m_1/8 + f_9 m_3/8 + f_9 m_4/16 \\ & + f_5 m_6/4 + f_5 m_7/8 \\ & + f_6 m_1/16 + f_6 m_3/8 + f_6 m_4/16 \\ & + f_6 m_6/8 + f_6 m_7/16 \\ & + f_8 m_1/4 + f_8 m_3/4 + f_8 m_4/8 \\ & + f_9 m_1/8 + f_9 m_3/8 + f_9 m_4/16 \end{aligned}$ |
| $X/Y^M; III/III^M; IV/IV$ | $m_4$ | $\begin{aligned} & f_1 m_4/4 + f_1 m_5/2 + f_1 m_7/2 + f_1 m_8/1 \\ & + f_2 m_4/8 + f_2 m_5/4 + f_2 m_7/4 + f_2 m_8/2 \\ & \quad + f_3 m_3/8 + f_3 m_4/8 + f_3 m_5/8 \\ & \quad + f_3 m_6/4 + f_3 m_7/4 + f_3 m_8/4 \\ & + f_4 m_3/4 + f_4 m_4/8 + f_4 m_6/2 + f_4 m_7/4 \\ & \quad + f_5 m_1/8 + f_5 m_2/4 + f_5 m_4/8 \\ & \quad + f_5 m_5/4 + f_5 m_7/8 + f_5 m_8/4 \\ & + f_6 m_1/8 + f_6 m_2/8 + f_6 m_3/8 + f_6 m_4/8 \\ & + f_6 m_5/8 + f_6 m_6/8 + f_6 m_7/8 + f_6 m_8/8 \\ & \quad + f_7 m_1/8 + f_7 m_3/4 + f_7 m_4/8 \\ & \quad + f_7 m_6/4 + f_7 m_7/8 \\ & + f_8 m_1/4 + f_8 m_2/2 + f_8 m_4/8 + f_8 m_5/4 \\ & \quad + f_9 m_1/4 + f_9 m_2/4 + f_9 m_3/8 \\ & \quad + f_9 m_4/8 + f_9 m_5/8 \\ & + f_{10} m_1/4 + f_{10} m_3/4 + f_{10} m_4/8 \end{aligned}$ |
| $X/Y^M; III^M/III^M; IV/IV$ | $m_5$ | $\begin{aligned} & f_3 m_4/16 + f_3 m_5/8 + f_3 m_7/8 + f_3 m_8/4 \\ & + f_4 m_4/8 + f_4 m_5/4 + f_4 m_7/4 + f_4 m_8/2 \\ & \quad + f_6 m_1/16 + f_6 m_2/8 + f_6 m_4/16 \\ & \quad + f_6 m_5/8 + f_6 m_7/16 + f_6 m_8/8 \\ & \quad + f_7 m_1/8 + f_7 m_2/4 + f_7 m_4/8 \\ & \quad + f_7 m_5/4 + f_7 m_7/8 + f_7 m_8/4 \\ & + f_9 m_1/8 + f_9 m_2/4 + f_9 m_4/16 + f_9 m_5/8 \\ & + f_{10} m_1/4 + f_{10} m_2/2 + f_{10} m_4/8 + f_{10} m_5/4 \end{aligned}$ |
| $Y^M/Y^M; III/III; IV/IV$ | $m_6$ | $\begin{aligned} & f_5 m_3/8 + f_5 m_4/16 + f_5 m_6/4 + f_5 m_7/8 \\ & + f_6 m_3/16 + f_6 m_4/32 + f_6 m_6/8 + f_6 m_7/16 \\ & + f_8 m_3/4 + f_8 m_4/8 + f_8 m_6/2 + f_8 m_7/4 \\ & + f_9 m_3/8 + f_9 m_4/16 + f_9 m_6/4 + f_9 m_7/8 \end{aligned}$ |
| $Y^M/Y^M; \text{III}/\text{III}^M;$<br>$\text{IV}/\text{IV}$ | $m_7$ | $\begin{aligned} & f_5 m_4/16 + f_5 m_5/8 + f_5 m_7/8 + f_5 m_8/4 \\ & + f_6 m_3/16 + f_6 m_4/16 + f_6 m_5/16 \\ & + f_6 m_6/8 + f_6 m_7/8 + f_6 m_8/8 \\ & + f_7 m_3/8 + f_7 m_4/16 + f_7 m_6/4 + f_7 m_7/8 \\ & + f_8 m_4/8 + f_8 m_5/4 + f_8 m_7/4 + f_8 m_8/2 \\ & + f_9 m_3/8 + f_9 m_4/8 + f_9 m_5/8 \\ & + f_9 m_6/4 + f_9 m_7/4 + f_9 m_8/4 \\ & + f_{10} m_3/4 + f_{10} m_4/8 + f_{10} m_6/2 + f_{10} m_7/4 \end{aligned}$ |
| $Y^M/Y^M; \text{III}^M/\text{III}^M;$<br>$\text{IV}/\text{IV}$ | $m_8$ | $\begin{aligned} & f_6 m_4/32 + f_6 m_5/16 + f_6 m_7/16 + f_6 m_8/8 \\ & + f_7 m_4/16 + f_7 m_5/8 + f_7 m_7/8 + f_7 m_8/4 \\ & + f_9 m_4/16 + f_9 m_5/8 + f_9 m_7/8 + f_9 m_8/4 \\ & + f_{10} m_4/8 + f_{10} m_5/4 + f_{10} m_7/4 + f_{10} m_8/2 \end{aligned}$ |

## Supplemental Methods

### Fitness effects and genotypic fitness

To generate the fitness of the 18 possible multi-chromosome genotypes (Table 1), I first assigned fitness effects to each single chromosome genotype (e.g., for the X and Y^M^ chromosomes there are three possible genotypes: X/X, X/Y^M^, and Y^M^/Y^M^). The single chromosome genotype is analogous to a single locus genotype in other population genetic models (e.g., Kidwell *et al*. 1977). I generated the single chromosome genotype fitness values by assigning a sex-specific fitness effect, *s*_*ij*_, to each *j* proto-Y or proto-W chromosome (Y^M^, III^M^, or IV^F^) in each sex *i*. The fitness effect of IV^F^ was only assigned for females because the female-determining proto-W chromosome (carrying *Md-tra^D^*) cannot be found in male genotypes (Table 1). Thus, there are five fitness effects: Y^M^ in males (*i*=male, *j*=Y^M^), Y^M^ in females (*i*=female, *j*=Y^M^), III^M^ in males (*i*=male, *j*=III^M^), III^M^ in females (*i*=female, *j*=III^M^), and IV^F^ in females (*i*=female, *j*=IV^F^). Each was drawn from a uniform distribution between -1 and 1. Positive fitness effects (*s_ij_*>0) mean that the proto-Y or proto-W chromosome increases fitness (i.e., it is beneficial), and negative values (*s_ij_*<0) mean that it decreases fitness (i.e., it is deleterious).

The fitness effects of the proto-Y and proto-W chromosome were then used to calculate the fitness of each single chromosome genotype, using one of four dominance scenarios. The calculations depend on whether the proto-Y chromosome has beneficial (*s_ij_*>0) or deleterious (*s_ij_*>0) effects, which ensures that the maximum genotypic fitness for a given chromosome and sex is equal to 1 (Table 2). First, in the additive model, the sex-specific single chromosome fitness values are defined such that the heterozygous genotype (X/Y^M^ or III/III^M^) is intermediate between the two homozygous genotypes. Second, when the proto-Y chromosomes have dominant fitness effects, the single chromosome fitness of the heterozygote is equal to the proto-Y homozygote. Third, when the effects of the proto-Y chromosomes are recessive, the single chromosome fitness of the heterozygote is equal to the proto-X homozygote and different from the proto-Y homozygote (Table 2). Fourth, the Y^M^ and III^M^ proto-Y chromosomes have overdominant effects in males, but are additive in females. In the overdominant model, the fitness effect of recessive deleterious proto-Y alleles is equal in magnitude to the fitness benefit of dominant advantageous proto-Y alleles. This simplifying assumption may not be biologically realistic, and future work could examine the implications of this assumption. In all four models, there is a single fitness effect associated with carrying a copy of the IV^F^ proto-W chromosome in females; only females can carry IV^F^ and it is impossible to be homozygous for IV^F^ (Table 1). In all four dominance models, the fitness of each of the multi-chromosomal genotypes (Table 1) is calculated as the product of the three relevant sex-specific single chromosome genotype fitness values.

I performed a normalization to ensure that the the maximum multi-chromosome genotype fitness in each sex is equal to 1. This is necessary because not all multi-chromosome genotypes are possible for both males and females, which means that the maximum product of all single chromosome genotypes can be <1 even though the maximum single chromosome genotype fitness values are set to 1. To ensure that the maximum female multi-chromosome genotype is 1, I divided each female multi-chromosome genotype fitness by the maximum female multi-chromosome genotype fitness. I did the same for males.

All four dominance scenarios (additive, dominant, recessive, or overdominant) attribute a single sex-specific fitness effect to each proto-Y and proto-W chromosome, whether it is Y^M^, III^M^, or IV^F^. I am therefore assuming that all copies of each proto-sex chromosome in the population carry an identical suite of beneficial and deleterious alleles (i.e., all copies of III^M^ are identical, all copies of Y^M^ are identical, and all copies of IV^F^ are identical). In addition, all models assume complete linkage between any allele(s) under selection and the *Mdmd* locus on the Y^M^ and III^M^ chromosomes. Similarly, I assume complete linkage between the female-determining *Md-tra^D^* locus and any alleles on the IV^F^ chromosome. Complete linkage may be achieved if, for example, chromosomal inversions suppress recombination between the proto-Y and proto-X (or proto-Z and proto-W) chromosomes in heterozygotes (Bergero and Charlesworth 2009; Wright *et al*. 2016). Free recombination in sex chromosome homozygotes would not affect genetic variation on a sex chromosome because all copies of each chromosome are assumed to carry the same alleles.

### Simulations with infinite population size

I used forward simulations to determine how the fitness effects of the proto-sex chromosomes affect their frequencies in populations. These simulations were performed with non-overlapping generations and random mating, assuming a population of infinite size (simulations with finite population sizes are described later). Each generation of a simulation consists of two discrete steps (Figure 1A). First, the frequency of each of the 18 genotypes is multiplied by its corresponding fitness value, which models differential survival across genotypes. Second, recursion equations developed by Hamm (2008) and previously implemented by Meisel *et al*. (2016) are used to model the production of progeny by random mating (Supplemental Table S1). After 1,000 generations of selection and random mating, the frequency of each genotype and proto-sex chromosome was calculated. This process was performed for 1,000,000 fitness arrays for each dominance scenario (additive, dominant, recessive, and overdominant) and four possible initial genotype frequencies (16,000,000 total simulations).

I considered four possible initial genotype frequencies for each simulation. The first three initial frequencies are based on the observed frequencies of Y^M^, III^M^, and *Md-tra^D^* from three North American populations (Meisel *et al*. 2016). These populations were sampled in Chino, CA (Meisel *et al*. 2016), Wake County, NC (Hamm and Scott 2008), and Chemung County, NY (Scott *et al*. 2013). Initial genotype frequencies were calculated based on the observed frequencies of Y^M^, III^M^, and *Md-tra^D^*, assuming random mating (Meisel *et al*. 2016). These three populations were chosen because Y^M^, III^M^, and *Md-tra^D^* have all remained at a frequency >1% across multiple years of sampling (Hamm *et al*. 2005; Hamm and Scott 2008; Meisel *et al*. 2016). I used frequencies from actual populations as the initial frequencies because I am specifically interested in how selection can maintain PSD at the frequencies observed in natural populations. The fourth starting values consist of all 18 genotypes at the same initial frequency (i.e., 1/18 for each genotype). This allows me to evaluate how selection pressures can drive the proto-sex chromosomes to the frequencies observed in natural populations if they start at arbitrary values that deviate from the observed frequencies.

To determine whether the fitness effects maintain PSD, I first selected fitness arrays that produced frequencies of each proto-sex chromosome (Y^M^, III^M^, IV^F^, and their homologous chromosomes, X, III, and IV) that are all >0.1% after 1,000 generations. This criterion ensures that all three chromosomes are polymorphic with both alleles at a frequency that is measurable (i.e., 1/1,000) given the sampling schemes used in previous studies of natural house fly populations (Hamm *et al*. 2005; Hamm and Scott 2008; Meisel *et al*. 2016). I refer to genotype fitness arrays with all proto-sex chromosomes present at a frequency >0.1% as maintaining PSD, although not necessarily at the frequencies observed in natural populations.

From the genotype fitness arrays that maintain PSD (i.e., all proto-sex chromosomes >0.1%), I selected those that produce proto-sex chromosome frequencies most similar to the frequencies observed in natural populations. To do so, I calculated the mean squared error (MSE) between the frequencies of Y^M^, III^M^, and IV^F^ after 1,000 generations in a simulation (*S*) and the observed frequencies (*O*) in a natural population (CA, NC, or NY):

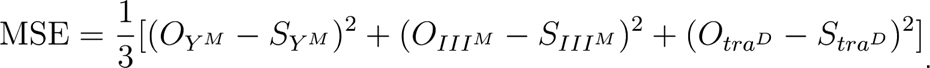

When using simulations that started with proto-sex chromosome frequencies observed in a population (e.g., CA), I only used MSE to compare with observed frequencies from that same population. When simulations started with equal frequencies of all 18 genotypes, I compared simulated chromosomes frequencies with the observed frequencies in each of the three populations (CA, NC, and NY).

I retained the 1,000 fitness arrays that produce proto-sex chromsome frequencies with the lowest MSE for a given dominance model, population, and starting genotype frequency. I refer to each of these 1,000 fitness arrays as the best-fitting arrays for each population. Simulations for the 1,000 best-fitting arrays were then run for 1,000,000 generations and compared to the 1,000 generation simulations to assess the stability of chromosomal frequencies. I also selected 1,000 random fitness arrays and 1,000 fitness arrays that maintain PSD to perform 1,000,000 generation simulations.

### Simulations with finite population sizes

I assessed how well the 1,000 best-fitting fitness arrays for each population, starting frequency, and dominance model maintain PSD when population sizes are finite. Finite population sizes introduce stochasticity to the change in allele frequencies across generations (i.e., genetic drift), in contrast to the deterministic effects of constant fitness values in a population of infinite size (Hartl and Clark 2007). It is not possible to calculate the probability of fixation for alleles in a finite population when there is complex sex-linked inheritance, as in the house fly PSD system (Meisel *et al*. 2016). To overcome this limitation, I estimated the probability of fixation using simulations that modeled finite populations by including multinomial sampling each generation.

I simulated finite populations by adding one additional step to the simulations described above in order to model a population size (*N*) of 10,000 individuals (Figure 1B). This population size was chosen because it is small enough to capture the effects of drift within the time scale of my simulations. Each simulation was started with the frequencies of the proto-sex chromosomes observed in the population where the fitness array was identified as best-fitting. In each generation, following multiplication by the fitness array (i.e., natural selection), I used multinomial sampling to calculate the new frequencies of all 18 genotypes. Multinomial sampling for all genotype was performed with 10,000 trials (i.e., *N*=10^4^) and a probability of success equal to the frequency of each genotype after selection. The resulting array of 18 genotype frequencies was then divided by the sum of all values to ensure that the genotype frequencies sum to 1. This array of genotype frequencies (after selection and multinomial sampling) was next used as input into the same recursion equations described above to calculate genotype frequencies after random mating (Supplemental Table S1). The process (selection, multinomial sampling, and random mating) was repeated for 1,000 generations. I simulated 100 replicate finite populations for each fitness array. I then used the frequency with which fixation and loss occur in those 100 replicates as an estimate of the probability of fixation and loss of proto-sex chromosomes within 1,000 generations.

I determined a null expectation for fixation or loss of proto-sex chromosomes in finite populations by using simulations without selection (i.e., genetic drift and no fitness differences across genotypes). To those ends, I performed simulations with 10,000 individuals for 1,000 generations, as was done in the simulations with selection and drift. I performed 1,000 replicate “drift-only” simulations with starting values at the observed frequencies of the proto-sex chromosomes in each population (CA, NC, and NY). These drift-only simulations included the same steps as the simulations with both selection and drift, except that there is no differential survival (i.e., natural selection) step in the drift-only simulation. After 1,000 generations, I calculated the frequency of each proto-Y and proto-W chromosome (Y^M^, III^M^, and IV^F^), as well as the frequency with which each chromosome was lost, across the 1,000 simulations.

