## Supplementary material for "The maintenance of polygenic sex determination depends on the dominance of fitness effects which are predictive of the role of sexual antagonism": Code: drift_allele_pub.pl

```
#!/usr/bin/perl

# Run recursions in a finite population with no selection
# Start at observed genotype frequencies to see if it will hold. 


# cd ~/Projects/Calyptrates/Musca/PopGenModel2020/
# perl drift_allele_pub.pl generations popsize reps controlfile output
# perl drift_allele_pub.pl 1000 10000 1000 CA_femHW.txt drift_CA.out
# perl drift_allele_pub.pl 1000 10000 1000 NC_femHW.txt drift_NC.out
# perl drift_allele_pub.pl 1000 10000 1000 NY_femHW.txt drift_NY.out


use strict;
use warnings;
use List::Util qw(sum min max);
use POSIX; # for rounding
use Math::Random qw(random_binomial); # for random number generation: random_binomial($n, $nt, $p)
	# $n is number of outcomes
	# $nt is number of trials
	# $p is probability of success


my $popsize=10000; # population size (can be reset by user)
my $generations=1000; # number of generations to run each simulation (can be reset by user)
my $reps=100; # how many replicate simulations (can be reset by user)
my $controlfile;
my $outfile;
my @initial_freq;


($generations, $popsize, $reps, $controlfile, $outfile) = @ARGV;


# Make sure generations are >0
if($generations<1){
	die "\nGenerations must be >0\n\n";
}

# Make sure population size is >0
if($popsize<1){
	die "\nPopulation size must be >0\n\n";
}

# Make sure reps are >0
if($reps<1){
	die "\nReplicates must be >0\n\n";
}

# 0.332142857 			0.034314191 		0.004289274 			0.000134040 			0.074230699 		0.009278837 			0.000289964 			0.040145174 		0.005018147 		0.039215686 			0.000000000 			0.382352941 		0.019607843 			0.000000000 			0.058823529 		0.000000000 			0.000000000 			0.000156817

# obtain initial genotype frequencies for simulation
open INPUT, "$controlfile" or die "\nCould not open control file, $controlfile\n";
my $counter=0;
while(<INPUT>){
	if($counter==0){
		chomp;
		@initial_freq = split /\s+/, $_; # delimit each genotype frequency by white space
	}else{
		last;
	}
	$counter++;
}
close INPUT;


# 1 female f1 M0 F0 Y0	III /III	tra+/tra+	X/X
# 2 female f2 M0 F1 Y0	III /III	traD/tra+	X/X
# 3 female f3 M1 F1 Y0	IIIM/III	traD/tra+	X/X
# 4 female f4 M2 F1 Y0	IIIM/IIIM	traD/tra+	X/X
# 5 female f5 M0 F1 Y1	III /III	traD/tra+	X/Y
# 6 female f6 M1 F1 Y1	IIIM/III	traD/tra+	X/Y
# 7 female f7 M2 F1 Y1	IIIM/IIIM	traD/tra+	X/Y
# 8 female f8 M0 F1 Y2	III /III	traD/tra+	Y/Y
# 9 female f9 M1 F1 Y2	IIIM/III	traD/tra+	Y/Y
#10 male   m1 M1 F0 Y0	IIIM/III	tra+/tra+	X/X
#11 male   m2 M2 F0 Y0	IIIM/IIIM	tra+/tra+	X/X
#12 male   m3 M0 F0 Y1	III /III	tra+/tra+	Y/X
#13 male   m4 M1 F0 Y1	IIIM/III	tra+/tra+	Y/X
#14 male   m5 M2 F0 Y1	IIIM/IIIM	tra+/tra+	Y/X
#15 male   m6 M0 F0 Y2	III /III	tra+/tra+	Y/Y
#16 male   m7 M1 F0 Y2	IIIM/III	tra+/tra+	Y/Y
#17 male   m8 M2 F0 Y2	IIIM/IIIM	tra+/tra+	Y/Y
#18 female m9 M2 F1 Y2	IIIM/IIIM	traD/tra+	Y/Y <- should be f10


open OUTPUT, ">$outfile" or die "\nCould not open output file, $outfile\n";
print OUTPUT "run\t";
print OUTPUT "f1\tf2\tf3\tf4\tf5\tf6\tf7\tf8\tf9\t";
print OUTPUT "m1\tm2\tm3\tm4\tm5\tm6\tm7\tm8\tm9\t";
print OUTPUT "female\tmale\t";
print OUTPUT "X\tY\tIII\tIIIM\ttra\ttraD\n";

# Run simulations
for(my $i=1; $i <= $reps; $i++){
					
	my @simulation_results;
	$simulation_results[0] = [ @initial_freq ];
	for(my $a=1; $a <= $generations; $a++){
		$simulation_results[$a] = [ recursion_drift(\@{$simulation_results[$a-1]}, $popsize) ]; # need to pass the function references to the array, not the arrays themselves
	}

	# Print final results of simulation to file
	print OUTPUT "$i\t";

	# Final genotype frequencies
	for(my $b=0; $b<18; $b++){
		print OUTPUT "$simulation_results[$generations][$b]\t";
	}
	
	# Final sex ratio
	my @sex_ratios = sex_ratio(@{$simulation_results[$generations]});
	print OUTPUT "$sex_ratios[0]\t$sex_ratios[1]";
	
	# Final allele frequencies
	my @allele_freq = allele_freqs(@{$simulation_results[$generations]});
	for(my $b=0; $b<6; $b++){
		print OUTPUT "\t$allele_freq[$b]";
	}
	print OUTPUT "\n";	
	
}
close OUTPUT;


#print "\n$counter parameterizations read from $controlfile and printed to $counter outfiles named $outfile\n\n";


# Function performs 1 generation of genetic drift (binomial sampling) and random mating
# Send function one array and one scalar:
#	array: counts of the 18 genotypes
#	scalar: population size
sub recursion_drift {
	my($raw_counts_ref, $popnsize) = @_; # pass the function reference to the genotype frequency array
	
	# Dereference into arrays
	my @raw_counts = @{$raw_counts_ref}; 
	
	# binomial sampling for genetic drift
	# random_binomial($n, $nt, $p)
		# $n is number of outcomes
		# $nt is number of trials
		# $p is probability of success
	my @drift_counts;
	for(my $i=0; $i<18; $i++){
		$drift_counts[$i] = random_binomial(1, $popnsize, $raw_counts[$i]) / $popnsize;
	}
	
	# normalize back to sum to 1
	my @counts;
	for (my $i=0; $i<18; $i++) {
		$counts[$i] = $drift_counts[$i] / sum(@drift_counts);
	}

	
	# Calculate genotype frequencies after random mating
	my $f1 = $counts[0];
	my $f2 = $counts[1];
	my $f3 = $counts[2];
	my $f4 = $counts[3];
	my $f5 = $counts[4];
	my $f6 = $counts[5];
	my $f7 = $counts[6];
	my $f8 = $counts[7];
	my $f9 = $counts[8];

	my $m1 = $counts[9];
	my $m2 = $counts[10];
	my $m3 = $counts[11];
	my $m4 = $counts[12];
	my $m5 = $counts[13];
	my $m6 = $counts[14];
	my $m7 = $counts[15];
	my $m8 = $counts[16];
	my $m9 = $counts[17];
	
	my @after_mating = (
		$f1*$m1*.5 + $f1*$m3*.5 + $f1*$m4*.25 + $f2*$m1*.25 + $f2*$m3*.25 + $f2*$m4*.125 + $f3*$m1*.25 + $f3*$m3*.125 + $f3*$m4*.0625  + $f5*$m1*.125 + $f5*$m3*.125 + $f5*$m4*.0625 + $f6*$m1*.0625 + $f6*$m3*.0625 + $f6*$m4*.03125,
		$f2*$m1*.25 + $f2*$m3*.25 + $f2*$m4*.125 + $f3*$m1*.125 + $f3*$m3*.125 + $f3*$m4*.0625 + $f5*$m1*.125 + $f5*$m3*.125 + $f5*$m4*.0625 + $f6*$m1*.0625 + $f6*$m3*.0625 + $f6*$m4*.03125,
		$f2*$m1*.25 + $f2*$m2*.5 + $f2*$m4*.125 + $f2*$m5*.25 + $f3*$m1*.25 + $f3*$m2*.25 + $f3*$m3*.125 + $f3*$m4*.125 + $f3*$m5*.125 + $f4*$m1*.25 + $f4*$m3*.25 + $f4*$m4*.125 + $f5*$m1*.125 + $f5*$m2*.25 + $f5*$m4*.0625 + $f5*$m5*.125 + $f6*$m1*.125 + $f6*$m2*.125 + $f6*$m3*.0625 + $f6*$m4*.0625 + $f6*$m5*.0625 + $f7*$m1*.125 + $f7*$m3*.125 + $f7*$m4*.0625,
		$f3*$m1*.125 + $f3*$m2*.25 + $f3*$m4*.0625 + $f3*$m5*.125 + $f4*$m1*.25 + $f4*$m2*.5 + $f4*$m4*.125 + $f4*$m5*.25 + $f6*$m1*.0625 + $f6*$m2*.125 + $f6*$m4*.03125 + $f6*$m5*.0625 + $f7*$m1*.125 + $f7*$m2*.25 + $f7*$m4*.0625 + $f7*$m5*.125,
		$f2*$m3*.25 + $f2*$m4*.125 + $f2*$m6*.5 + $f2*$m7*.25 + $f3*$m3*.125 + $f3*$m4*.0625 + $f3*$m6*.25 + $f3*$m7*.125 + $f5*$m1*.125 + $f5*$m3*.25 + $f5*$m4*.125 + $f5*$m6*.25 + $f5*$m7*.125 + $f6*$m1*.0625 + $f6*$m3*.125 + $f6*$m4*.0625 + $f6*$m6*.125 + $f6*$m7*.0625 + $f8*$m1*.25 + $f8*$m3*.25 + $f8*$m4*.125 + $f9*$m1*.125 + $f9*$m3*.125 + $f9*$m4*.0625,
		$f2*$m4*.125 + $f2*$m5*.25 + $f2*$m7*.25 + $f2*$m8*.5 + $f3*$m3*.125 + $f3*$m4*.125 + $f3*$m5*.125 + $f3*$m6*.25 + $f3*$m7*.25 + $f3*$m8*.25 + $f4*$m3*.25 + $f4*$m4*.125 + $f4*$m6*.5 + $f4*$m7*.25 + $f5*$m1*.125 + $f5*$m2*.25 + $f5*$m4*.125 + $f5*$m5*.25 + $f5*$m7*.125 + $f5*$m8*.25 + $f6*$m1*.125 + $f6*$m2*.125 + $f6*$m3*.125 + $f6*$m4*.125 + $f6*$m5*.125 + $f6*$m6*.125 + $f6*$m7*.125 + $f6*$m8*.125 + $f7*$m1*.125 + $f7*$m3*.25 + $f7*$m4*.125 + $f7*$m6*.25 + $f7*$m7*.125 + $f8*$m1*.25 + $f8*$m2*.5 + $f8*$m4*.125 + $f8*$m5*.25 + $f9*$m1*.25 + $f9*$m2*.25 + $f9*$m3*.125 + $f9*$m4*.125 + $f9*$m5*.125 + $m9*$m1*.25 + $m9*$m3*.25 + $m9*$m4*.125,
		$f3*$m4*.0625 + $f3*$m5*.125 + $f3*$m7*.125 + $f3*$m8*.25 + $f4*$m4*.125 + $f4*$m5*.25 + $f4*$m7*.25 + $f4*$m8*.5 + $f6*$m1*.0625 + $f6*$m2*.125 + $f6*$m4*.0625 + $f6*$m5*.125 + $f6*$m7*.0625 + $f6*$m8*.125 + $f7*$m1*.125 + $f7*$m2*.25 + $f7*$m4*.125 + $f7*$m5*.25 + $f7*$m7*.125 + $f7*$m8*.25 + $f9*$m1*.125 + $f9*$m2*.25 + $f9*$m4*.0625 + $f9*$m5*.125 + $m9*$m1*.25 + $m9*$m2*.5 + $m9*$m4*.125 + $m9*$m5*.25,
		$f5*$m3*.125 + $f5*$m4*.0625 + $f5*$m6*.25 + $f5*$m7*.125 + $f6*$m3*.0625 + $f6*$m4*.03125 + $f6*$m6*.125 + $f6*$m7*.0625 + $f8*$m3*.25 + $f8*$m4*.125 + $f8*$m6*.5 + $f8*$m7*.25 + $f9*$m3*.125 + $f9*$m4*.0625 + $f9*$m6*.25 + $f9*$m7*.125,
		$f5*$m4*.0625 + $f5*$m5*.125 + $f5*$m7*.125 + $f5*$m8*.25 + $f6*$m3*.0625 + $f6*$m4*.0625 + $f6*$m5*.0625 + $f6*$m6*.125 + $f6*$m7*.125 + $f6*$m8*.125 + $f7*$m3*.125 + $f7*$m4*.0625 + $f7*$m6*.25 + $f7*$m7*.125 + $f8*$m4*.125 + $f8*$m5*.25 + $f8*$m7*.25 + $f8*$m8*.5 + $f9*$m3*.125 + $f9*$m4*.125 + $f9*$m5*.125 + $f9*$m6*.25 + $f9*$m7*.25 + $f9*$m8*.25 + $m9*$m3*.25 + $m9*$m4*.125 + $m9*$m6*.5 + $m9*$m7*.25,

		$f1*$m1*.5 + $f1*$m2*1 + $f1*$m4*.25 + $f1*$m5*.5 + $f2*$m1*.25 + $f2*$m2*.5 + $f2*$m4*.125 + $f2*$m5*.25 + $f3*$m1*.125 + $f3*$m2*.25 + $f3*$m3*.125 + $f3*$m4*.125 + $f3*$m5*.125 + $f4*$m1*.25 + $f4*$m3*.25 + $f4*$m4*.125 + $f5*$m1*.125 + $f5*$m2*.25 + $f5*$m4*.0625 + $f5*$m5*.125 + $f6*$m1*.125 + $f6*$m2*.125 + $f6*$m3*.0625 + $f6*$m4*.0625 + $f6*$m5*.0625 + $f7*$m1*.125 + $f7*$m3*.125 + $f7*$m4*.0625,
		$f3*$m1*.125 + $f3*$m2*.25 + $f3*$m4*.0625 + $f3*$m5*.125 + $f4*$m1*.25 + $f4*$m2*.5 + $f4*$m4*.125 + $f4*$m5*.25 + $f6*$m1*.0625 + $f6*$m2*.125 + $f6*$m4*.03125 + $f6*$m5*.0625 + $f7*$m1*.125 + $f7*$m2*.25 + $f7*$m4*.0625 + $f7*$m5*.125,
		$f1*$m3*.5 + $f1*$m4*.25 + $f1*$m6*1 + $f1*$m7*.5 + $f2*$m3*.25 + $f2*$m4*.125 + $f2*$m6*.5 + $f2*$m7*.25 + $f3*$m3*.125 + $f3*$m4*.0625 + $f3*$m6*.25 + $f3*$m7*.125 + $f5*$m1*.125 + $f5*$m3*.25 + $f5*$m4*.125 + $f5*$m6*.25 + $f5*$m7*.125 + $f6*$m1*.0625 + $f6*$m3*.125 + $f6*$m4*.0625 + $f6*$m6*.125 + $f6*$m7*.0625 + $f8*$m1*.25 + $f8*$m3*.25 + $f8*$m4*.125 + $f9*$m1*.125 + $f9*$m3*.125 + $f9*$m4*.0625,
		$f1*$m4*.25 + $f1*$m5*.5 + $f1*$m7*.5 + $f1*$m8*1 + $f2*$m4*.125 + $f2*$m5*.25 + $f2*$m7*.25 + $f2*$m8*.5 + $f3*$m3*.125 + $f3*$m4*.125 + $f3*$m5*.125 + $f3*$m6*.25 + $f3*$m7*.25 + $f3*$m8*.25 + $f4*$m3*.25 + $f4*$m4*.125 + $f4*$m6*.5 + $f4*$m7*.25 + $f5*$m1*.125 + $f5*$m2*.25 + $f5*$m4*.125 + $f5*$m5*.25 + $f5*$m7*.125 + $f5*$m8*.25 + $f6*$m1*.125 + $f6*$m2*.125 + $f6*$m3*.125 + $f6*$m4*.125 + $f6*$m5*.125 + $f6*$m6*.125 + $f6*$m7*.125 + $f6*$m8*.125 + $f7*$m1*.125 + $f7*$m3*.25 + $f7*$m4*.125 + $f7*$m6*.25 + $f7*$m7*.125 + $f8*$m1*.25 + $f8*$m2*.5 + $f8*$m4*.125 + $f8*$m5*.25 + $f9*$m1*.25 + $f9*$m2*.25 + $f9*$m3*.125 + $f9*$m4*.125 + $f9*$m5*.125 + $m9*$m1*.25 + $m9*$m3*.25 + $m9*$m4*.125,
		$f3*$m4*.0625 + $f3*$m5*.125 + $f3*$m7*.125 + $f3*$m8*.25 + $f4*$m4*.125 + $f4*$m5*.25 + $f4*$m7*.25 + $f4*$m8*.5 + $f6*$m1*.0625 + $f6*$m2*.125 + $f6*$m4*.0625 + $f6*$m5*.125 + $f6*$m7*.0625 + $f6*$m8*.125 + $f7*$m1*.125 + $f7*$m2*.25 + $f7*$m4*.125 + $f7*$m5*.25 + $f7*$m7*.125 + $f7*$m8*.25 + $f9*$m1*.125 + $f9*$m2*.25 + $f9*$m4*.0625 + $f9*$m5*.125 + $m9*$m1*.25 + $m9*$m2*.5 + $m9*$m4*.125 + $m9*$m5*.25,
		$f5*$m3*.125 + $f5*$m4*.0625 + $f5*$m6*.25 + $f5*$m7*.125 + $f6*$m3*.0625 + $f6*$m4*.03125 + $f6*$m6*.125 + $f6*$m7*.0625 + $f8*$m3*.25 + $f8*$m4*.125 + $f8*$m6*.5 + $f8*$m7*.25 + $f9*$m3*.125 + $f9*$m4*.0625 + $f9*$m6*.25 + $f9*$m7*.125,
		$f5*$m4*.0625 + $f5*$m5*.125 + $f5*$m7*.125 + $f5*$m8*.25 + $f6*$m3*.0625 + $f6*$m4*.0625 + $f6*$m5*.0625 + $f6*$m6*.125 + $f6*$m7*.125 + $f6*$m8*.125 + $f7*$m3*.125 + $f7*$m4*.0625 + $f7*$m6*.25 + $f7*$m7*.125 + $f8*$m4*.125 + $f8*$m5*.25 + $f8*$m7*.25 + $f8*$m8*.5 + $f9*$m3*.125 + $f9*$m4*.125 + $f9*$m5*.125 + $f9*$m6*.25 + $f9*$m7*.25 + $f9*$m8*.25 + $m9*$m3*.25 + $m9*$m4*.125 + $m9*$m6*.5 + $m9*$m7*.25,
		$f6*$m4*.03125 + $f6*$m5*.0625 + $f6*$m7*.0625 + $f6*$m8*.125 + $f7*$m4*.0625 + $f7*$m5*.125 + $f7*$m7*.125 + $f7*$m8*.25 + $f9*$m4*.0625 + $f9*$m5*.125 + $f9*$m7*.125 + $f9*$m8*.25 + $m9*$m4*.125 + $m9*$m5*.25 + $m9*$m7*.25 + $m9*$m8*.5,
		$f6*$m4*.03125 + $f6*$m5*.0625 + $f6*$m7*.0625 + $f6*$m8*.125 + $f7*$m4*.0625 + $f7*$m5*.125 + $f7*$m7*.125 + $f7*$m8*.25 + $f9*$m4*.0625 + $f9*$m5*.125 + $f9*$m7*.125 + $f9*$m8*.25 + $m9*$m4*.125 + $m9*$m5*.25 + $m9*$m7*.25 + $m9*$m8*.5
	);
	
	# Calculate standardized genotype frequencies to sum to 1
	my @after_mating_std;
	for (my $i=0; $i<18; $i++) {
		$after_mating_std[$i] = $after_mating[$i] / sum(@after_mating);
	}

	return @after_mating_std;
}

# Function calculates the genotype frequencies 
# given an array of all possible genotypes
sub genotype_freqs{
	my @counts = @_;
	my @freqs;
	for(my $i=0; $i<18; $i++){
		$freqs[$i] = $counts[$i]/sum(@counts)
	}
	return @freqs;
}

# Function calculates the allele frequencies 
# given an array of all possible genotypes
sub allele_freqs{
	my @freqs = @_;
	
	my $X_freq = ($freqs[0] + $freqs[1] + $freqs[2] + $freqs[3] + $freqs[9] + $freqs[10]
				+ 0.5*($freqs[4] + $freqs[5] + $freqs[6] + $freqs[11] + $freqs[12] + $freqs[13])) / sum(@freqs);
	my $Y_freq = 1 - $X_freq;

	my $III_freq = ($freqs[0] + $freqs[1] + $freqs[4] + $freqs[7] + $freqs[11] + $freqs[14]
				+ 0.5*($freqs[2] + $freqs[5] + $freqs[8] + $freqs[9] + $freqs[12] + $freqs[15])) / sum(@freqs);
	my $IIIM_freq = 1 - $III_freq;
	
	my $tra_freq = ($freqs[0] + $freqs[9] + $freqs[10] + $freqs[11] + $freqs[12] + $freqs[13] + $freqs[14] + $freqs[15] + $freqs[16]
				+ 0.5*($freqs[1] + $freqs[2] + $freqs[3] + $freqs[4] + $freqs[5] + $freqs[6] + $freqs[7] + $freqs[8] + $freqs[17])) / sum(@freqs);
	my $traD_freq = 1 - $tra_freq;

	return ($X_freq, $Y_freq, $III_freq, $IIIM_freq, $tra_freq, $traD_freq);
}

# Function calculates sex ratio
# given an array of all possible genotypes
sub sex_ratio{
	my @freqs = @_;

	my $female = ($freqs[0] + $freqs[1] + $freqs[2] + $freqs[3] + $freqs[4] + $freqs[5] + $freqs[6] + $freqs[7] + $freqs[8] + $freqs[17])/sum(@freqs);
	my $male = ($freqs[9] + $freqs[10] + $freqs[11] + $freqs[12] + $freqs[13] + $freqs[14] + $freqs[15] + $freqs[16])/sum(@freqs);
	
	return($female, $male);
}
```
