## Supplementary material for "The maintenance of polygenic sex determination depends on the dominance of fitness effects which are predictive of the role of sexual antagonism": Code: seln_allele_additive_v2.pl

```
#!/usr/bin/perl
# Simulate populations where proto-Y chromosomes have ADDITIVE fitness effects 

# perl seln_allele_additive_v2.pl generations simulations control output
# perl seln_allele_additive_v2.pl 100 10 CA_femHW.txt seln_allele_additive_v2_CA.out
# perl seln_allele_additive_v2.pl 1000 100 CA_femHW.txt seln_allele_additive_CA.out
# perl seln_allele_additive_v2.pl 1000 100 NC_femHW.txt seln_allele_additive_NC.out
# perl seln_allele_additive_v2.pl 1000 100 NY_femHW.txt seln_allele_additive_NY.out
# perl seln_allele_additive_v2.pl 1000 100 equal_freq.txt seln_allele_additive_equal.out


# New version does the following: 
# assigns value of "s" (ranging between 0-1) for each proto-sex chromosome
# assume h=0.5


use strict;
use warnings;
use List::Util qw(sum min max);
use Math::Random qw(random_uniform); # for random number generation -- random_uniform($n, $low, $high)
use POSIX; # for rounding

my $simulations=10; # number of simulations to perform
my $generations=100; # number of generations to run each simulation (can be reset by user)
my $ctlfile; # contains initial genotype frequencies & target frequencies
my $outfile;
my @initial_freq = (0.05556, 0.05556, 0.05555, 0.05556, 0.05555, 0.05556, 0.05555, 0.05556, 0.05555, 0.05556, 0.05555, 0.05556, 0.05555, 0.05556, 0.05555, 0.05556, 0.05555, 0.05556); # default to equal frequencies
my @allele_fitness; # Contains selection coefficients for each allele in each sex
my @fitness_array; # Contains fitness values for each genotype
my @simulation_results;

# 1 female f1 M0 F0 Y0	III /III	tra+/tra+	X/X
# 2 female f2 M0 F1 Y0	III /III	traD/tra+	X/X
# 3 female f3 M1 F1 Y0	IIIM/III	traD/tra+	X/X
# 4 female f4 M2 F1 Y0	IIIM/IIIM	traD/tra+	X/X
# 5 female f5 M0 F1 Y1	III /III	traD/tra+	X/Y
# 6 female f6 M1 F1 Y1	IIIM/III	traD/tra+	X/Y
# 7 female f7 M2 F1 Y1	IIIM/IIIM	traD/tra+	X/Y
# 8 female f8 M0 F1 Y2	III /III	traD/tra+	Y/Y
# 9 female f9 M1 F1 Y2	IIIM/III	traD/tra+	Y/Y
#10 male   m1 M1 F0 Y0	IIIM/III	tra+/tra+	X/X
#11 male   m2 M2 F0 Y0	IIIM/IIIM	tra+/tra+	X/X
#12 male   m3 M0 F0 Y1	III /III	tra+/tra+	Y/X
#13 male   m4 M1 F0 Y1	IIIM/III	tra+/tra+	Y/X
#14 male   m5 M2 F0 Y1	IIIM/IIIM	tra+/tra+	Y/X
#15 male   m6 M0 F0 Y2	III /III	tra+/tra+	Y/Y
#16 male   m7 M1 F0 Y2	IIIM/III	tra+/tra+	Y/Y
#17 male   m8 M2 F0 Y2	IIIM/IIIM	tra+/tra+	Y/Y
#18 female m9 M2 F1 Y2	IIIM/IIIM	traD/tra+	Y/Y <- should be f10


($generations, $simulations, $ctlfile, $outfile) = @ARGV;

# Make sure number of simulations are > 0
if($simulations < 1){
	die "\nSimulations must be >0\n";
}

# Make sure generations are >0
if($generations<1){
	die "\nGenerations must be >0\n";
}


# obtain initial genotype frequencies for simulation
open INPUT, "$ctlfile" or die "\nCould not open control file, $ctlfile\n";
my $counter=0;
while(<INPUT>){
	if($counter==0){
		chomp;
		@initial_freq = split /\s+/, $_; # delimit each genotype frequency by white space
	}else{
		last;
	}
	$counter++;
}
close INPUT;


	#	1 IIIM 	female
	#	2 YM   	female
	#	3 traD	female
	#	4 IIIM	male
	#	5 YM   	male

# Run simulations
open OUTPUT, ">$outfile" or die "\nCould not open output file, $outfile\n";
print OUTPUT "IIIM_fem\tYM_fem\ttraD_fem\tIIIM_male\tYM_male\t";
print OUTPUT "f1_fit\tf2_fit\tf3_fit\tf4_fit\tf5_fit\tf6_fit\tf7_fit\tf8_fit\tf9_fit\t";
print OUTPUT "m1_fit\tm2_fit\tm3_fit\tm4_fit\tm5_fit\tm6_fit\tm7_fit\tm8_fit\tm9_fit\t";
print OUTPUT "f1\tf2\tf3\tf4\tf5\tf6\tf7\tf8\tf9\t";
print OUTPUT "m1\tm2\tm3\tm4\tm5\tm6\tm7\tm8\tm9\t";
print OUTPUT "X\tY\tIII\tIIIM\ttra\ttraD\t";
print OUTPUT "female\tmale\n";

for (my $i=0; $i<$simulations; $i++){
	# set allele fitness values
	@allele_fitness = random_uniform(5, -1, 1);
	
	# calculate genotype fitness values
	@fitness_array = genotype_fitness(@allele_fitness);

	# Run the population simulation
	my @simulation_results;
	$simulation_results[0] = [ @initial_freq ];
	for(my $a=1; $a <= $generations; $a++){
		$simulation_results[$a] = [ recursion_seln(\@{$simulation_results[$a-1]}, \@fitness_array) ]; # need to pass the function references to the arrays, not the arrays themselves
	}
	# Print final results of simulation to file
	my @allele_freq = allele_freqs(@{$simulation_results[$generations]});
	my @sex_ratios = sex_ratio(@{$simulation_results[$generations]});
	
	for(my $b=0; $b<5; $b++){
		print OUTPUT "$allele_fitness[$b]\t";
	}
	for(my $b=0; $b<18; $b++){
		print OUTPUT "$fitness_array[$b]\t";
	}
#	print OUTPUT "$a\t"; # generation
	for(my $b=0; $b<18; $b++){
		print OUTPUT "$simulation_results[$generations][$b]\t";
	}
	for(my $b=0; $b<6; $b++){
		print OUTPUT "$allele_freq[$b]\t";
	}
	print OUTPUT "$sex_ratios[0]\t$sex_ratios[1]\n";															
}
close OUTPUT;


# Function performs 1 generation of selection, followed by random mating
# Send function 2 arrays:
#	1. counts of the 18 genotypes
#	2. fitness values for the 18 genotypes
sub recursion_seln {
	my($raw_counts_ref, $fitness_ref) = @_; # pass the function references to the arrays
	
	# Dereference into arrays
	my @raw_counts = @{$raw_counts_ref}; 
	my @fitness = @{$fitness_ref};

	my @counts;
	
	# impose selection pressures
	for(my $i=0; $i<18; $i++) {
		$counts[$i] = $raw_counts[$i] * $fitness[$i];
	}

	# Calculate standardized genotype frequencies to sum to 1
	my @counts_norm;
	for (my $i=0; $i<18; $i++) {
		$counts_norm[$i] = $counts[$i] / sum(@counts);
	}	
	
	my $f1 = $counts_norm[0];
	my $f2 = $counts_norm[1];
	my $f3 = $counts_norm[2];
	my $f4 = $counts_norm[3];
	my $f5 = $counts_norm[4];
	my $f6 = $counts_norm[5];
	my $f7 = $counts_norm[6];
	my $f8 = $counts_norm[7];
	my $f9 = $counts_norm[8];
	my $m1 = $counts_norm[9];
	my $m2 = $counts_norm[10];
	my $m3 = $counts_norm[11];
	my $m4 = $counts_norm[12];
	my $m5 = $counts_norm[13];
	my $m6 = $counts_norm[14];
	my $m7 = $counts_norm[15];
	my $m8 = $counts_norm[16];
	my $m9 = $counts_norm[17]; # actually female
	
	# Calculate genotype frequencies after random mating
	my @after_mating = (
		$f1*$m1*.5 + $f1*$m3*.5 + $f1*$m4*.25 + $f2*$m1*.25 + $f2*$m3*.25 + $f2*$m4*.125 + $f3*$m1*.25 + $f3*$m3*.125 + $f3*$m4*.0625  + $f5*$m1*.125 + $f5*$m3*.125 + $f5*$m4*.0625 + $f6*$m1*.0625 + $f6*$m3*.0625 + $f6*$m4*.03125,
		$f2*$m1*.25 + $f2*$m3*.25 + $f2*$m4*.125 + $f3*$m1*.125 + $f3*$m3*.125 + $f3*$m4*.0625 + $f5*$m1*.125 + $f5*$m3*.125 + $f5*$m4*.0625 + $f6*$m1*.0625 + $f6*$m3*.0625 + $f6*$m4*.03125,
		$f2*$m1*.25 + $f2*$m2*.5 + $f2*$m4*.125 + $f2*$m5*.25 + $f3*$m1*.25 + $f3*$m2*.25 + $f3*$m3*.125 + $f3*$m4*.125 + $f3*$m5*.125 + $f4*$m1*.25 + $f4*$m3*.25 + $f4*$m4*.125 + $f5*$m1*.125 + $f5*$m2*.25 + $f5*$m4*.0625 + $f5*$m5*.125 + $f6*$m1*.125 + $f6*$m2*.125 + $f6*$m3*.0625 + $f6*$m4*.0625 + $f6*$m5*.0625 + $f7*$m1*.125 + $f7*$m3*.125 + $f7*$m4*.0625,
		$f3*$m1*.125 + $f3*$m2*.25 + $f3*$m4*.0625 + $f3*$m5*.125 + $f4*$m1*.25 + $f4*$m2*.5 + $f4*$m4*.125 + $f4*$m5*.25 + $f6*$m1*.0625 + $f6*$m2*.125 + $f6*$m4*.03125 + $f6*$m5*.0625 + $f7*$m1*.125 + $f7*$m2*.25 + $f7*$m4*.0625 + $f7*$m5*.125,
		$f2*$m3*.25 + $f2*$m4*.125 + $f2*$m6*.5 + $f2*$m7*.25 + $f3*$m3*.125 + $f3*$m4*.0625 + $f3*$m6*.25 + $f3*$m7*.125 + $f5*$m1*.125 + $f5*$m3*.25 + $f5*$m4*.125 + $f5*$m6*.25 + $f5*$m7*.125 + $f6*$m1*.0625 + $f6*$m3*.125 + $f6*$m4*.0625 + $f6*$m6*.125 + $f6*$m7*.0625 + $f8*$m1*.25 + $f8*$m3*.25 + $f8*$m4*.125 + $f9*$m1*.125 + $f9*$m3*.125 + $f9*$m4*.0625,
		$f2*$m4*.125 + $f2*$m5*.25 + $f2*$m7*.25 + $f2*$m8*.5 + $f3*$m3*.125 + $f3*$m4*.125 + $f3*$m5*.125 + $f3*$m6*.25 + $f3*$m7*.25 + $f3*$m8*.25 + $f4*$m3*.25 + $f4*$m4*.125 + $f4*$m6*.5 + $f4*$m7*.25 + $f5*$m1*.125 + $f5*$m2*.25 + $f5*$m4*.125 + $f5*$m5*.25 + $f5*$m7*.125 + $f5*$m8*.25 + $f6*$m1*.125 + $f6*$m2*.125 + $f6*$m3*.125 + $f6*$m4*.125 + $f6*$m5*.125 + $f6*$m6*.125 + $f6*$m7*.125 + $f6*$m8*.125 + $f7*$m1*.125 + $f7*$m3*.25 + $f7*$m4*.125 + $f7*$m6*.25 + $f7*$m7*.125 + $f8*$m1*.25 + $f8*$m2*.5 + $f8*$m4*.125 + $f8*$m5*.25 + $f9*$m1*.25 + $f9*$m2*.25 + $f9*$m3*.125 + $f9*$m4*.125 + $f9*$m5*.125 + $m9*$m1*.25 + $m9*$m3*.25 + $m9*$m4*.125,
		$f3*$m4*.0625 + $f3*$m5*.125 + $f3*$m7*.125 + $f3*$m8*.25 + $f4*$m4*.125 + $f4*$m5*.25 + $f4*$m7*.25 + $f4*$m8*.5 + $f6*$m1*.0625 + $f6*$m2*.125 + $f6*$m4*.0625 + $f6*$m5*.125 + $f6*$m7*.0625 + $f6*$m8*.125 + $f7*$m1*.125 + $f7*$m2*.25 + $f7*$m4*.125 + $f7*$m5*.25 + $f7*$m7*.125 + $f7*$m8*.25 + $f9*$m1*.125 + $f9*$m2*.25 + $f9*$m4*.0625 + $f9*$m5*.125 + $m9*$m1*.25 + $m9*$m2*.5 + $m9*$m4*.125 + $m9*$m5*.25,
		$f5*$m3*.125 + $f5*$m4*.0625 + $f5*$m6*.25 + $f5*$m7*.125 + $f6*$m3*.0625 + $f6*$m4*.03125 + $f6*$m6*.125 + $f6*$m7*.0625 + $f8*$m3*.25 + $f8*$m4*.125 + $f8*$m6*.5 + $f8*$m7*.25 + $f9*$m3*.125 + $f9*$m4*.0625 + $f9*$m6*.25 + $f9*$m7*.125,
		$f5*$m4*.0625 + $f5*$m5*.125 + $f5*$m7*.125 + $f5*$m8*.25 + $f6*$m3*.0625 + $f6*$m4*.0625 + $f6*$m5*.0625 + $f6*$m6*.125 + $f6*$m7*.125 + $f6*$m8*.125 + $f7*$m3*.125 + $f7*$m4*.0625 + $f7*$m6*.25 + $f7*$m7*.125 + $f8*$m4*.125 + $f8*$m5*.25 + $f8*$m7*.25 + $f8*$m8*.5 + $f9*$m3*.125 + $f9*$m4*.125 + $f9*$m5*.125 + $f9*$m6*.25 + $f9*$m7*.25 + $f9*$m8*.25 + $m9*$m3*.25 + $m9*$m4*.125 + $m9*$m6*.5 + $m9*$m7*.25,

		$f1*$m1*.5 + $f1*$m2*1 + $f1*$m4*.25 + $f1*$m5*.5 + $f2*$m1*.25 + $f2*$m2*.5 + $f2*$m4*.125 + $f2*$m5*.25 + $f3*$m1*.125 + $f3*$m2*.25 + $f3*$m3*.125 + $f3*$m4*.125 + $f3*$m5*.125 + $f4*$m1*.25 + $f4*$m3*.25 + $f4*$m4*.125 + $f5*$m1*.125 + $f5*$m2*.25 + $f5*$m4*.0625 + $f5*$m5*.125 + $f6*$m1*.125 + $f6*$m2*.125 + $f6*$m3*.0625 + $f6*$m4*.0625 + $f6*$m5*.0625 + $f7*$m1*.125 + $f7*$m3*.125 + $f7*$m4*.0625,
		$f3*$m1*.125 + $f3*$m2*.25 + $f3*$m4*.0625 + $f3*$m5*.125 + $f4*$m1*.25 + $f4*$m2*.5 + $f4*$m4*.125 + $f4*$m5*.25 + $f6*$m1*.0625 + $f6*$m2*.125 + $f6*$m4*.03125 + $f6*$m5*.0625 + $f7*$m1*.125 + $f7*$m2*.25 + $f7*$m4*.0625 + $f7*$m5*.125,
		$f1*$m3*.5 + $f1*$m4*.25 + $f1*$m6*1 + $f1*$m7*.5 + $f2*$m3*.25 + $f2*$m4*.125 + $f2*$m6*.5 + $f2*$m7*.25 + $f3*$m3*.125 + $f3*$m4*.0625 + $f3*$m6*.25 + $f3*$m7*.125 + $f5*$m1*.125 + $f5*$m3*.25 + $f5*$m4*.125 + $f5*$m6*.25 + $f5*$m7*.125 + $f6*$m1*.0625 + $f6*$m3*.125 + $f6*$m4*.0625 + $f6*$m6*.125 + $f6*$m7*.0625 + $f8*$m1*.25 + $f8*$m3*.25 + $f8*$m4*.125 + $f9*$m1*.125 + $f9*$m3*.125 + $f9*$m4*.0625,
		$f1*$m4*.25 + $f1*$m5*.5 + $f1*$m7*.5 + $f1*$m8*1 + $f2*$m4*.125 + $f2*$m5*.25 + $f2*$m7*.25 + $f2*$m8*.5 + $f3*$m3*.125 + $f3*$m4*.125 + $f3*$m5*.125 + $f3*$m6*.25 + $f3*$m7*.25 + $f3*$m8*.25 + $f4*$m3*.25 + $f4*$m4*.125 + $f4*$m6*.5 + $f4*$m7*.25 + $f5*$m1*.125 + $f5*$m2*.25 + $f5*$m4*.125 + $f5*$m5*.25 + $f5*$m7*.125 + $f5*$m8*.25 + $f6*$m1*.125 + $f6*$m2*.125 + $f6*$m3*.125 + $f6*$m4*.125 + $f6*$m5*.125 + $f6*$m6*.125 + $f6*$m7*.125 + $f6*$m8*.125 + $f7*$m1*.125 + $f7*$m3*.25 + $f7*$m4*.125 + $f7*$m6*.25 + $f7*$m7*.125 + $f8*$m1*.25 + $f8*$m2*.5 + $f8*$m4*.125 + $f8*$m5*.25 + $f9*$m1*.25 + $f9*$m2*.25 + $f9*$m3*.125 + $f9*$m4*.125 + $f9*$m5*.125 + $m9*$m1*.25 + $m9*$m3*.25 + $m9*$m4*.125,
		$f3*$m4*.0625 + $f3*$m5*.125 + $f3*$m7*.125 + $f3*$m8*.25 + $f4*$m4*.125 + $f4*$m5*.25 + $f4*$m7*.25 + $f4*$m8*.5 + $f6*$m1*.0625 + $f6*$m2*.125 + $f6*$m4*.0625 + $f6*$m5*.125 + $f6*$m7*.0625 + $f6*$m8*.125 + $f7*$m1*.125 + $f7*$m2*.25 + $f7*$m4*.125 + $f7*$m5*.25 + $f7*$m7*.125 + $f7*$m8*.25 + $f9*$m1*.125 + $f9*$m2*.25 + $f9*$m4*.0625 + $f9*$m5*.125 + $m9*$m1*.25 + $m9*$m2*.5 + $m9*$m4*.125 + $m9*$m5*.25,
		$f5*$m3*.125 + $f5*$m4*.0625 + $f5*$m6*.25 + $f5*$m7*.125 + $f6*$m3*.0625 + $f6*$m4*.03125 + $f6*$m6*.125 + $f6*$m7*.0625 + $f8*$m3*.25 + $f8*$m4*.125 + $f8*$m6*.5 + $f8*$m7*.25 + $f9*$m3*.125 + $f9*$m4*.0625 + $f9*$m6*.25 + $f9*$m7*.125,
		$f5*$m4*.0625 + $f5*$m5*.125 + $f5*$m7*.125 + $f5*$m8*.25 + $f6*$m3*.0625 + $f6*$m4*.0625 + $f6*$m5*.0625 + $f6*$m6*.125 + $f6*$m7*.125 + $f6*$m8*.125 + $f7*$m3*.125 + $f7*$m4*.0625 + $f7*$m6*.25 + $f7*$m7*.125 + $f8*$m4*.125 + $f8*$m5*.25 + $f8*$m7*.25 + $f8*$m8*.5 + $f9*$m3*.125 + $f9*$m4*.125 + $f9*$m5*.125 + $f9*$m6*.25 + $f9*$m7*.25 + $f9*$m8*.25 + $m9*$m3*.25 + $m9*$m4*.125 + $m9*$m6*.5 + $m9*$m7*.25,
		$f6*$m4*.03125 + $f6*$m5*.0625 + $f6*$m7*.0625 + $f6*$m8*.125 + $f7*$m4*.0625 + $f7*$m5*.125 + $f7*$m7*.125 + $f7*$m8*.25 + $f9*$m4*.0625 + $f9*$m5*.125 + $f9*$m7*.125 + $f9*$m8*.25 + $m9*$m4*.125 + $m9*$m5*.25 + $m9*$m7*.25 + $m9*$m8*.5,
		$f6*$m4*.03125 + $f6*$m5*.0625 + $f6*$m7*.0625 + $f6*$m8*.125 + $f7*$m4*.0625 + $f7*$m5*.125 + $f7*$m7*.125 + $f7*$m8*.25 + $f9*$m4*.0625 + $f9*$m5*.125 + $f9*$m7*.125 + $f9*$m8*.25 + $m9*$m4*.125 + $m9*$m5*.25 + $m9*$m7*.25 + $m9*$m8*.5
	);
	
	# Calculate standardized genotype frequencies to sum to 1
	my @after_mating_std;
	for (my $i=0; $i<18; $i++) {
		$after_mating_std[$i] = $after_mating[$i] / sum(@after_mating);
	}

	return @after_mating_std;
}


# Function calculates the allele frequencies 
# given an array of all possible genotypes
sub allele_freqs{
	my @freqs = @_;
	
	my $X_freq = ($freqs[0] + $freqs[1] + $freqs[2] + $freqs[3] + $freqs[9] + $freqs[10]
				+ 0.5*($freqs[4] + $freqs[5] + $freqs[6] + $freqs[11] + $freqs[12] + $freqs[13])) / sum(@freqs);
	my $Y_freq = 1 - $X_freq;

	my $III_freq = ($freqs[0] + $freqs[1] + $freqs[4] + $freqs[7] + $freqs[11] + $freqs[14]
				+ 0.5*($freqs[2] + $freqs[5] + $freqs[8] + $freqs[9] + $freqs[12] + $freqs[15])) / sum(@freqs);
	my $IIIM_freq = 1 - $III_freq;
	
	my $tra_freq = ($freqs[0] + $freqs[9] + $freqs[10] + $freqs[11] + $freqs[12] + $freqs[13] + $freqs[14] + $freqs[15] + $freqs[16]
				+ 0.5*($freqs[1] + $freqs[2] + $freqs[3] + $freqs[4] + $freqs[5] + $freqs[6] + $freqs[7] + $freqs[8] + $freqs[17])) / sum(@freqs);
	my $traD_freq = 1 - $tra_freq;

	return ($X_freq, $Y_freq, $III_freq, $IIIM_freq, $tra_freq, $traD_freq);
}

# Function calculates sex ratio
# given an array of all possible genotypes
sub sex_ratio{
	my @freqs = @_;

	my $female = ($freqs[0] + $freqs[1] + $freqs[2] + $freqs[3] + $freqs[4] + $freqs[5] + $freqs[6] + $freqs[7] + $freqs[8] + $freqs[17])/sum(@freqs);
	my $male = ($freqs[9] + $freqs[10] + $freqs[11] + $freqs[12] + $freqs[13] + $freqs[14] + $freqs[15] + $freqs[16])/sum(@freqs);
	
	return($female, $male);
}


# Function determines the genotype fitness from allele fitness values
# send array with 5 fitness values
sub genotype_fitness{
	my($IIIM_fem, $YM_fem, $traD_fem, $IIIM_male, $YM_male) = @_;
	
	
	# Calculate single locus fitness effects of genotypes:	
	
	# IIIM female
	my $III_fem;  # III /III  (female)
	my $IIIh_fem; # IIIM/III  (female)
	my $IIIm_fem; # IIIM/IIIM (female)
	if($IIIM_fem < 0){ #if IIIM is deleterious to females
		$III_fem  = 1;
		$IIIh_fem = 1 + 0.5 * $IIIM_fem;
		$IIIm_fem = 1 + $IIIM_fem;
	}else{ # if IIIM is beneficial to females
		$III_fem  = 1 - $IIIM_fem;
		$IIIh_fem = 1 - 0.5 * $IIIM_fem;
		$IIIm_fem = 1; 
	}
	
	# YM female
	my $XX_fem; # X/X (female)
	my $XY_fem; # X/Y (female)
	my $YY_fem; # Y/Y (female)
	if($YM_fem < 0){ #if YM is deleterious to females
		$XX_fem = 1;
		$XY_fem = 1 + 0.5 * $YM_fem;
		$YY_fem = 1 + $YM_fem;
	}else{ # if YM is beneficial to females
		$XX_fem = 1 - $YM_fem;
		$XY_fem = 1 - 0.5 * $YM_fem;
		$YY_fem = 1; 	
	}
		
	# traD female
	my $tra_fem; # tra+/tra+ (female)
	my $trad_fem; # traD/tra+ (female)
	if($traD_fem < 0){ # if traD is deleterious to females
		$tra_fem = 1;
		$trad_fem = 1 + $traD_fem;
	}else{ # if traD is beneficial to females
		$tra_fem = 1 - $traD_fem;
		$trad_fem = 1;
	}
	
	# IIIM male
	my $III_male;  # III /III  (male)
	my $IIIh_male; # IIIM/III  (male)
	my $IIIm_male; # IIIM/IIIM (male)
	if($IIIM_male < 0){ #if IIIM is deleterious to males
		$III_male  = 1;
		$IIIh_male = 1 + 0.5 * $IIIM_male;
		$IIIm_male = 1 + $IIIM_male;
	}else{ # if IIIM is beneficial to males
		$III_male  = 1 - $IIIM_male;
		$IIIh_male = 1 - 0.5 * $IIIM_male;
		$IIIm_male = 1; 
	}

	# YM male
	my $XX_male; # X/X (male)
	my $XY_male; # X/Y (male)
	my $YY_male; # Y/Y (male)
	if($YM_male < 0){ #if YM is deleterious to males
		$XX_male = 1;
		$XY_male = 1 + 0.5 * $YM_male;
		$YY_male = 1 + $YM_male;
	}else{ # if YM is beneficial to males
		$XX_male = 1 - $YM_male;
		$XY_male = 1 - 0.5 * $YM_male;
		$YY_male = 1; 	
	}
	

	# Calculate the genotype fitness 
	my @genotype_fit = (
		$III_fem  * $tra_fem  * $XX_fem ,		# 1 female f1 M0 F0 Y0	III /III	tra+/tra+	X/X [0]
		$III_fem  * $trad_fem * $XX_fem ,		# 2 female f2 M0 F1 Y0	III /III	traD/tra+	X/X [1]
		$IIIh_fem * $trad_fem * $XX_fem ,		# 3 female f3 M1 F1 Y0	IIIM/III	traD/tra+	X/X [2]
		$IIIm_fem * $trad_fem * $XX_fem ,		# 4 female f4 M2 F1 Y0	IIIM/IIIM	traD/tra+	X/X [3]
		$III_fem  * $trad_fem * $XY_fem , 		# 5 female f5 M0 F1 Y1	III /III	traD/tra+	X/Y [4]
		$IIIh_fem * $trad_fem * $XY_fem ,		# 6 female f6 M1 F1 Y1	IIIM/III	traD/tra+	X/Y [5]
		$IIIm_fem * $trad_fem * $XY_fem ,		# 7 female f7 M2 F1 Y1	IIIM/IIIM	traD/tra+	X/Y [6]
		$III_fem  * $trad_fem * $YY_fem ,		# 8 female f8 M0 F1 Y2	III /III	traD/tra+	Y/Y [7]
		$IIIh_fem * $trad_fem * $YY_fem ,		# 9 female f9 M1 F1 Y2	IIIM/III	traD/tra+	Y/Y [8]
		$IIIh_male * $XX_male ,					#10 male   m1 M1 F0 Y0	IIIM/III	tra+/tra+	X/X [9]
		$IIIm_male * $XX_male ,					#11 male   m2 M2 F0 Y0	IIIM/IIIM	tra+/tra+	X/X [10]
		$III_male  * $XY_male ,					#12 male   m3 M0 F0 Y1	III /III	tra+/tra+	Y/X [11]
		$IIIh_male * $XY_male ,					#13 male   m4 M1 F0 Y1	IIIM/III	tra+/tra+	Y/X [12]
		$IIIm_male * $XY_male ,					#14 male   m5 M2 F0 Y1	IIIM/IIIM	tra+/tra+	Y/X [13]
		$III_male  * $YY_male ,					#15 male   m6 M0 F0 Y2	III /III	tra+/tra+	Y/Y [14]
		$IIIh_male * $YY_male ,					#16 male   m7 M1 F0 Y2	IIIM/III	tra+/tra+	Y/Y [15]
		$IIIm_male * $YY_male ,					#17 male   m8 M2 F0 Y2	IIIM/IIIM	tra+/tra+	Y/Y [16]
		$IIIm_fem * $trad_fem * $YY_fem			#18 female m9 M2 F1 Y2	IIIM/IIIM	traD/tra+	Y/Y [17] <- should be f10		
	);
	
	my @genotype_norm = @genotype_fit;
	# Normalize male fitness to ensure that max = 1
	for(my $i=9; $i < 17; $i++){
		$genotype_norm[$i] = $genotype_fit[$i] / max(@genotype_fit[9..16]);
	}
	# Normalize female fitness to ensure that max = 1
	for(my $i=0; $i < 9; $i++){
		$genotype_norm[$i] = $genotype_fit[$i] / max(@genotype_fit[0..8], $genotype_fit[17]);
	}
	$genotype_norm[17] = $genotype_fit[17] / max(@genotype_fit[0..8], $genotype_fit[17]);

	# return normalize genotype fitness values
	return(@genotype_norm);
}
```
