## Supplementary material for "The maintenance of polygenic sex determination depends on the dominance of fitness effects which are predictive of the role of sexual antagonism": Code: seln_allele_finite_2020_09.pl

```
#!/usr/bin/perl

# Use multinomial sampling each generation with a given population size
#	with best-fitting selection pressures
# Start at observed genotype frequencies to see if it will hold. 

# perl seln_allele_finite_2020_09.pl generations popsize reps control ctrlfile output

# Do 100 reps for 1000 generations
# perl seln_allele_finite_2020_09.pl 1000 10000 100 additive_CA_top1k_first.tsv CA_femHW.txt additive_CA_top1k_first


# "ctrfile" is observed genotype frequencies in population
# use this to determine initial genotype frequencies in simulations
# these are the same starting values as in the drift simulation


# "control" file contains best-fitting selection values 
# use this to get the fitness values for each genotype
# 0						1					2					3					4					5					6					7					8					9					10					11					12					13					14					15					16					17					18					19					20					21					22					23					24					25						26						27					28						29						30					31						32						33						34					35						36						37					38						39						40						41					42					43					44					45					46						47					48					49					50					51						52						53					54						55						56					57					58						59						60					61						62						63					64						65						66						67					68					69					70					71						72					73
# IIIM_fem				YM_fem				traD_fem			IIIM_male			YM_male				f1_fit				f2_fit				f3_fit				f4_fit				f5_fit				f6_fit				f7_fit				f8_fit				f9_fit				m1_fit				m2_fit				m3_fit				m4_fit				m5_fit				m6_fit				m7_fit				m8_fit				m9_fit				f1					f2					f3						f4						f5					f6						f7						f8					f9						m1						m2						m3					m4						m5						m6					m7						m8						m9						female				male				X					Y					III					IIIM					tra					traD				f1_exp				f2_exp				f3_exp					f4_exp					f5_exp				f6_exp					f7_exp					f8_exp				f9_exp				m1_exp					m2_exp					m3_exp				m4_exp					m5_exp					m6_exp				m7_exp					m8_exp					m9_exp					al0_exp				al1_exp				al2_exp				al0_obs_exp			al1_obs_exp				al2_obs_exp			MSE_allele
# -0.359135066869892	-0.984522935321764	0.746590366801332	0.413181123379802	0.737774379416752	0.341567352483462	0.768672892731473	0.563220768543688	0.563220768543688	0.205452124187785	0					0					0.205452124187785	0					0.577937864036513	0.577937864036513	0.763629488446949	1					1					0.763629488446949	1					1					0					0.358105786873304	0.0865109283609139	1.49433343748673e-06	5.5741469689477e-12		0.111210559493149	9.88407492238908e-07	1.4973025169851e-12		0.0184074520134298	5.36188909278174e-08	4.70567688842453e-06	5.5741469689477e-12		0.407348674550655	1.85102537543388e-06	1.4973025169851e-12		0.0184074520134298	5.36188909278174e-08	0						0						0.574237263107689	0.425762736892311	0.703903951995525	0.296096048004475	0.999995426645369	4.57335463055486e-06	0.891934261882808	0.108065738117192	0.0335158777694521	0.00248579501471527	4.48843793901366e-05	0.0773722359475807		0.00400998749544372	4.47897267194173e-05	0.0446486611334997		0.0013273283746633	0.0155220245941004	4.48843793901366e-05	0.430934924183599		0.00745687576968254	4.47897267194173e-05	0.0446486611334997		0.0013273283746633	8.69658884887749e-06	8.69658884887749e-06	0.648098826381104		0.648098826381104	0.983738588798449	0.918270871784843	0.055805125614421	0.01625683784692		-0.026336609902035	0.00407211384275432
# 0.304099801391588		-0.86802171207119	0.698972937843157	-0.345951127077381	0.990069156119544	0.256445520139773	0.774577893701083	1					1					0.131133597608582	0.356555703907499	0.356555703907499	0.131133597608582	0.356555703907499	0					0					0.990360746433085	0.733915226293312	0.733915226293312	0.990360746433085	0.733915226293312	0.733915226293312	0.356555703907499	0.32272510573245	0.128656479610996	2.32876330093792e-16	7.86844550295347e-32	0.153702224936924	3.16863362130632e-16	1.21444964640425e-31	0.0153776258487765	5.89850819418204e-17	3.2011285732335e-16		7.86844550295347e-32	0.364160938022076	4.2813728965796e-16		1.21444964640425e-31	0.0153776258487765	5.89850819418204e-17	2.68878965315965e-32	2.68878965315965e-32	0.620461436129147	0.379538563870853	0.710313166822946	0.289686833177054	0.999999999999999	8.88178419700125e-16	0.851131834801651	0.148868165198349	0.0335158777694521	0.00248579501471527	4.48843793901366e-05	0.0773722359475807		0.00400998749544372	4.47897267194173e-05	0.0446486611334997		0.0013273283746633	0.0155220245941004	4.48843793901366e-05	0.430934924183599		0.00745687576968254	4.47897267194173e-05	0.0446486611334997		0.0013273283746633	8.69658884887749e-06	8.69658884887749e-06	0.648098826381104		0.648098826381104	0.983738588798449	0.918270871784843	0.0622143404418419	0.01626141120155		-0.067139036983192	0.00864270793790973
# 0.940895181603772		-0.755269873513812	0.561254870475579	-0.623820715595391	-0.406166143493784	0.40676586425726	0.628418519767334	1					1					0.330144808664852	0.701726288897518	0.701726288897518	0.330144808664852	0.701726288897518	0.160404495389764	0.160404495389764	0.246361368867496	0					0					0.246361368867496	0					0					0.701726288897518	0.414228256212976	0.0340782966291516	2.25543983308936e-11	1.43131946596933e-21	0.0532713068002032	3.55343585078971e-11	9.01007506419852e-22	0.0178424382642311	1.36499289275138e-11	4.95825171364549e-11	1.43131946596933e-21	0.462737263658701	3.55343585078971e-11	9.01007506419852e-22	0.0178424382642311	1.36499289275138e-11	0						0						0.519420297978301	0.480579702021699	0.706310838179251	0.293689161820749	0.999999999914747	8.52528048156387e-11	0.947403979117338	0.0525960208826622	0.0335158777694521	0.00248579501471527	4.48843793901366e-05	0.0773722359475807		0.00400998749544372	4.47897267194173e-05	0.0446486611334997		0.0013273283746633	0.0155220245941004	4.48843793901366e-05	0.430934924183599		0.00745687576968254	4.47897267194173e-05	0.0446486611334997		0.0013273283746633	8.69658884887749e-06	8.69658884887749e-06	0.648098826381104		0.648098826381104	0.983738588798449	0.918270871784843	0.0582120117981469	0.016261411116298		0.029133107332495	0.00450180975192753
# -0.710459630651897	-0.680419925058118	0.953600403878854	0.608533052134006	0.648703706515867	0.420979399794359	0.640569812821745	0.47696866175307	0.313367510684395	0.483886057479548	0.320284906410873	0.156683755342197	0.32720230213735	0.163601151068675	0.561109399441359	0.701239399088358	0.57035970025018	0.71048969989718	0.850619699544179	0.719740000706001	0.859870000353001	1					0					0.236153416683278	0.0671102401567787	0.0124903285775913		0.000491146043109489	0.131325505466398	0.0161714047593005		0.000416735418435208	0.0574890272359578	0.0045593191403587		0.0341722438542178		0.000491146043109489	0.350088030774063	0.0264013906245354		0.000416735418435208	0.0574890272359578	0.0045593191403587		8.74917140577775e-05	8.74917140577775e-05	0.526294615195265	0.473705384804735	0.613318422588668	0.386681577411332	0.948832250600614	0.0511677493993861		0.854929400744006	0.145070599255994	0.336553558819184	0.0335158777694521	0.00248579501471527		4.48843793901366e-05	0.0773722359475807	0.00400998749544372		4.47897267194173e-05	0.0446486611334997	0.0013273283746633	0.0155220245941004		4.48843793901366e-05	0.430934924183599	0.00745687576968254		4.47897267194173e-05	0.0446486611334997	0.0013273283746633		8.69658884887749e-06	8.69658884887749e-06	0.648098826381104	0.983738588798449	0.918270871784843	-0.0347804037924361	-0.034906338197835		-0.063341471040837	0.00644027088796372
# 0.494723892329042		-0.26717679654715	0.512406243269579	-0.372213200031836	0.0511139870363704	0.331404036106637	0.559517248012143	0.779758624006072	1					0.440575377167431	0.66081675316136	0.881058129155288	0.321633506322719	0.541874882316647	0.165702018053318	0					0.354158980858605	0.188456962805287	0.0227549447519682	0.376913925610573	0.211211907557255	0.0455098895039364	0.762116258310576	0.138809380034938	0.0646929840708424	0.0041674190923489		5.59055430437316e-05	0.184271728225137	0.0115201006737476		0.000136883469105194	0.12180661822812	0.00766842890124205		0.00556833587666215		5.59055430437316e-05	0.318081670593226	0.0133862026269885		0.000136883469105194	0.12180661822812	0.00766842890124205		8.32532615436825e-05	8.32532615436825e-05	0.533212701500068	0.466787298499932	0.477116664689533	0.522883335310467	0.974458457416499	0.0255415425835009		0.802798339267435	0.197201660732565	0.336553558819184	0.0335158777694521	0.00248579501471527		4.48843793901366e-05	0.0773722359475807	0.00400998749544372		4.47897267194173e-05	0.0446486611334997	0.0013273283746633	0.0155220245941004		4.48843793901366e-05	0.430934924183599	0.00745687576968254		4.47897267194173e-05	0.0446486611334997	0.0013273283746633		8.69658884887749e-06	8.69658884887749e-06	0.648098826381104	0.983738588798449	0.918270871784843	-0.170982161691571	-0.00928013138195005	-0.115472532517408	0.0426549262211726

# IIIM_fem				YM_fem				traD_fem			IIIM_male			YM_male				f1_fit				f2_fit	f3_fit				f4_fit				f5_fit				f6_fit				f7_fit				f8_fit				f9_fit				m1_fit				m2_fit				m3_fit				m4_fit				m5_fit				m6_fit				m7_fit				m8_fit				m9_fit				f1					f2					f3						f4						f5					f6						f7						f8					f9						m1						m2						m3					m4						m5						m6					m7						m8						m9						X					Y					III					IIIM				tra					traD				female				male				f1_exp				f2_exp				f3_exp				f4_exp					f5_exp				f6_exp				f7_exp					f8_exp				f9_exp				m1_exp				m2_exp					m3_exp				m4_exp				m5_exp					m6_exp				m7_exp				m8_exp					m9_exp					al0_exp				al1_exp				al2_exp				al0_obs_exp			al1_obs_exp			al2_obs_exp			MSE_allele
# -0.785199363595781	-0.800255971505194	0.559603191244542	-0.297404538038832	-0.309263335209053	0.440396808755458	1		0.60740031820211	0.214800636404219	0.599872014247403	0.364362452334413	0.128852890421423	0.199744028494806	0.121324586466717	1					0.825322841107388	0.993034870916101	0.845368332395474	0.697701793874847	0.811392582939589	0.690736664790947	0.570080746642306	0.0429051444386269	0.259406350668249	0.105778792987872	0.00282252979107911		1.42860607842837e-05	0.168327411418258	0.0022647848486802		6.00905001021688e-06	0.0490989215769643	0.000352679409031465	0.0058692756142779		1.42860607842837e-05	0.353688887984041	0.00289691106635522		6.00905001021688e-06	0.0490989215769643	0.000352679409031465	6.31713803090387e-07	6.31713803090387e-07	0.637500527891725	0.362499472108275	0.992678716281577	0.007321283718423	0.835666976571758	0.164333023428242	0.588072397524732	0.411927602475267	0.336553558819184	0.0335158777694521	0.00248579501471527	4.48843793901366e-05	0.0773722359475807	0.00400998749544372	4.47897267194173e-05	0.0446486611334997	0.0013273283746633	0.0155220245941004	4.48843793901366e-05	0.430934924183599	0.00745687576968254	4.47897267194173e-05	0.0446486611334997	0.0013273283746633	8.69658884887749e-06	8.69658884887749e-06	0.648098826381104	0.983738588798449	0.918270871784843	-0.0105982984893791	0.008940127483128	-0.082603895213085	0.00701565331465888
# -0.244502799484291	-0.512461409233147	0.819123065390373	-0.873279793760172	-0.763652445706752	0.180876934609627	1		0.877748600257855	0.755497200515709	0.743769295383427	0.652842457937573	0.56191562049172	0.487538590766853	0.427936315617292	0.911329668042989	0.204991235352533	1					0.563360103119914	0.126720206239828	0.382331899266553	0.215390538196838	0.0484491771271227	0.368334040467731	0.154761929255325	0.122261627351012	0.000579000156880889	6.41650830831881e-07	0.236578004061478	0.000716618063468083	4.67364054293451e-07	0.102575768348287	0.00018972561512925		0.000675092423931418	6.41650830831881e-07	0.278148422745527	0.000745934418128516	4.67364054293451e-07	0.102575768348287	0.00018972561512925		8.27838224205601e-08	8.27838224205601e-08	0.536373889497167	0.463626110502833	0.998449568256251	0.00155043174374891	0.768549032302518	0.231450967697482	0.617663864650289	0.382336135349711	0.336553558819184	0.0335158777694521	0.00248579501471527	4.48843793901366e-05	0.0773722359475807	0.00400998749544372	4.47897267194173e-05	0.0446486611334997	0.0013273283746633	0.0155220245941004	4.48843793901366e-05	0.430934924183599	0.00745687576968254	4.47897267194173e-05	0.0446486611334997	0.0013273283746633	8.69658884887749e-06	8.69658884887749e-06	0.648098826381104	0.983738588798449	0.918270871784843	-0.111724936883937	0.014710979457802	-0.149721839482325	0.0351155036562987
# -0.748546966643302	-0.806046098244338	0.476507789223996	0.570274248473957	0.837149777523117	0.523492210776004	1		0.625726516678349	0.251453033356698	0.596976950877831	0.373544308010047	0.150111665142263	0.193953901755662	0.121362099341745	0.116415578358472	0.162850222476883	0.249853342883052	0.415639227060747	0.581425111238441	0.429725751526043	0.714862875763021	1					0.0487702969278281	0.173805483551242	0.0693816284877238	0.0279665354892006		0.00230775657131129		0.129832544036102	0.0474990454839616		0.00372057563318891		0.0431792137075186	0.0150893713212651		0.0575098904589112		0.00230775657131129		0.279555094406178	0.0836000151467979		0.00372057563318891		0.0431792137075186	0.0150893713212651		0.00112796423665755		0.00112796423665755		0.607242976299409	0.392757023700591	0.862310292506984	0.137689707493016	0.829947682516535	0.170052317483465	0.513910118518172	0.486089881481828	0.336553558819184	0.0335158777694521	0.00248579501471527	4.48843793901366e-05	0.0773722359475807	0.00400998749544372	4.47897267194173e-05	0.0446486611334997	0.0013273283746633	0.0155220245941004	4.48843793901366e-05	0.430934924183599	0.00745687576968254	4.47897267194173e-05	0.0446486611334997	0.0013273283746633	8.69658884887749e-06	8.69658884887749e-06	0.648098826381104	0.983738588798449	0.918270871784843	-0.040855850081695	-0.121428296291465	-0.0883231892683081	0.0242150173886711
# -0.892925776959719	-0.84463276378521	0.635916497117329	-0.269900813671559	-0.326980455216644	0.364083502882671	1		0.553537111520141	0.107074223040281	0.577683618107395	0.319769321339671	0.0618550245719479	0.15536723621479	0.0860015311592023	1					0.843996913122459	0.96700787908801	0.836509772391678	0.706011665695347	0.778012671298478	0.673019544783356	0.568026418268235	0.0166358261036145	0.238939468759646	0.118786078935442	0.0366160400043256		0.00198044911691874		0.149591474510354	0.0225544730449			0.000664356319303722	0.0333297495417739	0.00272652279365436		0.0636010068850701		0.00198044911691874		0.264863221928903	0.0275349133070282		0.000664356319303722	0.0333297495417739	0.00272652279365436		5.55835405148278e-05	5.55835405148278e-05	0.694839890533217	0.305160109466783	0.916719482632209	0.0832805173677907	0.816847636096407	0.183152363903593	0.605244196566833	0.394755803433167	0.336553558819184	0.0335158777694521	0.00248579501471527	4.48843793901366e-05	0.0773722359475807	0.00400998749544372	4.47897267194173e-05	0.0446486611334997	0.0013273283746633	0.0155220245941004	4.48843793901366e-05	0.430934924183599	0.00745687576968254	4.47897267194173e-05	0.0446486611334997	0.0013273283746633	8.69658884887749e-06	8.69658884887749e-06	0.648098826381104	0.983738588798449	0.918270871784843	0.046741064152113	-0.06701910616624	-0.101423235688436	0.0169629604069057
# -0.123496451180986	-0.844106532982111	0.936407176123285	-0.860418393339758	-0.506621173612214	0.0635928238767149	1		0.938251774409507	0.876503548819014	0.577946733508945	0.542259548228946	0.506572362948947	0.155893467017889	0.146267322048385	0.763089436199309	0.186933957002543	1					0.569790803330121	0.139581606660242	0.660755084603925	0.376492170460932	0.0922292563179397	0.13664117707888	0.15558714992699	0.142563211913015	0.0172709055420959		0.000507728226447739	0.235151164703788	0.0182450551592673		0.000306586151001305	0.0671060471544096	0.00363174006504819		0.0179121674649546		0.000507728226447739	0.251514429731093	0.0185614461983784		0.000306586151001305	0.0671060471544096	0.00363174006504819		4.51330833024305e-05	4.51330833024305e-05	0.596391525347215	0.403608474652785	0.958654577831101	0.0413454221688992	0.757586214000812	0.242413785999188	0.640414721925365	0.359585278074635	0.336553558819184	0.0335158777694521	0.00248579501471527	4.48843793901366e-05	0.0773722359475807	0.00400998749544372	4.47897267194173e-05	0.0446486611334997	0.0013273283746633	0.0155220245941004	4.48843793901366e-05	0.430934924183599	0.00745687576968254	4.47897267194173e-05	0.0446486611334997	0.0013273283746633	8.69658884887749e-06	8.69658884887749e-06	0.648098826381104	0.983738588798449	0.918270871784843	-0.0517073010338891	-0.025084010967348	-0.160684657784031	0.0291224118335904


# Columns in 'control' file: 

#	Allele fitness effects: 
# 0 IIIM female fitness effect
# 1   YM female fitness effect
# 2	traD female fitness effect
# 3 IIIM   male fitness effect
# 4   YM   male fitness effect

#	Genotype fitness values: 
# 5 female f1 M0 F0 Y0	III /III	tra+/tra+	X/X
# 6 female f2 M0 F1 Y0	III /III	traD/tra+	X/X
# 7 female f3 M1 F1 Y0	IIIM/III	traD/tra+	X/X
# 8 female f4 M2 F1 Y0	IIIM/IIIM	traD/tra+	X/X
# 9 female f5 M0 F1 Y1	III /III	traD/tra+	X/Y
#10 female f6 M1 F1 Y1	IIIM/III	traD/tra+	X/Y
#11 female f7 M2 F1 Y1	IIIM/IIIM	traD/tra+	X/Y
#12 female f8 M0 F1 Y2	III /III	traD/tra+	Y/Y
#13 female f9 M1 F1 Y2	IIIM/III	traD/tra+	Y/Y
#14 male   m1 M1 F0 Y0	IIIM/III	tra+/tra+	X/X
#15 male   m2 M2 F0 Y0	IIIM/IIIM	tra+/tra+	X/X
#16 male   m3 M0 F0 Y1	III /III	tra+/tra+	Y/X
#17 male   m4 M1 F0 Y1	IIIM/III	tra+/tra+	Y/X
#18 male   m5 M2 F0 Y1	IIIM/IIIM	tra+/tra+	Y/X
#19 male   m6 M0 F0 Y2	III /III	tra+/tra+	Y/Y
#20 male   m7 M1 F0 Y2	IIIM/III	tra+/tra+	Y/Y
#21 male   m8 M2 F0 Y2	IIIM/IIIM	tra+/tra+	Y/Y
#22 female m9 M2 F1 Y2	IIIM/IIIM	traD/tra+	Y/Y

#	Frequencies:
#23 female f1 M0 F0 Y0	III /III	tra+/tra+	X/X
#24 female f2 M0 F1 Y0	III /III	traD/tra+	X/X
#25 female f3 M1 F1 Y0	IIIM/III	traD/tra+	X/X
#26 female f4 M2 F1 Y0	IIIM/IIIM	traD/tra+	X/X
#27 female f5 M0 F1 Y1	III /III	traD/tra+	X/Y
#28 female f6 M1 F1 Y1	IIIM/III	traD/tra+	X/Y
#29 female f7 M2 F1 Y1	IIIM/IIIM	traD/tra+	X/Y
#30 female f8 M0 F1 Y2	III /III	traD/tra+	Y/Y
#31 female f9 M1 F1 Y2	IIIM/III	traD/tra+	Y/Y
#32 male   m1 M1 F0 Y0	IIIM/III	tra+/tra+	X/X
#33 male   m2 M2 F0 Y0	IIIM/IIIM	tra+/tra+	X/X
#34 male   m3 M0 F0 Y1	III /III	tra+/tra+	Y/X
#35 male   m4 M1 F0 Y1	IIIM/III	tra+/tra+	Y/X
#36 male   m5 M2 F0 Y1	IIIM/IIIM	tra+/tra+	Y/X
#37 male   m6 M0 F0 Y2	III /III	tra+/tra+	Y/Y
#38 male   m7 M1 F0 Y2	IIIM/III	tra+/tra+	Y/Y
#39 male   m8 M2 F0 Y2	IIIM/IIIM	tra+/tra+	Y/Y
#40 female m9 M2 F1 Y2	IIIM/IIIM	traD/tra+	Y/Y
#41 female	
#42   male	
#43      X	
#44      Y	
#45    III	
#46    IIIM	
#47    tra	
#48    traD	


#	Expected genotype frequencies -- also starting values
#49 female f1 M0 F0 Y0	III /III	tra+/tra+	X/X
#50 female f2 M0 F1 Y0	III /III	traD/tra+	X/X
#51 female f3 M1 F1 Y0	IIIM/III	traD/tra+	X/X
#52 female f4 M2 F1 Y0	IIIM/IIIM	traD/tra+	X/X
#53 female f5 M0 F1 Y1	III /III	traD/tra+	X/Y
#54 female f6 M1 F1 Y1	IIIM/III	traD/tra+	X/Y
#55 female f7 M2 F1 Y1	IIIM/IIIM	traD/tra+	X/Y
#56 female f8 M0 F1 Y2	III /III	traD/tra+	Y/Y
#57 female f9 M1 F1 Y2	IIIM/III	traD/tra+	Y/Y
#58 male   m1 M1 F0 Y0	IIIM/III	tra+/tra+	X/X
#59 male   m2 M2 F0 Y0	IIIM/IIIM	tra+/tra+	X/X
#60 male   m3 M0 F0 Y1	III /III	tra+/tra+	Y/X
#61 male   m4 M1 F0 Y1	IIIM/III	tra+/tra+	Y/X
#62 male   m5 M2 F0 Y1	IIIM/IIIM	tra+/tra+	Y/X
#63 male   m6 M0 F0 Y2	III /III	tra+/tra+	Y/Y
#64 male   m7 M1 F0 Y2	IIIM/III	tra+/tra+	Y/Y
#65 male   m8 M2 F0 Y2	IIIM/IIIM	tra+/tra+	Y/Y
#66 female m9 M2 F1 Y2	IIIM/IIIM	traD/tra+	Y/Y


# 	Expected allele frequencies
#67	  X:	 al0_exp	
#68	III: al1_exp	
#69	tra: al2_exp	

#	Fit of allele frequencies
#70	  X:	 al0_obs_exp	
#71 III: al1_obs_exp	
#72 tra: al2_obs_exp	
#73 MSE: MSE_allele


use strict;
use warnings;
use List::Util qw(sum min max);
use POSIX; # for rounding
use Math::Random qw(random_binomial); # for random number generation: random_binomial($n, $nt, $p)
	# $n is number of outcomes
	# $nt is number of trials
	# $p is probability of success


my $popsize=10000; # population size (can be reset by user)
my $generations=1000; # number of generations to run each simulation (can be reset by user)
my $reps=100; # how many replicates per parameter/selection vector (can be reset by user)
my $controlfile; # fitness values
my $ctrlfile; # initial genotype frequencies
my $outfile;
my @initial_freq;

($generations, $popsize, $reps, $controlfile, $ctrlfile, $outfile) = @ARGV;


# Make sure generations are >0
if($generations<1){
	die "\nGenerations must be >0\n\n";
}

# Make sure population size is >0
if($popsize<1){
	die "\nPopulation size must be >0\n\n";
}

# Make sure reps are >0
if($reps<1){
	die "\nReplicates must be >0\n\n";
}


# obtain initial genotype frequencies for simulation
# only first line is necessary
open INPUT, "$ctrlfile" or die "\nCould not open control file, $ctrlfile\n";
my $linecounter=0;
while(<INPUT>){
	if($linecounter==0){
		chomp;
		@initial_freq = split /\s+/, $_; # delimit each genotype frequency by white space
	}else{
		last;
	}
	$linecounter++;
}
close INPUT;
#print "\nInitial genotypes: scalar(@initial_freq)\n";
if(scalar(@initial_freq) != 18){
	die "\nInitial genotypes should be 18\n\n";
}


# 1 female f1 M0 F0 Y0	III /III	tra+/tra+	X/X
# 2 female f2 M0 F1 Y0	III /III	traD/tra+	X/X
# 3 female f3 M1 F1 Y0	IIIM/III	traD/tra+	X/X
# 4 female f4 M2 F1 Y0	IIIM/IIIM	traD/tra+	X/X
# 5 female f5 M0 F1 Y1	III /III	traD/tra+	X/Y
# 6 female f6 M1 F1 Y1	IIIM/III	traD/tra+	X/Y
# 7 female f7 M2 F1 Y1	IIIM/IIIM	traD/tra+	X/Y
# 8 female f8 M0 F1 Y2	III /III	traD/tra+	Y/Y
# 9 female f9 M1 F1 Y2	IIIM/III	traD/tra+	Y/Y
#10 male   m1 M1 F0 Y0	IIIM/III	tra+/tra+	X/X
#11 male   m2 M2 F0 Y0	IIIM/IIIM	tra+/tra+	X/X
#12 male   m3 M0 F0 Y1	III /III	tra+/tra+	Y/X
#13 male   m4 M1 F0 Y1	IIIM/III	tra+/tra+	Y/X
#14 male   m5 M2 F0 Y1	IIIM/IIIM	tra+/tra+	Y/X
#15 male   m6 M0 F0 Y2	III /III	tra+/tra+	Y/Y
#16 male   m7 M1 F0 Y2	IIIM/III	tra+/tra+	Y/Y
#17 male   m8 M2 F0 Y2	IIIM/IIIM	tra+/tra+	Y/Y
#18 female m9 M2 F1 Y2	IIIM/IIIM	traD/tra+	Y/Y <- should be f10


# Open input file and prep for reading in for each simulation
open INPUT, "$controlfile" or die "\nCould not open input file, $controlfile\n\n";
my $inputline = <INPUT>; # first line is a header
my $counter=0; 


# Run simulations
while(my $inline=<INPUT>){
	$counter++;
	
	# Get information for this parameterization
	chomp $inline;
	my @inarray = split /\t/, $inline; 
	my @allele_fitness = @inarray[0..4];
	my @fitness_array = @inarray[5..22];

	# create output file
	my $outputfilename;
	if($counter < 10){
		$outputfilename = $outfile."_top000".$counter.".out";
	}elsif($counter<100){
		$outputfilename = $outfile."_top00".$counter.".out";
	}elsif($counter<1000){
		$outputfilename = $outfile."_top0".$counter.".out";
	}else{
		$outputfilename = $outfile."_top".$counter.".out";	
	}
	open OUTPUT, ">$outputfilename" or die "\nCould not open output file, $outputfilename\n\n";
	
	# Header
	print OUTPUT "run\t";
	print OUTPUT "IIIM_fem\tYM_fem\ttraD_fem\tIIIM_male\tYM_male\t";
	print OUTPUT "f1_fit	f2_fit	f3_fit	f4_fit	f5_fit	f6_fit	f7_fit	f8_fit	f9_fit	m1_fit	m2_fit	m3_fit	m4_fit	m5_fit	m6_fit	m7_fit	m8_fit	m9_fit\t";
	print OUTPUT "f1\tf2\tf3\tf4\tf5\tf6\tf7\tf8\tf9\t";
	print OUTPUT "m1\tm2\tm3\tm4\tm5\tm6\tm7\tm8\tm9\t";
	print OUTPUT "female\tmale\t";
	print OUTPUT "X\tY\tIII\tIIIM\ttra\ttraD\n";

	
	# Run replicate simulations for this parameterization
	for (my $i=1; $i <= $reps; $i++){
				
		my @simulation_results;
		$simulation_results[0] = [ @initial_freq ];
		for(my $a=1; $a <= $generations; $a++){
			$simulation_results[$a] = [ recursion_seln(\@{$simulation_results[$a-1]}, \@fitness_array, $popsize) ]; # need to pass the function references to the arrays, not the arrays themselves
		}

		# Print initial conditions to file
		print OUTPUT "$i\t";
		for(my $b=0; $b<5; $b++){
			print OUTPUT "$allele_fitness[$b]\t";
		}
		for(my $b=0; $b<18; $b++){
			print OUTPUT "$fitness_array[$b]\t";
		}

		# Print final results of simulation to file

		# Final genotype frequencies
		for(my $b=0; $b<18; $b++){
			print OUTPUT "$simulation_results[$generations][$b]\t";
		}
		
		# Final sex ratio
		my @sex_ratios = sex_ratio(@{$simulation_results[$generations]});
		print OUTPUT "$sex_ratios[0]\t$sex_ratios[1]";
		
		# Final allele frequencies
		my @allele_freq = allele_freqs(@{$simulation_results[$generations]});
		for(my $b=0; $b<6; $b++){
			print OUTPUT "\t$allele_freq[$b]";
		}
		print OUTPUT "\n";
	
	}
	close OUTPUT;

}

close INPUT;

#print "\n$counter parameterizations read from $controlfile and printed to $counter outfiles named $outfile\n\n";


# Function performs 1 generation of selection, followed by genetic drift (multinomial sampling) and random mating
# Send function 2 arrays and one scalar:
#	1. counts of the 18 genotypes
#	2. fitness values for the 18 genotypes
#	3. population size
sub recursion_seln {
	my($raw_counts_ref, $fitness_ref, $popnsize) = @_; # pass the function references to the arrays
	
	# Dereference into arrays
	my @raw_counts = @{$raw_counts_ref}; 
	my @fitness = @{$fitness_ref};

	# impose selection pressures
	my @seln_counts;
	for(my $i=0; $i<18; $i++) {
		$seln_counts[$i] = $raw_counts[$i] * $fitness[$i];
	}
	
	# normalize back to sum to 1
	my @norm_counts;
	for (my $i=0; $i<18; $i++) {
		$norm_counts[$i] = $seln_counts[$i] / sum(@seln_counts);
	}
	
	# multinomial sampling for genetic drift
	# random_binomial($n, $nt, $p)
		# $n is number of outcomes
		# $nt is number of trials
		# $p is probability of success
	my @drift_counts;
	for(my $i=0; $i<18; $i++){
		$drift_counts[$i] = random_binomial(1, $popnsize, $norm_counts[$i]) / $popnsize;
	}
	
	# normalize back to sum to 1
	my @counts;
	for (my $i=0; $i<18; $i++) {
		$counts[$i] = $drift_counts[$i] / sum(@drift_counts);
	}

	
	# Calculate genotype frequencies after random mating
	my $f1 = $counts[0];
	my $f2 = $counts[1];
	my $f3 = $counts[2];
	my $f4 = $counts[3];
	my $f5 = $counts[4];
	my $f6 = $counts[5];
	my $f7 = $counts[6];
	my $f8 = $counts[7];
	my $f9 = $counts[8];

	my $m1 = $counts[9];
	my $m2 = $counts[10];
	my $m3 = $counts[11];
	my $m4 = $counts[12];
	my $m5 = $counts[13];
	my $m6 = $counts[14];
	my $m7 = $counts[15];
	my $m8 = $counts[16];
	my $m9 = $counts[17];
	
	my @after_mating = (
		$f1*$m1*.5 + $f1*$m3*.5 + $f1*$m4*.25 + $f2*$m1*.25 + $f2*$m3*.25 + $f2*$m4*.125 + $f3*$m1*.25 + $f3*$m3*.125 + $f3*$m4*.0625  + $f5*$m1*.125 + $f5*$m3*.125 + $f5*$m4*.0625 + $f6*$m1*.0625 + $f6*$m3*.0625 + $f6*$m4*.03125,
		$f2*$m1*.25 + $f2*$m3*.25 + $f2*$m4*.125 + $f3*$m1*.125 + $f3*$m3*.125 + $f3*$m4*.0625 + $f5*$m1*.125 + $f5*$m3*.125 + $f5*$m4*.0625 + $f6*$m1*.0625 + $f6*$m3*.0625 + $f6*$m4*.03125,
		$f2*$m1*.25 + $f2*$m2*.5 + $f2*$m4*.125 + $f2*$m5*.25 + $f3*$m1*.25 + $f3*$m2*.25 + $f3*$m3*.125 + $f3*$m4*.125 + $f3*$m5*.125 + $f4*$m1*.25 + $f4*$m3*.25 + $f4*$m4*.125 + $f5*$m1*.125 + $f5*$m2*.25 + $f5*$m4*.0625 + $f5*$m5*.125 + $f6*$m1*.125 + $f6*$m2*.125 + $f6*$m3*.0625 + $f6*$m4*.0625 + $f6*$m5*.0625 + $f7*$m1*.125 + $f7*$m3*.125 + $f7*$m4*.0625,
		$f3*$m1*.125 + $f3*$m2*.25 + $f3*$m4*.0625 + $f3*$m5*.125 + $f4*$m1*.25 + $f4*$m2*.5 + $f4*$m4*.125 + $f4*$m5*.25 + $f6*$m1*.0625 + $f6*$m2*.125 + $f6*$m4*.03125 + $f6*$m5*.0625 + $f7*$m1*.125 + $f7*$m2*.25 + $f7*$m4*.0625 + $f7*$m5*.125,
		$f2*$m3*.25 + $f2*$m4*.125 + $f2*$m6*.5 + $f2*$m7*.25 + $f3*$m3*.125 + $f3*$m4*.0625 + $f3*$m6*.25 + $f3*$m7*.125 + $f5*$m1*.125 + $f5*$m3*.25 + $f5*$m4*.125 + $f5*$m6*.25 + $f5*$m7*.125 + $f6*$m1*.0625 + $f6*$m3*.125 + $f6*$m4*.0625 + $f6*$m6*.125 + $f6*$m7*.0625 + $f8*$m1*.25 + $f8*$m3*.25 + $f8*$m4*.125 + $f9*$m1*.125 + $f9*$m3*.125 + $f9*$m4*.0625,
		$f2*$m4*.125 + $f2*$m5*.25 + $f2*$m7*.25 + $f2*$m8*.5 + $f3*$m3*.125 + $f3*$m4*.125 + $f3*$m5*.125 + $f3*$m6*.25 + $f3*$m7*.25 + $f3*$m8*.25 + $f4*$m3*.25 + $f4*$m4*.125 + $f4*$m6*.5 + $f4*$m7*.25 + $f5*$m1*.125 + $f5*$m2*.25 + $f5*$m4*.125 + $f5*$m5*.25 + $f5*$m7*.125 + $f5*$m8*.25 + $f6*$m1*.125 + $f6*$m2*.125 + $f6*$m3*.125 + $f6*$m4*.125 + $f6*$m5*.125 + $f6*$m6*.125 + $f6*$m7*.125 + $f6*$m8*.125 + $f7*$m1*.125 + $f7*$m3*.25 + $f7*$m4*.125 + $f7*$m6*.25 + $f7*$m7*.125 + $f8*$m1*.25 + $f8*$m2*.5 + $f8*$m4*.125 + $f8*$m5*.25 + $f9*$m1*.25 + $f9*$m2*.25 + $f9*$m3*.125 + $f9*$m4*.125 + $f9*$m5*.125 + $m9*$m1*.25 + $m9*$m3*.25 + $m9*$m4*.125,
		$f3*$m4*.0625 + $f3*$m5*.125 + $f3*$m7*.125 + $f3*$m8*.25 + $f4*$m4*.125 + $f4*$m5*.25 + $f4*$m7*.25 + $f4*$m8*.5 + $f6*$m1*.0625 + $f6*$m2*.125 + $f6*$m4*.0625 + $f6*$m5*.125 + $f6*$m7*.0625 + $f6*$m8*.125 + $f7*$m1*.125 + $f7*$m2*.25 + $f7*$m4*.125 + $f7*$m5*.25 + $f7*$m7*.125 + $f7*$m8*.25 + $f9*$m1*.125 + $f9*$m2*.25 + $f9*$m4*.0625 + $f9*$m5*.125 + $m9*$m1*.25 + $m9*$m2*.5 + $m9*$m4*.125 + $m9*$m5*.25,
		$f5*$m3*.125 + $f5*$m4*.0625 + $f5*$m6*.25 + $f5*$m7*.125 + $f6*$m3*.0625 + $f6*$m4*.03125 + $f6*$m6*.125 + $f6*$m7*.0625 + $f8*$m3*.25 + $f8*$m4*.125 + $f8*$m6*.5 + $f8*$m7*.25 + $f9*$m3*.125 + $f9*$m4*.0625 + $f9*$m6*.25 + $f9*$m7*.125,
		$f5*$m4*.0625 + $f5*$m5*.125 + $f5*$m7*.125 + $f5*$m8*.25 + $f6*$m3*.0625 + $f6*$m4*.0625 + $f6*$m5*.0625 + $f6*$m6*.125 + $f6*$m7*.125 + $f6*$m8*.125 + $f7*$m3*.125 + $f7*$m4*.0625 + $f7*$m6*.25 + $f7*$m7*.125 + $f8*$m4*.125 + $f8*$m5*.25 + $f8*$m7*.25 + $f8*$m8*.5 + $f9*$m3*.125 + $f9*$m4*.125 + $f9*$m5*.125 + $f9*$m6*.25 + $f9*$m7*.25 + $f9*$m8*.25 + $m9*$m3*.25 + $m9*$m4*.125 + $m9*$m6*.5 + $m9*$m7*.25,

		$f1*$m1*.5 + $f1*$m2*1 + $f1*$m4*.25 + $f1*$m5*.5 + $f2*$m1*.25 + $f2*$m2*.5 + $f2*$m4*.125 + $f2*$m5*.25 + $f3*$m1*.125 + $f3*$m2*.25 + $f3*$m3*.125 + $f3*$m4*.125 + $f3*$m5*.125 + $f4*$m1*.25 + $f4*$m3*.25 + $f4*$m4*.125 + $f5*$m1*.125 + $f5*$m2*.25 + $f5*$m4*.0625 + $f5*$m5*.125 + $f6*$m1*.125 + $f6*$m2*.125 + $f6*$m3*.0625 + $f6*$m4*.0625 + $f6*$m5*.0625 + $f7*$m1*.125 + $f7*$m3*.125 + $f7*$m4*.0625,
		$f3*$m1*.125 + $f3*$m2*.25 + $f3*$m4*.0625 + $f3*$m5*.125 + $f4*$m1*.25 + $f4*$m2*.5 + $f4*$m4*.125 + $f4*$m5*.25 + $f6*$m1*.0625 + $f6*$m2*.125 + $f6*$m4*.03125 + $f6*$m5*.0625 + $f7*$m1*.125 + $f7*$m2*.25 + $f7*$m4*.0625 + $f7*$m5*.125,
		$f1*$m3*.5 + $f1*$m4*.25 + $f1*$m6*1 + $f1*$m7*.5 + $f2*$m3*.25 + $f2*$m4*.125 + $f2*$m6*.5 + $f2*$m7*.25 + $f3*$m3*.125 + $f3*$m4*.0625 + $f3*$m6*.25 + $f3*$m7*.125 + $f5*$m1*.125 + $f5*$m3*.25 + $f5*$m4*.125 + $f5*$m6*.25 + $f5*$m7*.125 + $f6*$m1*.0625 + $f6*$m3*.125 + $f6*$m4*.0625 + $f6*$m6*.125 + $f6*$m7*.0625 + $f8*$m1*.25 + $f8*$m3*.25 + $f8*$m4*.125 + $f9*$m1*.125 + $f9*$m3*.125 + $f9*$m4*.0625,
		$f1*$m4*.25 + $f1*$m5*.5 + $f1*$m7*.5 + $f1*$m8*1 + $f2*$m4*.125 + $f2*$m5*.25 + $f2*$m7*.25 + $f2*$m8*.5 + $f3*$m3*.125 + $f3*$m4*.125 + $f3*$m5*.125 + $f3*$m6*.25 + $f3*$m7*.25 + $f3*$m8*.25 + $f4*$m3*.25 + $f4*$m4*.125 + $f4*$m6*.5 + $f4*$m7*.25 + $f5*$m1*.125 + $f5*$m2*.25 + $f5*$m4*.125 + $f5*$m5*.25 + $f5*$m7*.125 + $f5*$m8*.25 + $f6*$m1*.125 + $f6*$m2*.125 + $f6*$m3*.125 + $f6*$m4*.125 + $f6*$m5*.125 + $f6*$m6*.125 + $f6*$m7*.125 + $f6*$m8*.125 + $f7*$m1*.125 + $f7*$m3*.25 + $f7*$m4*.125 + $f7*$m6*.25 + $f7*$m7*.125 + $f8*$m1*.25 + $f8*$m2*.5 + $f8*$m4*.125 + $f8*$m5*.25 + $f9*$m1*.25 + $f9*$m2*.25 + $f9*$m3*.125 + $f9*$m4*.125 + $f9*$m5*.125 + $m9*$m1*.25 + $m9*$m3*.25 + $m9*$m4*.125,
		$f3*$m4*.0625 + $f3*$m5*.125 + $f3*$m7*.125 + $f3*$m8*.25 + $f4*$m4*.125 + $f4*$m5*.25 + $f4*$m7*.25 + $f4*$m8*.5 + $f6*$m1*.0625 + $f6*$m2*.125 + $f6*$m4*.0625 + $f6*$m5*.125 + $f6*$m7*.0625 + $f6*$m8*.125 + $f7*$m1*.125 + $f7*$m2*.25 + $f7*$m4*.125 + $f7*$m5*.25 + $f7*$m7*.125 + $f7*$m8*.25 + $f9*$m1*.125 + $f9*$m2*.25 + $f9*$m4*.0625 + $f9*$m5*.125 + $m9*$m1*.25 + $m9*$m2*.5 + $m9*$m4*.125 + $m9*$m5*.25,
		$f5*$m3*.125 + $f5*$m4*.0625 + $f5*$m6*.25 + $f5*$m7*.125 + $f6*$m3*.0625 + $f6*$m4*.03125 + $f6*$m6*.125 + $f6*$m7*.0625 + $f8*$m3*.25 + $f8*$m4*.125 + $f8*$m6*.5 + $f8*$m7*.25 + $f9*$m3*.125 + $f9*$m4*.0625 + $f9*$m6*.25 + $f9*$m7*.125,
		$f5*$m4*.0625 + $f5*$m5*.125 + $f5*$m7*.125 + $f5*$m8*.25 + $f6*$m3*.0625 + $f6*$m4*.0625 + $f6*$m5*.0625 + $f6*$m6*.125 + $f6*$m7*.125 + $f6*$m8*.125 + $f7*$m3*.125 + $f7*$m4*.0625 + $f7*$m6*.25 + $f7*$m7*.125 + $f8*$m4*.125 + $f8*$m5*.25 + $f8*$m7*.25 + $f8*$m8*.5 + $f9*$m3*.125 + $f9*$m4*.125 + $f9*$m5*.125 + $f9*$m6*.25 + $f9*$m7*.25 + $f9*$m8*.25 + $m9*$m3*.25 + $m9*$m4*.125 + $m9*$m6*.5 + $m9*$m7*.25,
		$f6*$m4*.03125 + $f6*$m5*.0625 + $f6*$m7*.0625 + $f6*$m8*.125 + $f7*$m4*.0625 + $f7*$m5*.125 + $f7*$m7*.125 + $f7*$m8*.25 + $f9*$m4*.0625 + $f9*$m5*.125 + $f9*$m7*.125 + $f9*$m8*.25 + $m9*$m4*.125 + $m9*$m5*.25 + $m9*$m7*.25 + $m9*$m8*.5,
		$f6*$m4*.03125 + $f6*$m5*.0625 + $f6*$m7*.0625 + $f6*$m8*.125 + $f7*$m4*.0625 + $f7*$m5*.125 + $f7*$m7*.125 + $f7*$m8*.25 + $f9*$m4*.0625 + $f9*$m5*.125 + $f9*$m7*.125 + $f9*$m8*.25 + $m9*$m4*.125 + $m9*$m5*.25 + $m9*$m7*.25 + $m9*$m8*.5
	);
	
	# Calculate standardized genotype frequencies to sum to 1
	my @after_mating_std;
	for (my $i=0; $i<18; $i++) {
		$after_mating_std[$i] = $after_mating[$i] / sum(@after_mating);
	}

	return @after_mating_std;
}


# Function calculates the allele frequencies 
# given an array of all possible genotypes
sub allele_freqs{
	my @freqs = @_;
	
	my $X_freq = ($freqs[0] + $freqs[1] + $freqs[2] + $freqs[3] + $freqs[9] + $freqs[10]
				+ 0.5*($freqs[4] + $freqs[5] + $freqs[6] + $freqs[11] + $freqs[12] + $freqs[13])) / sum(@freqs);
	my $Y_freq = 1 - $X_freq;

	my $III_freq = ($freqs[0] + $freqs[1] + $freqs[4] + $freqs[7] + $freqs[11] + $freqs[14]
				+ 0.5*($freqs[2] + $freqs[5] + $freqs[8] + $freqs[9] + $freqs[12] + $freqs[15])) / sum(@freqs);
	my $IIIM_freq = 1 - $III_freq;
	
	my $tra_freq = ($freqs[0] + $freqs[9] + $freqs[10] + $freqs[11] + $freqs[12] + $freqs[13] + $freqs[14] + $freqs[15] + $freqs[16]
				+ 0.5*($freqs[1] + $freqs[2] + $freqs[3] + $freqs[4] + $freqs[5] + $freqs[6] + $freqs[7] + $freqs[8] + $freqs[17])) / sum(@freqs);
	my $traD_freq = 1 - $tra_freq;

	return ($X_freq, $Y_freq, $III_freq, $IIIM_freq, $tra_freq, $traD_freq);
}

# Function calculates sex ratio
# given an array of all possible genotypes
sub sex_ratio{
	my @freqs = @_;

	my $female = ($freqs[0] + $freqs[1] + $freqs[2] + $freqs[3] + $freqs[4] + $freqs[5] + $freqs[6] + $freqs[7] + $freqs[8] + $freqs[17])/sum(@freqs);
	my $male = ($freqs[9] + $freqs[10] + $freqs[11] + $freqs[12] + $freqs[13] + $freqs[14] + $freqs[15] + $freqs[16])/sum(@freqs);
	
	return($female, $male);
}
```
